# Genomic diversity and biosynthetic capabilities of sponge-associated chlamydiae

**DOI:** 10.1101/2021.12.21.473556

**Authors:** Jennah E. Dharamshi, Natalia Gaarslev, Karin Steffen, Tom Martin, Detmer Sipkema, Thijs J. G. Ettema

## Abstract

Sponge microbiomes contribute to host health, nutrition, and defense through the production of secondary metabolites. Chlamydiae, a phylum of obligate intracellular bacteria ranging from animal pathogens to endosymbionts of microbial eukaryotes, are frequently found associated with sponges. However, sponge-associated chlamydial diversity has not yet been investigated at the genomic level and host-interactions remain thus far unexplored. Here, we sequenced the microbiomes of three sponge species and found high, though variable, Chlamydiae relative abundances of up to 21.2% of bacterial diversity. Using genome-resolved metagenomics 18 high-quality sponge-associated chlamydial genomes were reconstructed, covering four chlamydial families. Among these, Sorochlamydiaceae shares a common ancestor with Chlamydiaceae animal pathogens, suggesting long-term co-evolution with animals. Sponge-associated chlamydiae genomes mostly resembled environmental chlamydial endosymbionts, but not pathogens, and encoded genes for degrading diverse compounds associated with sponges, such as taurine. Unexpectedly, we identified widespread genetic potential for secondary metabolite biosynthesis across Chlamydiae, which may represent an explored reservoir of novel natural products. This finding suggests that chlamydiae may partake in defensive symbioses and that secondary metabolites play a wider role in mediating intracellular interactions. Furthermore, sponge-associated chlamydiae relatives were found in other marine invertebrates, pointing towards wider impacts of this phylum on marine ecosystems.

## INTRODUCTION

Porifera (*i*.*e*., sponges) are ubiquitous filter-feeding metazoans that provide essential ecosystem services. These animals have complex, deeply integrated, and essential microbiomes that play important roles, such as in global nutrient cycling (1-4). The sponge microbiome also produces secondary (or specialized) metabolites that may contribute to host chemical defence (5). Generally, sponges are known as a source of novel secondary metabolites with medical and industrial relevance (6-9). With increasing exposure to anthropogenic threats, further investigation of the sponge microbiome is essential for understanding host impacts, from acting as detrimental opportunists to providing resilience against dysbiosis and disease (4, 10). Many sponge-associated microbial groups remain uncultured to date and have only recently been explored through cultivation-independent sequencing approaches (11, 12). In a recent large-scale survey, sponge-associated microbial communities were shown to be diverse, yet structured, and composed of taxonomic groups with both generalist and specialist host ranges (13). One of these generalist phyla is Chlamydiae, which is found at high relative abundance in some sponge species (14, 15). Yet, the genomic diversity of sponge-associated chlamydiae has not been previously investigated.

Chlamydiae is a bacterial phylum of obligate eukaryotic endosymbionts well-known for animal pathogens, such as *Chlamydia trachomatis* and other Chlamydiaceae (16-18). Though many chlamydiae instead infect microbial eukaryotes and have more extensive metabolic repertoires (16, 18, 19). Chlamydial environmental distribution and abundance has been underestimated, as their 16S rRNA genes are often missed by primers used to survey microbial diversity (20-22). However, with the recent use of cultivation-independent approaches sequenced chlamydial genomic diversity is quickly expanding, resulting in a widening view of the potential lifestyles of uncultivated chlamydial groups (22-27). Retrieving additional chlamydiae genomes is needed to further our understanding of their ecological impacts, range of host interactions along the parasite-mutualist spectrum, and the evolution of endosymbiosis and pathogenicity (28, 29).

The sponge species *Halichondria panicea, Haliclona oculata*, and *Haliclona xena*, sampled from an estuary in The Netherlands, were previously found to have high chlamydiae relative abundances (15). In the present study, we performed genome-centered analyses of these Chlamydiae in order to gain insight into their sponge-associated lifestyle. Comparative analyses of 18 high-quality sponge-associated Chlamydiae draft genomes revealed degradative capacity also found in other sponge symbionts, and that they share metabolic features with environmental chlamydiae lineages but that are typically absent from known animal pathogens. Unexpectedly, we also identified extensive genetic potential for secondary metabolite biosynthesis across the Chlamydiae phylum. Finally, we found that relatives of these sponge-associated chlamydiae are also associated with additional sponge species and other marine invertebrates, indicating that chlamydiae have important ecological impacts on animals found in marine ecosystems and represent an untapped reservoir for secondary metabolite discovery.

## RESULTS AND DISCUSSION

### Specific chlamydiae lineages vary in relative abundance across three sponge species

Using bacterial-specific small-subunit (SSU) rRNA gene amplicon sequencing high relative abundances of Chlamydiae were found in the sponges *H. panicea* P_S1 (3.9 %), *H. panicea* P_S2 (2.8 %), and *H. oculata* O_S4 (21.2 %) (Figure 1a), which had been collected during the same sampling event as a prior study (15) (Figure S1 and Data S1). However, Chlamydiae relative abundance was substantially lower in three additional sponge specimens, *H. panicea* P_S3 (0.51 %), *H. oculata* O_S5 (0.21 %), and *H. xena* X_S6 (0.26 %) (Figure 1a), that were collected from a similar location, but at different dates (Figure S1 and Data S1). This variation was surprising, as Chlamydiae were previously found to have consistently high relative abundance across all three sponge species (15). Although, Chlamydiae were not detected in other studies of the *H. panicea* microbiome (30). Beyond Chlamydiae, bacterial composition across the sponge species was consistent with a prior investigation, with the same phyla represented and Proteobacteria being the most dominant phylum (Figure 1a) (15). SSU rRNA gene amplicon sequences were subsequently clustered into operational-taxonomic-units (OTUs) at species-level (97% identity) (Data S2), revealing variability in the specific bacterial OTUs present in different sponge specimens (Figure 1b). However, three specific chlamydiae OTUs with high relative abundances (up to 8%; Data S3) were found in all samples with higher overall abundances of Chlamydiae (P_S1, P_S2, and O_S4) (Figure 1a-b). Yet, these OTUs were largely undetected in amplicon data from the other sponge specimens (P_S3, O_S5, and X_S6) (Data S3). In a prior study, these same chlamydial lineages were found at high relative abundance across all specimens of the three sponge species, but were largely undetected in surrounding seawater (15). Together, these results support a close association of these specific chlamydial lineages with the investigated sponge species, while suggesting that any association or increase in abundance is transient.

**Figure 1.**
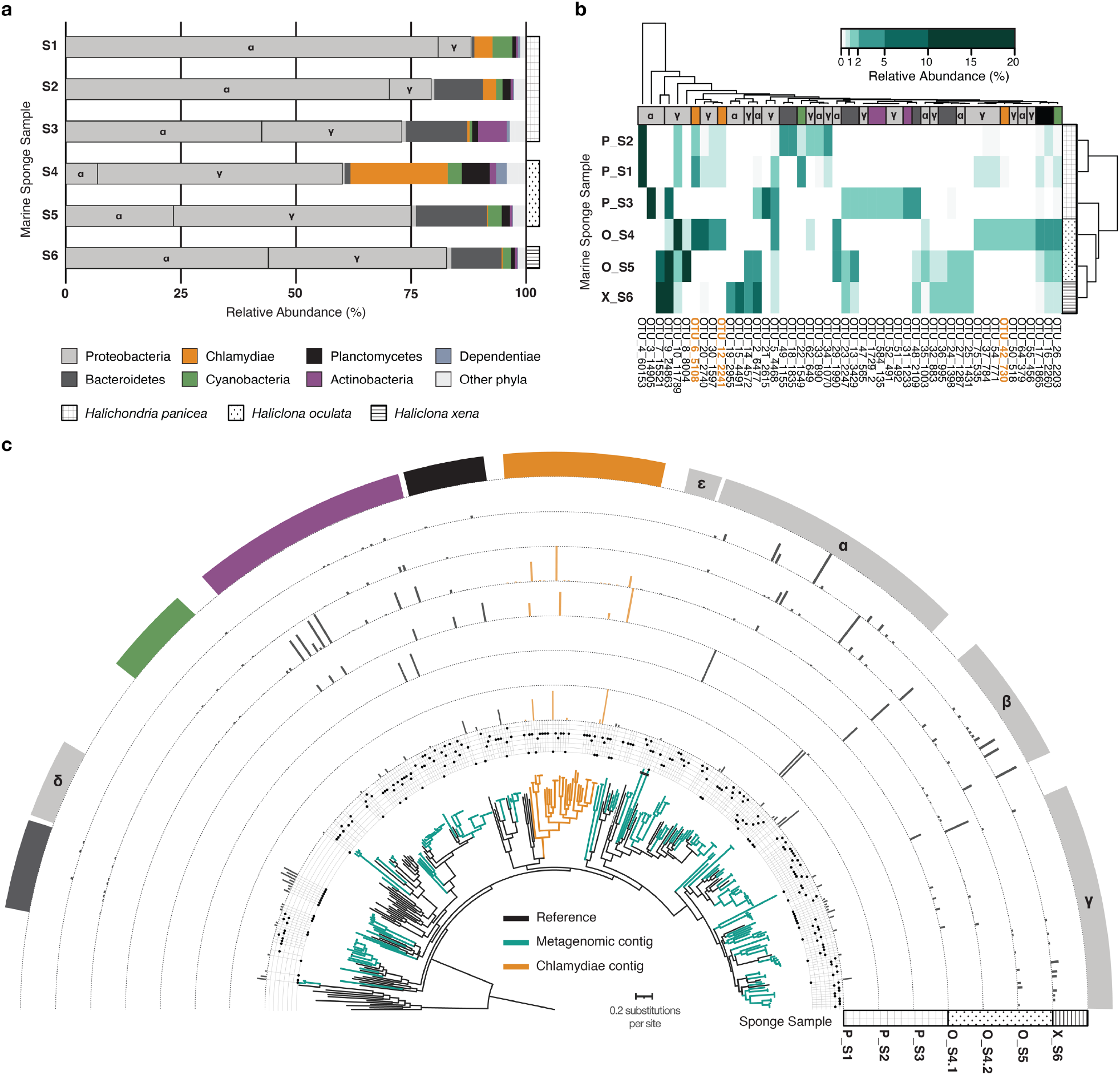
Bacterial SSU rRNA gene amplicon sequencing and metagenomes revealed variably high relative abundances of Chlamydiae across sponge specimens. **a**. Relative abundances of phyla with ≥ 1% relative abundance across samples. **b**. Abundance heat map of OTUs with ≥ 1% relative abundance in a sponge sample. Both OTUs and sponges are hierarchically clustered based on presence patterns. **c**. Concatenated maximum likelihood phylogenetic tree of contigs encoding ribosomal proteins (≥ 5) from across sponge metagenomic assemblies in the context of bacterial reference taxa. The tree is rooted by an archaeal outgroup. The metagenomic origin of each sequence is indicated by the dot plot, with lines corresponding to samples in the following order: P_S1, P_S2, P_S3, O_S4.1, O_S4.2, O_S5, and X_S6. Bars indicate the relative coverage of each ribosomal protein-encoding contig in each metagenome. Phylum affiliation is indicated for each clade according to the colour legend in panel a. In addition, classes are indicated for Proteobacteria: Alphaproteobacteria (α), Betaproteobacteria (β), Deltaproteobacteria (δ), Epsilonbacteria (ε), and Gammaproteobacteria (γ). See legend in panel a for phylum colour assignment and patterns corresponding to each sponge species. See Data S1 for sample information, Data S2 for amplicon OTUs, Data S3 for relative abundances and read counts, Data S4 for metagenomic ribosomal protein contigs and their corresponding coverage, and Data S5 for the uncollapsed ribosomal protein phylogenetic tree.

Chlamydiae have also been detected with varying presence and abundance across a diverse range of other sponge species (13). Despite this, sponge-associated chlamydiae have not been previously investigated at the genome level. With the aim of exploring sponge-associated chlamydial genomic diversity, we sequenced and assembled seven high-quality metagenomes from the six sampled sponge specimens, with two generated for O_S4 (O_S4.1 and O_S4.2) (Figure S1 and Data S1). We assessed microbial diversity in the resulting metagenomes by identifying contigs encoding ribosomal protein gene clusters, and thus representing distinct microbial lineages. Ribosomal protein sequences from each contig were concatenated and a maximum-likelihood (ML) phylogenetic tree reconstructed (Figure 1c and Data S4-S5). Relative abundance of each microbial lineage was inferred by comparing coverage of these contigs from each sponge metagenome (Figure 1c and Data S4). These analyses confirmed the larger patterns in sponge microbial community composition seen in the SSU rRNA gene amplicon results. However, several phyla had lower (i.e., Bacteroides, Cyanobacteria, and Proteobacteria) and higher (i.e., Actinobacteria and Chlamydiae) relative abundances in metagenomes than amplicons, perhaps due to differences in SSU rRNA gene copy number (30). Overall, the metagenomes confirmed the chlamydial patterns seen in amplicon data, with three distinct chlamydiae lineages found in high relative abundance when Chlamydiae were present (P_S1, O_S4.1, and O_S4.2) (Figure 1c and Data S4-S5). This provides further support for transient associations of diverse chlamydiae with several sponge species.

### Genome-resolved metagenomics of sponges expands sequenced chlamydial diversity

Metagenome-assembled genomes (MAGs) were obtained from each sponge metagenome assembly using differential coverage profiles and consensus results from several binning tools (Figure S1). This resulted in 106 medium to high quality MAGs (median 89% completeness and 1.4% redundancy) (Data S4). Chlamydiae MAGs were further collected, manually curated, and reassembled (from P_S1, O_S4.1, and O_S4.2; Figure S1). This resulted in 18 high-quality draft chlamydiae genomes with high contiguity (median 35 contigs), high completeness (median 98.7%), and low redundancy (median 1.01%) (Data S6). Exceptionally, *Chlamydiae* bacterium O_S4.1_1 and O_S1_54 were 99% complete and retrieved on only three contigs each. Phylogenomic trees were then inferred to determine the placement of sponge-associated chlamydiae MAGs within the Chlamydiae phylum, using four subsets of concatenated marker proteins with chlamydial and outgroup species representatives (Figures S2-S3, and Data S6-S7). The obtained species tree topology was consistent across ML reconstructions and the placement of sponge-associated chlamydiae MAGs was highly supported (Figures 2a and S3). A Bayesian phylogenetic tree was also inferred using the smallest subset of marker proteins (n=15), which included those that best resolved the phyla present in the dataset (Figure S2 and Data S7). Here, species topology was overall consistent with the ML trees, with a few exceptions for long-branching taxa.

**Figure 2.**
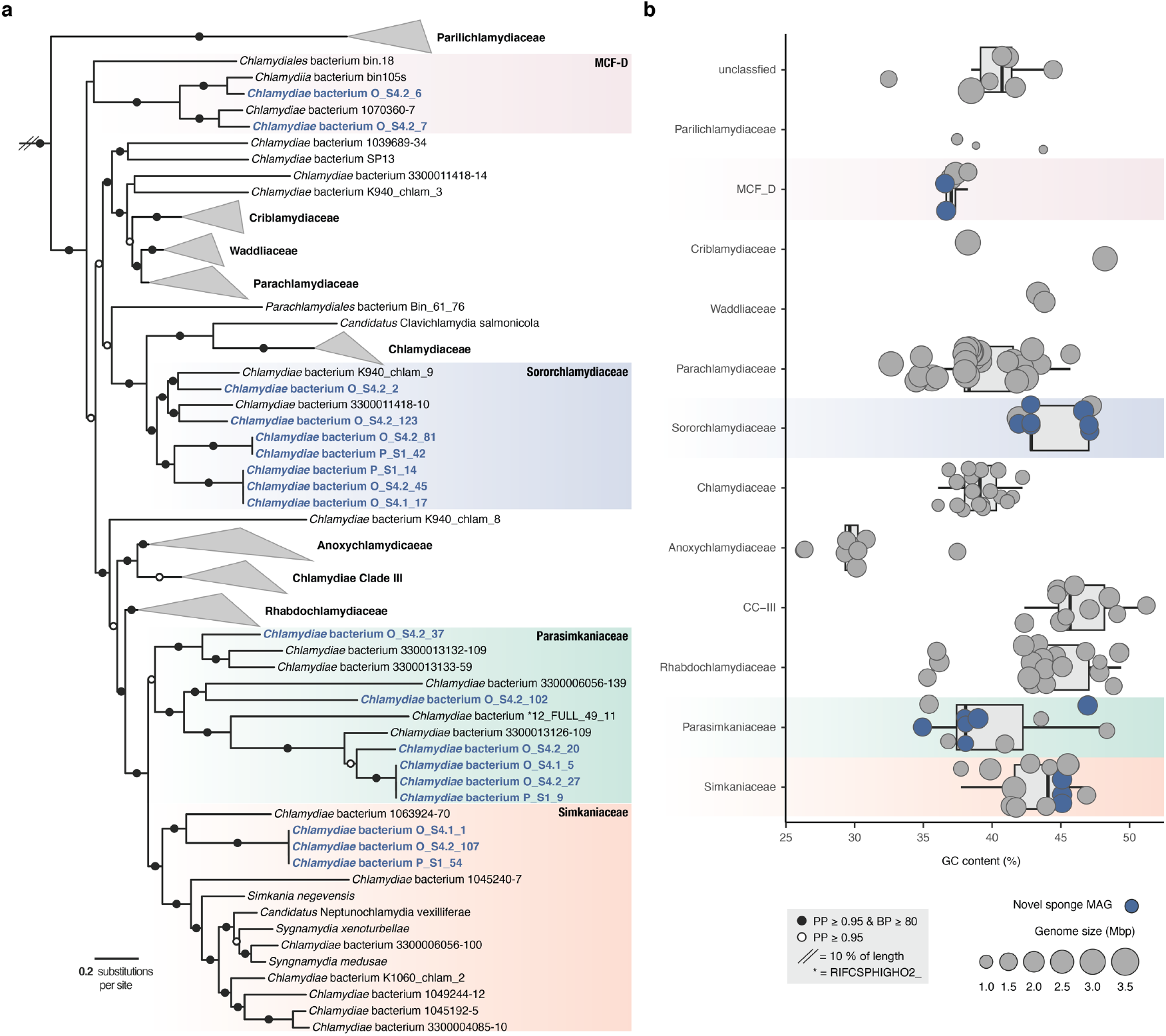
Chlamydiae metagenome-assembled genomes (MAGs) retrieved from sponges are phylogenetically diverse. **a**. Concatenated Bayesian phylogeny of Chlamydiae species relationships inferred using 15 marker gene NOGs (37) under the CAT+GTR+Γ4 model of evolution. Branch support is indicated by coloured circles and includes posterior probability (PP) from the Bayesian inference and non-parametric bootstrap support (BP) from a maximum-likelihood reconstruction of the same dataset inferred with the PMSF approximation (38) of the LG+C60+F+R4 model of evolution (Data S5). Chlamydiae family names are indicated and those not including sponge-associated chlamydiae MAGs collapsed. Sponge-associated chlamydiae MAGs retrieved in this study are highlighted in blue. The tree is rooted by a PVC outgroup (not shown). See Figure S3 and Data S5 for additional species trees and uncollapsed phylogenies. **b**. Genome characteristics of sponge MAGs (blue) in the context of chlamydiae species representatives (grey). Boxplots indicate the distribution of GC content across different chlamydial families, with the area of circles indicating genome size. See Data S6 for genome characteristics.

Sponge-associated chlamydiae MAGs were placed in four distinct chlamydial families (Figures 2a and S3). Interestingly, MAGs with a high level of relatedness were obtained from P_S1, O_S4.1, and O_S4.2, corresponding to the three highly abundant chlamydial lineages identified in amplicon and metagenomic analyses (Figures 1 and 2a). Two MAGs affiliated with the recently described Metagenomic Chlamydial Family D (MCF-D) (26), which includes a MAG from the glass sponge *Vazella pourtalesii* (31), and indicates more widespread association of this family with sponges (Figure 2a). Three additional MAGs and *Chlamydiae* bacterium 1063924-70 (26) formed a well-supported clade with Simkaniaceae from marine, coastal, and host-associated environments (Figures 2a and S3, and Data S6) (26, 32, 33). However, five other MAGs consistently form a group sister to Simkaniaceae with several uncharacterized freshwater chlamydiae (Figures 2a and S3, and Data S6) (26, 34). Based on its well-supported phylogenetic position and conserved gene content, we propose this Simkaniaceae-like sister clade as a new chlamydial family *Candidatus* Parasimkaniaceae (Figures 2 and S4-S5). In ML phylogenies a few long-branching taxa, including one sponge-associated MAG (*Chlamydiae* bacterium O_S4.2_37), clustered together with Simkaniaceae, yet were part of the Parasimkaniaceae with high support in the Bayesian tree (Figures 2a and S3).

The remaining seven sponge-associated chlamydiae MAGs formed a clade with chlamydiae from marine sediments (22) and fungal mycelium (26) that shares a common ancestor with Chlamydiaceae and was previously referred to as Chlamydiae Clade IV (Figures 2a and S3) (22). To reflect its consistent phylogenetic position, we propose to name this family *Candidatus* Sororchlamydiaceae on the basis of improved taxon sampling here, distinct genomic characteristics such as genome size, and conserved gene content (Figures 2b and S4-S5). Identifying sponge-associated Sororchlamydiaceae is also interesting from an evolutionary perspective. Sororchlamydiaceae shares a common ancestor with exclusively animal-associated chlamydiae including *Clavichlamydia salmonicola*, a fish pathogen (35), and the Chlamydiaceae family, that have thus far only been obtained from tetrapods (*e*.*g*., mammals, birds, and reptiles) (36). If many Sororchlamydiaceae are indeed sponge symbionts, this could indicate an ancestral association of these chlamydiae from the Chlamydiales order with metazoa, and subsequent long-term evolution with animal hosts.

The observed differences in the relative abundance of Chlamydiae across the sponge species was unexpected, given that healthy sponges typically have stable microbiomes (4). It was also surprising that three phylogenetically distinct chlamydial lineages were present in these cases. These chlamydiae could be actively acquired from the surrounding water by the filtering action of the sponge. Conversely, in contrast to most other chlamydiae (22, 27), Sororchlamydiaceae genomes also encode flagellar genes (Data S8), which could be used to encounter new hosts. As part of their conserved lifestyles, chlamydiae have both an intracellular dividing phase, and an extracellular non-dividing phase as elementary bodies (19). Chlamydiae can also enter states of persistence inside their host cell when under environmental stress, where they then refrain from dividing (19). As sponge-associated chlamydiae encode genes for the typical chlamydial lifecycle (Data S8; see below), it is possible that they only increase in abundance under specific environmental conditions, and otherwise remain as elementary bodies or in persistence states.

### Similarity in gene content of sponge-associated and environmental chlamydiae

We compared the presence of central metabolic pathways across Chlamydiae and sponge-associated MAGs to see if they resembled previously sequenced groups in terms of core metabolism. Metabolic profiles of sponge-associated chlamydiae did not resemble known animal pathogens (*i*.*e*., Chlamydiaceae and Parilichlamydiaceae), and instead their gene content was similar compared to other members from the same chlamydial family (Figures S4-S5). Generally, patterns also were similar across these chlamydial families, although Sororchlamydiaceae and MCF-D genomes encode more genes for *de novo* biosynthesis of coenzymes and the amino acid asparginine, and most Sororchlamydiaceae for pyrimidines (Figure S5 and Data S9). Overall, our results mirrored prior findings (16, 24), with environmental chlamydiae (*i*.*e*., Amoebachlamydiales) genomes encoding greater metabolic potential, and genomes from the Chlamydiaceae and Parilichlamydiaceae encoding less, consistent with their highly specialized lifestyles as animal pathogens (Figure S5 and Data S9).

Despite generally conserved core metabolism, some pathways are more common in sponge-associated genomes, such as *de novo* biosynthesis of aromatic amino acids (*e*.*g*., tryptophan) in sponge-associated Simkaniaceae (Figure S5 and Data S9). It has been suggested that members of the sponge microbiome exchange aromatic amino acids (39) and sponge-associated Simkaniaceae may likewise do so or provide them to the sponge host. Similarly, most sponge-associated Parasimkaniaceae encode a sodium-transporting NADH dehydrogenase, which is absent in all other members of the family (Figure S5 and Data S9). Conversely, the first three genes of the TCA cycle (*i*.*e*., citrate synthase, aconitase, and isocitrate dehydrogenase) are absent in most sponge-associated Parasimkaniaceae genomes and present in other family members (Data S9). These genes are also absent in Chlamydiaceae, which depend on the uptake of host-derived TCA cycle intermediates (16) and this could likewise be the case for sponge-associated Parasimkaniaceae.

Chlamydial families share a large proportion of gene content, with few genes unique to sponge-associated lineages and more genes shared between closely related lineages (Figure S4). Thus, the sponge-associated chlamydiae accessory genome corresponds more to phylogenetic affiliation than to an ecological association with sponges. This could likewise indicate that sponge-associated chlamydiae have similar lifestyles to close environmental relatives. Supporting this, sponge-associated chlamydiae encode hallmark genes associated with endosymbiosis and the typical chlamydial lifecycle that have been found conserved across Chlamydiae (22, 26). These include nucleotide transporters (NTTs) that can be used for energy parasitism of ATP, the UhpC transporter that can be used to uptake host-derived glucose-6-phosphate, the transcription factor EUO that acts as the master regulator of the chlamydial biphasic lifecycle, and a type III secretion system that can be used to mediate host interactions through the secretion of effectors (16, 18) (Data S9). The presence of these genes strongly suggests that sponge-associated chlamydiae are likewise symbionts with the potential for an endosymbiotic lifestyle within the sponge host. Furthermore, the metabolic profiles of sponge-associated chlamydiae are inconsistent with known chlamydial pathogens of animals, suggesting the potential for host-beneficial interactions.

### degradative capacity indicate sponge-associated chlamydiae engage in sponge interactions

Genes more commonly found in sponge-associated chlamydiae genomes were further investigated and revealed degradative capacities consistent with a sponge-symbiont lifestyle and the use of host-derived compounds (Figures S4-S6, and Data S8-S9). For example, all Sororchlamydiaceae genomes encode taurine dioxygenase (TauD) (Figures S6 and S8). This enzyme degrades taurine to sulfite and is widespread among members of the sponge microbiome who can use it to degrade host-derived taurine (40). Taurine is present in, and released from, nearly all marine metazoans (41). The MAGs *Chlamydiae* bacterium S1_14, S4.1_17, and S4.2_45, which correspond to the abundant Sororchlamydiaceae lineages (Figures 1c and 2a), also encode a putative scyllo-inositol 2-dehydrogenase (IolW), an enzyme that can degrade scyllo-inositols (Figure S6 and Data S8). Similarly, the abundant Simkaniaceae sponge MAGs, *Chlamydiae* bacterium S1_54, S4.1_1, and S4.2_107 (Figures 1c and 2a), have the genetic potential to degrade xylose using the concerted action of xylose isomerase (XylA) and xylulokinase (XylB) (Figure S6 and Data S8). Genes for degrading scyllo-inositol and xylose are rare or absent in other chlamydiae genomes (Figure S6 and Data S8). However, known sponge symbionts do encode these genes as well, such as Poribacteria, a candidate phylum so far only found to reside in the sponge extracellular matrix (42-44). Inositols are common in eukaryotes and rare in bacteria, and the sponge host has been suggested as the source of inositols for inositol-degrading sponge-associated bacteria (43-45).

Several typically-eukaryotic genes were also identified in some sponge-associated chlamydiae genomes and further suggest host-interactions (Figure S6 and Data S8). These genes include sterol reductases identified in Sororchlamydiaceae and sponge-associated Simkaniaceae, and carnitine O-acetyltransferase in sponge-associated Simkaniaceae (Figure S6 and Data S8). Sterol reductases perform the final step in ergosterol or cholesterol biosynthesis and have previously been found in intracellular bacteria, including several chlamydiae (46). In phylogenetic trees of these sterol reductases (K00223/K00213 and K09828), chlamydial sequences branch together with both bacterial and eukaryotic homologs, and could have been acquired by horizontal gene transfer (HGT) from either (Data S5). Carnitine O-acetyltransferase is used in eukaryotes to transport carnitine across the mitochondrial matrix (47), and sponge-associated chlamydiae may use it to obtain carnitine from a sponge host. Carnitine is abundant in animal tissues, and in bacteria it is used as an osmoprotectant or metabolized (47). Genes related to carnitine degradation have been found in members of the sponge microbiome, and carnitine is also abundant in the sponge extracellular matrix (48). In a phylogenetic tree of carnitine O-acetyltransferase (K00624) chlamydial sequences branch within eukaryotic homologs, suggesting that it was obtained by HGT from a eukaryotic host (Data S5).

We also identified genes indicating that several sponge-associated chlamydiae can degrade acetoin, possibly derived from other members of the sponge microbiome, as a carbon and energy source (Figure S6 and Data S8). Acetoin is a volatile organic compound that some bacteria can use as an energy and carbon storage compound, and which can act as a sole carbon source under glucose-limitation (49, 50). Most Sororchlamydiaceae, MCF-D, and sponge-associated Simkaniaceae appear to be acetoin-degrading bacteria as they encode an acetoin dehydrogenase complex (AcoA-B) (Figure S6). These genes are rare in other chlamydiae and largely absent in Parasimkaniaceae (Figure S6). The AcoA-B complex is used to degrade acetoin to acetaldehyde and acetyl-CoA, with the concerted reduction of NAD^+^ to NADH (49-51). Acetaldehyde can then undergo further fermentation to ethanol by alcohol dehydrogenases, such as those found in Sororchlamydiaceae and MCF-D, acetyl-CoA can enter the TCA cycle, and NADH can be shuttled into the electron transport chain or used as a reducing agent in other reactions (Figure S6). Some sponge-associated Simkaniaceae and Sororchlamydiaceae may also be able to produce acetoin through the fermentation of pyruvate to acetolactate using Acetolactate synthase (Figure S6). Acetolactate is then converted to acetoin spontaneously under aerobic conditions or through the action of acetolactate decarboxylase (49), which is encoded in some sponge-associated Simkaniaceae genomes (Figure S6).

In addition, we identified genes suggesting that some sponge-associated chlamydiae are involved in degrading both organic pollutants and toxic compounds produced by members of the sponge microbiome. A putative S-(hydroxymethyl)glutathione dehydrogenase gene was found in several sponge-associated Sororchlamydiaceae and MCF-D MAGs, and in a few other chlamydial groups (Figure S6 and Data S8). This enzyme is involved in oxidizing formaldehyde, a toxic compound that can, for example, be produced as an intermediate during methylotrophy (52), for which genes have been found expressed by sponge symbionts (53). We also identified several genes involved in chlorocatechol and catechol degradation (*i*.*e*., dienelactone hydrolase, 3-oxoadipate enol-lactonase, and catechol 2,3-dioxygenase) (Figure S6 and Data S8). Anthropogenic contamination is a growing concern in marine ecosystems, and microbially-mediated degradation of such organic pollutants and other aromatics reach a branching point at chlorocatechol and catechol (54). Genes involved in these degradative pathways have been found in members of the sponge microbiome, and may also be used to detoxify compounds produced by other microbial members (55). Dienelactone hydrolases perform a key step in the modified *ortho*-cleavage pathway by degrading dienelactones to maleylacetate, which can then be further catabolised before entering the TCA cycle (56, 57). Sponge-associated Simkaniaceae encode dienelactone hydrolase (Figure S6), and genes with the dienelactone hydrolase family protein domain (PF01738) are found across sponge-associated Sororchlamydiaceae genomes (Data S8). These sponge-associated chlamydiae may use these genes to protect themselves, or the sponge host, from these toxic compounds, whether they originate from external sources or other microbial community members.

### Widespread potential for secondary metabolite biosynthesis across the phylum Chlamydiae

Sponges are well-known as reservoirs for the discovery of natural products with medical and industrial importance, many of which are secondary metabolites produced by microbiome members (6-8). Typically, genes for producing secondary metabolites are organized in biosynthetic gene clusters (BGCs) (7). Recently, metagenomic analyses have revealed that BGCs are found across a broad phylogenetic range of bacteria in sponge microbiomes (58). To gain further perspective on the genetic capacity for secondary metabolite biosynthesis across Chlamydiae, we used antiSMASH (59) to investigate BGCs in representative genomes and our sponge-associated chlamydiae MAGs (Figure 3 and Data S10). Unexpectedly, our analysis revealed that BGCs are widespread across the phylum Chlamydiae, with many chlamydial genomes encoding multiple and different types of BGCs, most with unknown functions (Figure 3a and Data S10).

**Figure 3.**
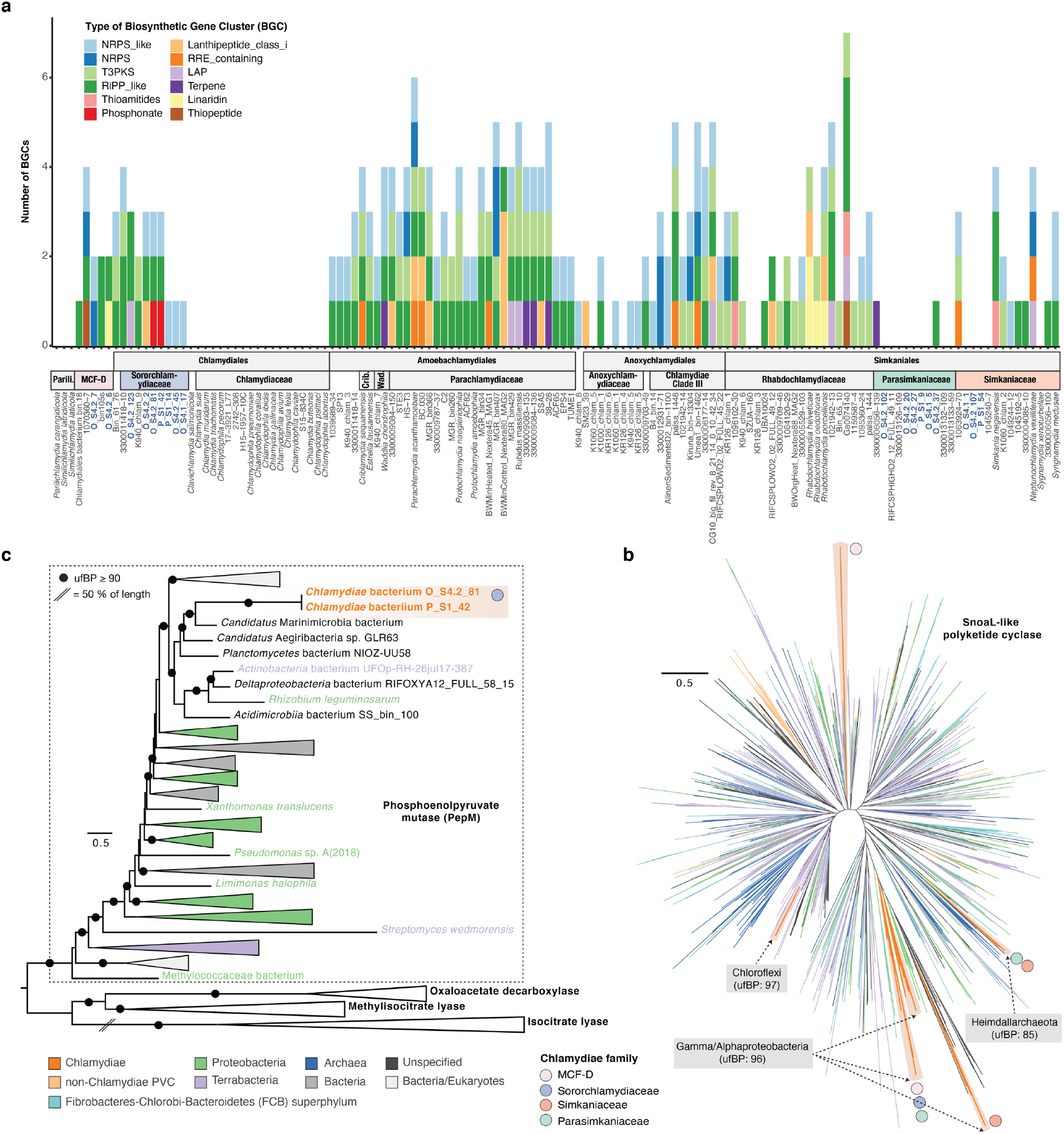
Biosynthetic gene clusters (BGCs) are found across Chlamydiae and sponge-associated chlamydiae genomes. **a**. The number and type of BGCs (in > 1 genome) identified across Chlamydiae using antiSMASH. Order and family classification is indicated alongside shortened species names. See Data S10 for a full overview of BGCs. Maximum-likelihood phylogenies of a group of polyketide cyclases (SnoaL-like; PF07366) (**b**), which are involved in secondary metabolite biosynthesis, and of Phosphophenolpyruvate mutase (PepM; PF13714) (**c**), which performs the first step in phosphonate biosynthesis. Ultrafast bootstrap support values are indicated by a black circle (**c**). Branch and clade colours indicate taxonomy according to the legend (**b, c**). The presence of chlamydiae families including sponge-associated chlamydiae genomes is indicated by the coloured circles, with the taxonomy of the sister clade indicated where supported (**c**). See Data S5 for uncollapsed phylogenies and sequence accessions.

Genomes of Sororchlamydiaceae and Amoebachlamydiales encode the most conserved set of BGCs, with many encoding NRPS-like (non-ribosomal peptide synthetase), RiPP-like (Ribosomally-synthesized and post-translationally modified peptides), and T3PKS (type III polyketide synthase) BGCs (Figure 3a). However, fewer BGCs were identified in Anoxychlamydiaceae, were largely absent in Parasimkaniaceae and sponge-associated Simkaniaceae, and completely absent in all Chlamydiaceae and Parilichlamydiaceae genomes (Figure 3a). We found additional PKS and NRPS genes, which are central in the biosynthesis of various secondary metabolites, in sponge-associated chlamydiae MAGs. MCF-D member *Chlamydiae* bacterium S4.2_7 encodes NRPS gene homologs related to those found in *Simkania negevensis* (Data S8). One group of PKS genes, SnoaL-like polyketide cyclases (PF07366), were found across many sponge-associated chlamydiae MAGs (Data S8). Phylogenetic analysis of this protein family showed that it has been gained multiple times by different chlamydial groups and from diverse potential HGT partners (Figure 3b). Despite BGCs not being widely identified in Parasimkaniaceae, some genomes do encode this PKS gene.

Two Sororchlamydiaceae sponge MAGs (*Chlamydiae* bacterium S1_42 and S4.2_81) encode a phosphonate BGC (Figure 3a), with closest homology to BGCs used to produce the antibiotic fosfomycin (Data S10). Fosfomycin inhibits bacterial cell wall biosynthesis by binding to the active site of MurA, which performs the initial step in peptidoglycan biosynthesis (60). Phosphoenolpyruvate mutase (PepM) performs the first committed step for synthesizing fosfomycin and other phosphonates (61). In a phylogenetic tree, chlamydial PepM homologs formed a well-supported clade together with known PepM sequences, indicated that they likely have the same function (Figure 3c). However, we were unable to determine the HGT donor, since chlamydial homologs formed a clade with diverse bacteria, primarily represented by MAGs (Figure 3c). Interestingly, *Chlamydia* spp. are resistant to very large quantities of fosfomycin due to conserved changes in the MurA active site (62). Further investigation into MurA sequence evolution and the potential for fosfomycin production across chlamydiae could elucidate whether Chlamydiaceae MurA resistance may be connected to an ancestral capacity for fosfomycin production.

Overall, our results show that many chlamydiae have the potential to produce secondary metabolites, which may play a previously unrecognized role in their endosymbiotic lifestyles. As far as we are aware, secondary metabolite biosynthesis had not been previously noted or investigated in chlamydiae. This could be explained by the absence of these gene clusters in the most-studied chlamydial animal pathogens (*i*.*e*., Chlamydiaceae and Parilichlamydiaceae). Chlamydiae is part of the PVC superphylum and other PVC members are also associated with sponges (13, 29). In particular, Planctomycetes have been identified as a potential reservoir of novel secondary metabolites (63, 64). Despite Planctomycetes having substantially larger genomes (64), comparable numbers of BGCs were identified in Chlamydiae genomes (Figure 3a). The Chlamydiae phylum may thus likewise represent a reservoir for the discovery of secondary metabolites. Based on their endosymbiotic lifestyles and smaller genome sizes, it was surprising to identify BGCs as common in many chlamydiae genomes. These chlamydial BGCs could function in inter-microbial warfare, in communication, or in mediating host-interactions (65).

In addition, chlamydial BGCs could function in providing chemical defence to the host. Members of the sponge microbiome have been suggested to provide chemical defence to the sponge host (5). Recently, an endosymbiosis was identified between a *Haliclona* sponge species and a gammaproteobacteria mediated by chemical defence through antibiotic production (66). Sponge-associated chlamydiae that encode BGCs (*i*.*e*., Sororchlamydiaceae and MCF-D) may likewise participate in host-beneficial defensive endosymbioses. MAGs for other typically endosymbiontic bacteria, including Legionellales and Rickettsiales, were also obtained from the metagenomes (Data S4). This suggests the potential for antagonistic intracellular interactions between co-infecting endosymbionts, perhaps mediated by secondary metabolites. Some protist-infecting Parachlamydiaceae have previously been shown to be mutualists that protect their host amoeba against *Legionella* infection through an unknown mechanism (67, 68). This mechanism may have a basis in the production of secondary metabolites as many BGCs were identified in Parachlamydiaceae. Some sponge-associated chlamydiae could offer similar defensive benefits again co-infection to their host. Interestingly, novel antimicrobial compounds have also been isolated from sponges that are active against chlamydial species (69), and could indicate antagonistic interactions with the sponge host or members of the sponge microbiome.

### Chlamydiae are associated with other sponge species and marine invertebrates

To determine whether sponge-associated chlamydiae are associated with other hosts or environments we screened publicly available SSU rRNA gene amplicon datasets for close relatives (≥ 95% identity) (Figure 4a and Data S11). Relatives of sponge-associated chlamydiae were found almost exclusively in marine environments, with higher prevalence in marine invertebrates such as sponges, corals, sea squirts, and molluscs (Figure 4a). MCF-D was present in a higher proportion and wider range of environments, yet still primarily of marine origin. Unfortunately, no SSU rRNA gene was obtained for any of the sponge-associated Parasimkaniaceae MAGs, and their environmental distribution could not be assessed. Despite the clear association with marine environments here, other chlamydiae have also been found associated with sponges from freshwater lakes (70). Relatively few environments were identified with higher abundances of sponge-associated chlamydiae relatives (Data S11), possibly due to chlamydiae being undetected and underestimated by many common primers for surveying microbial diversity (20-22). Those identified included incubations of Great Barrier Reef lagoon water (71), and humic acid amended aquaculture systems (Data S11). In addition to our study, and the previous study of these sponge species (15), Chlamydiae have also previously been found in high relative abundance in association with sponges. This includes *Suberites zeteki*, a marine sponge invasive in Hawaii (14), and *H. panicea*, but only after *ex situ* cultivation (72).

**Figure 4.**
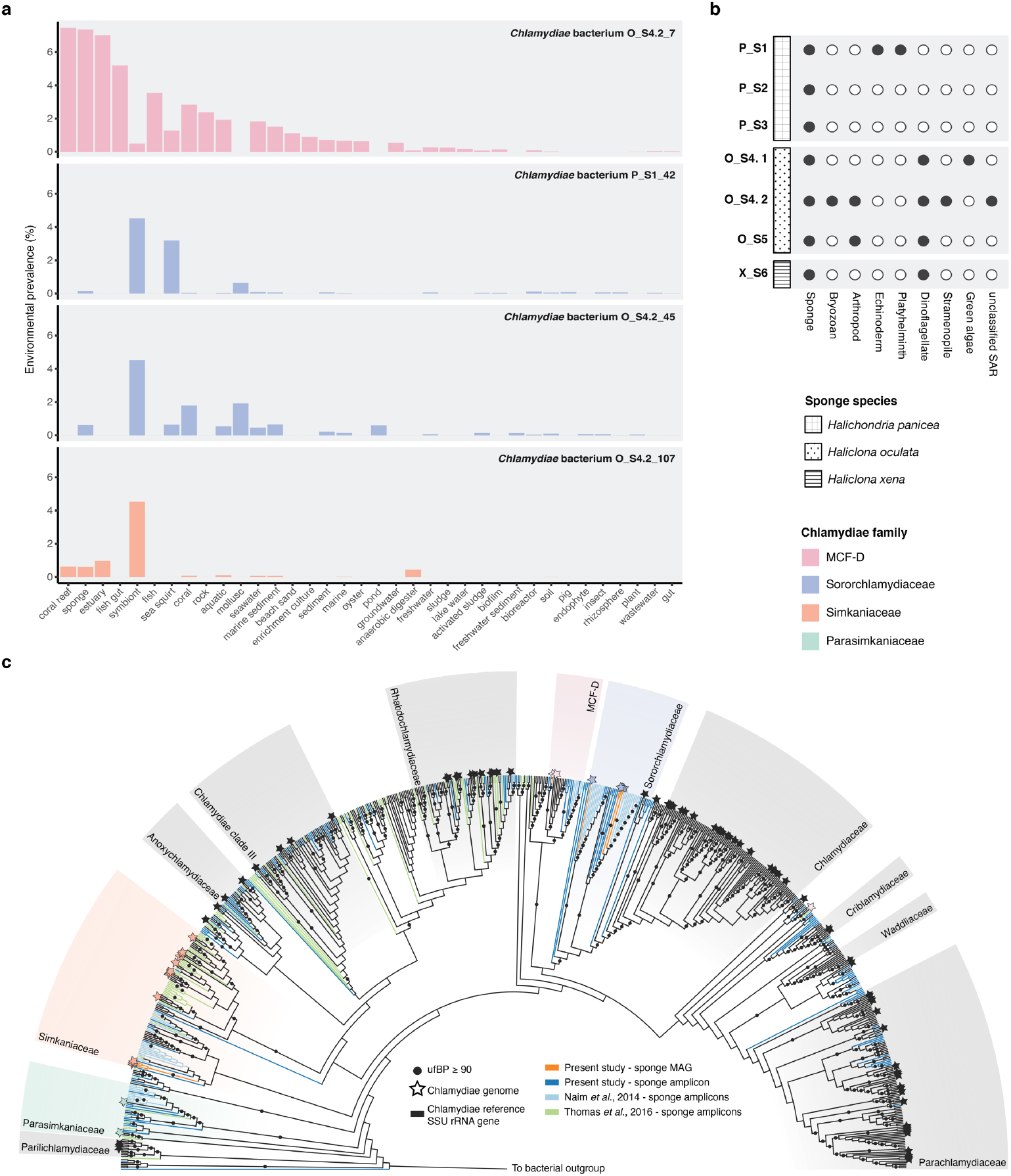
Relatives of sponge-associated chlamydiae MAGs are primarily found in marine habitats, and no other putative eukaryotic hosts were identified in the sponge metagenomes. **a**. Percentage of amplicon samples from various environments with SSU rRNA genes ≥ 95% identity to the indicated sponge MAG. A representative from each phylogroup with MAGs containing SSU rRNA genes is shown. Only environments with ≥ 100 samples, and clear labels are shown. See Data S11 for full IMNGS (73) results of SSU rRNA gene searches against SRA amplicon datasets. **b**. Presence of eukaryotic SSU rRNA genes, and their corresponding taxonomy, across sponge sample metagenome assemblies. See Data S4 for full taxonomy, contig IDs, and contig coverage. **c**. Sponge-associated chlamydiae are restricted to specific chlamydial groups. Maximum-likelihood phylogeny of small subunit rRNA genes from Chlamydiae (and outgroup sequences) inferred using the GTR+F+R10 model and shown as a cladogram for clarity (See Data S5 for sequence accessions and branch lengths.). Included are sequences from reference chlamydiae (black), sponge-associated chlamydiae genomes and amplicons from the present study (orange and blue), chlamydiae amplicons previously obtained from these sponge species (light blue; Naim et al., 2014), and chlamydial amplicons obtained from a prior study of sponge microbial diversity (green; Thomas et al., 2016). Sequences corresponding to chlamydiae genomes, and sequenced diversity, are indicated with stars. Chlamydial families are coloured and labelled.

Although chlamydiae have been found across different sponges and relatives of sponge-associated chlamydiae were detected in various marine invertebrates, it is possible that they instead have another sponge-associated eukaryotic host. To help elucidate this, eukaryotic SSU rRNA genes across the metagenomes were collected and classified (Figure 4b and Data S4). Some of the identified eukaryotes are present in multiple samples. However, importantly, no other eukaryotes apart from the sponge were found across the samples with high chlamydiae relative abundances (Figure 1 and 4b). Still, additional eukaryotes may have been missed in the metagenomes, for example due to DNA extraction bias. We did identify a mitochondrial cytochrome c oxidase subunit 1 (CO1) gene most closely related to the green algae *Picochlorum* in samples with high chlamydiae relative abundances, and this could represent an alternative host (Data S4).

Chlamydiae were found across a wide range of sponge microbiomes in a recent amplicon survey (13). This prompted us to examine whether specific chlamydial groups are associated with sponges. To answer this question we inferred a SSU rRNA gene phylogeny using chlamydial sequences obtained from this wide survey (13), from our study, and from the prior study of these sponge species (15), in the context of a representative dataset of bacterial and chlamydial SSU rRNA gene sequences (20, 21), and sequenced chlamydiae species representatives (Data S6). In the resulting phylogenetic tree, a striking separation became apparent, with the vast majority of sponge-associated sequences affiliating with less studied chlamydial families (Figure 4c). No sponge sequences grouped together with the well-studied Chlamydiaceae animal pathogens, and few sequences affiliated with the protist-infecting Amoebachlamydiales families (*i*.*e*., Criblamydiaceae, Waddliaceae, and Parachlamydiaceae) (Figure 4c). As expected, sequences from the present study and previous study of these sponge species (15) clustered together in Sororchlamydiaceae, MCF-D, Simkaniaceae, and Parasimkaniaceae (Figure 4c). Chlamydial sequences from the wide survey of sponge microbiomes (13) primarily grouped together with Parasimkaniaceae, Simkaniaceae, Rhabdochlamydiaceae, and in unclassified groups (Figure 4c). Altogether, these observations provide evidence for widespread associations between less-studied chlamydial groups and sponges.

### Conclusions and future perspectives

Using genome-resolved metagenomics we have expanded sequenced chlamydiae with 18 high-quality genomes that provide insight into chlamydial associations with animal hosts. All cultivated chlamydiae are obligately intracellular (18) and based on their gene content sponge-associated chlamydiae are likely endosymbionts. However, direct confirmation is needed. Given that sponge-associated chlamydiae are capable of acquiring carbon and energy directly from eukaryotic hosts (*e*.*g*., through the action of NTTs and UhpC *etc*.), it is surprising that they have the capacity to degrade a wide range of sponge-derived compounds and compounds present in the larger microbial community. The presence of genes for degrading toxins and pollutants, and BGCs in some sponge-associated chlamydiae could suggest host-beneficial effects. More generally, our findings open the door for further exploration of BGCs in Chlamydiae, and suggest larger roles for secondary metabolites in endosymbiotic interactions. Relatives of sponge-associated chlamydiae are prevalent in other sponges and marine invertebrates, where they have unknown effects.

We found that sponge-associated chlamydiae vary in their relative abundance across specimens of the same sponge species. This points to important unanswered questions about the nature of sponge-chlamydiae associations. Foremost, it is currently unclear whether the presence of chlamydiae is beneficial or detrimental to the sponge, and if their presence potentially represents an indicator for environmental conditions and host health? Future studies are also needed to confirm whether chlamydiae associate directly with the sponge host as symbionts, or if their interaction is secondary through another eukaryotic host. Altogether, our work represents a first step in untangling the potentially wide impacts of chlamydiae on marine ecosystems.

### Description of *Candidatus* Sororchlamydiaceae fam. nov

(So.ror.chla.my.di.a.ce’ae. L. fem. n. *soror* sister; *Chlamydiaceae* taxonomic name of a bacterial family; L. suff. *-aceae* ending to denote a family; *Sororchlamydiaceae* referring to the close relationship to the bacterial family *Chlamydiaceae*)

The family *Candidatus* Sororchlamydiaceae (formally Chlamydiae Clade IV) represents a distinct monophyletic lineage, sister to Chlamydiaceae, supported by concatenated marker protein phylogenies in the present study (Figures 2a and S3) and in prior work (22, 26). Sororchlamydiaceae share conserved gene content and metabolism, and have larger genome sizes and greater metabolic potential than Chlamydiaceae members (Figures 2b and S4-S6) (22).

### Description of *Candidatus* Parasimkaniaceae fam. nov

(Par.a.sim.ka.ni. a.ce’ae. Gr. prep. *para* alike, alongside of; N.L. fem. n. *Simkania* taxonomic name of a bacterial genus; L. suff. *-aceae* ending to denote a family; *Parasimkaniaceae* referring to the close relationship to the bacterial family *Simkaniaceae*)

The family Candidatus Parasimkaniaceae represents a distinct monophyletic lineage supported by concatenated marker protein phylogenies that is closely related to the Simkaniaceae family (Figures 2a and S3). Parasimkaniaceae share conserved gene content, and have on average smaller genome sizes and reduced metabolic potential relative to Simkaniaceae members (Figures 2b and S4-S6).

## MATERIALS AND METHODS

### Sponge collection and DNA extraction

Six sponge specimens from three sponge species (*Halichondria panicea –* P_S1, P_S2, and P_S3; *Haliclona oculata –* O_S4 and O_S5; *Haliclona xena* – X_S6) were collected from the Oosterchelde estuary in the Netherlands, washed in autoclaved seawater, and stored at -80 °C (Figure S1 and Data S1). Several were sampled alongside those investigated in a previous study (15). The sponge SSU rRNA gene from each metagenome was used to confirm sponge species identification (Data S1).

For generating SSU rRNA gene amplicons, DNA was extracted from 0.2 g of each specimen using the DNAeasy PowerLyser Powersoil Kit according to manufacturer’s protocols, with DNA elution in water and bead beating with the PowerLyzer 24 homogenizer at 4000 rpm for 45 s. For O_S4 (O_S4.2), O_S5, and X_S5 this DNA was also used for metagenomic sequencing (Figure S1). However, due to high levels of fragmentation an additional extraction optimized for high-molecular-weight DNA was performed for metagenomic sequencing of P_S1, P_S2, P_S3 and O_S4 (O_S4.1) (Figure S1). Here, bead beating with the DNAeasy PowerLyser Powersoil Kit was used as described above, but with the addition of 0.2 M ethylenediamine tetraacetic acid (EDTA) (1:1 ratio) prior to the lysis step to inhibit DNAses. After bead beating, 10% cetyl trimethylammonium bromide buffer (CTAB) (1:4 ratio), 0.5 M NaCl (1:2 ratio), 0.1 M EDTA (1:4 ratio), 10 µL β-Mercaptoethanol (100%), and 5 µL Proteinase K (600 mAU/mL) were added and samples incubated overnight at 56 ºC. RNAse A was then added (final concentration of 0.3 ng/µl) and the sample incubated for 30 min at 37 ºC. Two rounds of chloroform/Isoamylalcohol 24:1 (1:1 ratio) addition, incubation for 2 min at room temperature, centrifugation (10000 x g for 10 min), and transfer of the aqueous phase were then performed. DNA was precipitated using isopropanol (6 hr) and pelleted with centrifugation for 15 min at 10000 x g at 4 ºC, before being washed twice with 80% ethanol and eluted in water.

### Generation of bacterial SSU rRNA gene amplicons

A two-step PCR approach was used to obtain SSU rRNA gene fragments for amplicon sequencing (Figure S1), using the bacterial-specific primers S-D-0564-a-S-15 (AYTGGGYDTAAAGNG) and S-D-Bact-1061-a-A-17 (CRRCACGAGCTGACGAC) (74), that capture most chlamydial lineages (22). HotStarTaq DNA Polymerase (QIAGEN) was used with the following reaction conditions: initial heat activation at 95 °C (15 min), followed by 28 cycles of denaturation at 94 °C (60 s), a step-down to 70 °C (1 s), a ramping rate of 0.4 °C/s to 50 °C for annealing (60 s), and a ramping rate of 0.8 °C/s to 72 °C for extension (60 s), with a final extension at 72 °C (10 min). A second PCR reaction was performed according to the manufacturers protocol to obtain sequence libraries with adaptor sequences from the TruSeq DNA LT Sample Prep Kit (Illumina). PCR products were purified using magnetic Agencourt AMPure XP beads (Beckman Coulter) and sequencing performed on the Illumina MiSeq platform (2×300 bp).

Sequence reads were demultiplexing and quality control performed using cutadapt v. 1.10 (75) to remove remaining adaptor and primer sequences, trim 3’ read ends to a minimum Phred quality score of 10, and remove reads shorter than 100 bp in length. VSEARCH v. 1.11.1 (76) was then used to merge forward and reverse reads (–fastq-minovlen 16), to de-replicate reads (– derep_fulllength), and to obtain centroid OTU clusters at 97% identity. Chimeric sequences were detected and removed using UCHIME (77) with the SILVA123.1_SSUref_tax:99 database (78). OTUs were taxonomically classified using the LCAClassifier from CREST-2.0.5 (79) (Data S2). OTU relative abundance across samples are available in Data S3.

### Metagenome sequencing and assembly

The Nextera DNA Library Prep Kit (Illumina) was used to prepare sequence libraries with 25 ng of input DNA, followed by sequencing with Illumina NovaSeq6000 System for P_S1, P_S2, P_S3 and O_S4 (O_S4.1), and with Ilumina HiSeq2500 System for O_S4 (O_S4.2), O_S5 and X_S6. Quality control of resulting sequence reads was performed to remove adaptors and low-quality sequences using Trimmomatic v. 0.35 (80) with the options: ILLUMINACLIP:TruSeq3-PE.fa:2:30:10 LEADING:3 TRAILING:3 SLIDINGDOWN:4:15 MINLEN:50. Read quality was assessed using FastQC v0.11.4 (81). Resulting paired sequence reads were then assembled using MEGAHIT v3.13 (82) (--meta --only-assembler), and assembly statistics obtained with QUAST v5.0.2 (83) (Data S1). Proteins sequences were predicted using prodigal v2.6.3 (84). Barrnap v. 0.9 (85) was used to identify metagenomic SSU rRNA genes, which were classified using the LCAClassifier from CREST-3.1.0 (79) (Data S4).

### Binning of metagenome-assembled genomes

Differential read coverage was obtained by mapping each set of sequence reads against each assembled metagenome using Bowtie2 v2.2.6 (86). MAGs from each metagenome were obtained using differential coverage binning with metaBAT 2.12.1 (87), CONCOCT v. 1.1.0 (88), and MaxBin v. 2.2.7 (89) (Figure S1). The metaWRAP v. 1.2.4 “bin_refinement” module was then used to consolidate resulting bins into hybridized bin sets. The highest quality hybridized or original bin was selected with a cut-off of 70% completeness and 10% redundancy (90) (Data S4). Several assemblies had smaller sizes (P1_S2 and P1_S3) and few MAGs above quality thresholds were obtained for these (Data S1 and S4). Further manual refinement of chlamydiae MAGs was performed using anvi’o v.6.2 (91, 92), followed by reassembly with the metaWRAP v. 1.2.4 “reassemble_bins” module (90), and additional manual curation (Figure S1).

### Genome characteristics and annotation

Chlamydiae MAGs were annotated using Prokka v1.14.6 (93). In addition, protein-coding genes were annotated with NCBI NR protein database (94) top blastp hits, and Pfam (95) and TIGRFAM (96) domains identified by InterProScan 5.47-82.0 (97). Comparative genomic analyses were performed between chlamydiae MAGs obtained here and a set chlamydiae species representatives with high-quality genomes. Genome characteristics and genome quality were determined with MiComplete v. 1.1.1 (98) (Data S6) using a marker gene-set conserved in complete chlamydiae genomes (22) (Data S6). Genes were also assigned to eggNOG 4.5 (37) NOGs at the universal-level using eggNOG-mapper 1.0.3 (“-d NOG”) (99), and KEGG KOs (100) identified using GhostKOALA (101) (chlamydiae representatives) and BlastKOALA (chlamydiae from the present study). Identified COGs were also assigned to COG pathways (Data S8) (102). AntiSMASH 6 beta (59) was used to identify BGCs, and top hits to MIBiG clusters (103) where found (Data S10).

### Ribosomal protein phylogeny of metagenomic contigs

Microbial community composition was assessed by identifying metagenomic contigs encoding ribosomal proteins (at least 5 of 15), typically found together in a conserved gene cluster (104), using a previously described pipeline (105) (Data S4). Ribosomal proteins from each contig were concatenated and a ML phylogeny inferred using RAxML 8.2.4 (106) with the PROTCATLG model of evolution, and 100 rapid bootstrap replicates (Data S5). Read coverages of these metagenomic contigs were compared to measure relative abundances (Data S4).

### Chlamydiae species phylogeny

Protein sequences from 74 single-copy marker genes (Data S7), found conserved across chlamydiae MAGs and PVC species representatives (Data S6), were each aligned using MAFFT-L-INS-i v7.471 (107), and trimmed with BMGE (108) (entropy of 0.6). IQTREE v. 1.6.12 (109) was used to infer single-gene trees using ModelFinder (110) for model selection from LG (111) and LG empirical mixture models (C10 to C60) (112), with gamma or free-distributed rates (+G or +R) (113), and with or without empirically determined amino acid frequencies (+F). Trees were manually inspected, divergent sequences removed, and the process repeated where necessary (Data S7). To determine which marker genes had the strongest phylogenetic signal, the monophyly of PVC phyla in the single-gene trees was assessed (Data S7). Four marker gene datasets were chosen to include all genes (74 NOGs), genes where Chlamydiae was monophyletic (60 NOGs), genes where Chlamydiae and most other PVC phyla were monophyletic (40 NOGs), and genes all PVC phyla were monophyletic (15 NOGs) (Data S7). ML species phylogenies were then inferred as described above for concatenated alignments of each dataset.

The topology across all species trees was consistent and the placement of sponge-associated chlamydiae MAGs supported (Figure S3 and Data S5). Further analyses were thus performed with only the smallest concatenated dataset (15 NOGs). A ML tree was inferred using IQTREE v. 1.6.12 (109) with 100 non-parametric bootstraps under the PMSF approximation (38) of the LG+C60+F+R4 model of evolution (Figure 2a). A Bayesian phylogeny was also inferred using PhyloBayes-MPI 1.7b (114) and the CAT+GTR+Γ4 model of evolution (115), with four independent MCMC chains. After approximately 100,000 generations the four chains had not converged (with a burn-in of 5000, and sampling every 10 generations) (Data S5). However, two chains had converged (maxdiff of 0.16) (Figure 2a), and the topology of both deeper nodes and the placement of sponge-associated chlamydiae was consistent across all chains (Data S5).

### Single-protein phylogenies of genes of interest

The eukaryotic affiliation of typically-eukaryotic genes was confirmed using a phylogenetic workflow (https://github.com/jennahd/HGT_trees) (Data S5). Additionally, phylogenetic trees were inferred for protein sequences with SnoaL-like PKS (PF07366) and PepM (PF13714) protein domains (Data S5). Here, DIAMOND blastp (116) searches (with “max-target-seqs 2000” and “more-sensitive”) were performed against NCBI’s NR database (117), and sequence redundancy removed using CD-HIT v. 4.8.1 (118) at 80% identity. For PepM, the top 100 hits to the Swiss-Prot database, of proteins with curated annotations, were additionally retrieved (119). Sequences were aligned and trimmed as above. For SnoaL-like PKS an initial tree was inferred using FastTree 2 (120) and a subset of more distantly-related sequences removed. ML phylogenies was then inferred using IQTREE v. 1.6.12 (109), with model selection by ModelFinder (110) as described above. The LG+C60+F+R4 model was selected in both cases, and 1000 ultrafast bootstraps inferred (121) (Data S5).

### Small subunit rRNA gene phylogeny of sponge-associated Chlamydiae diversity

Chlamydial SSU rRNA gene amplicon OTUs were collected from the present study, the prior study of these marine sponge species (15), and from a survey of sponge microbial diversity (13). These were combined with full-length and near full-length SSU rRNA genes from chlamydiae MAGs and reference chlamydiae genomes (Data S6), and prior surveys of chlamydial (20). Sequences were added to a previously published bacterial SSU rRNA gene alignment of 85% sequence identity representatives (21) using MAFFT-L-INS-i v7.471 (107) (“--add” for near full-length sequences, and “--addfragments” for amplicon OTUs). The alignment was trimmed using trimAl v1.4.rev15 (122) with a gap threshold of 0.1. A ML phylogeny was then inferred using IQTREE v. 1.6.12 (109), with the GTR+F+R10 model selected from GTR models (123) by ModelFinder (110) and 1000 ultrafast bootstraps (121).

### Environmental distribution

The environmental distribution of specific chlamydial lineages was assessed using IMNGS (73). Here, environmental samples were screened for sequences with at least 95% identity to SSU rRNA genes from several sponge-associated chlamydiae MAGs (Data S11). Samples with at least 0.1% relative abundance of these chlamydial lineages were also identified (Data S11).

### Data visualization and availability

Phylogenetic trees were visualized using the ETE3 Toolkit (124), iTOL (125), and Figtree v1.4.4 (http://tree.bio.ed.ac.uk/software/figtree/). Plots were generated using R version 4.0.3 (R Core Team, 2020), with ggplot2 v. 3.3.2 (126), and with UpSetR v. 1.4.0 (127) for intersection plots.

Assembled metagenomes, metagenome sequence reads, amplicon sequence reads, and MAGs generated from each sponge sample can be found deposited under BioProject PRJNA504765. Accessions for data analyzed or obtained in this study can be found in Data S1, S4, and S6. Whole Genome Shotgun projects for sponge metagenome assemblies P_S1, P_S2, P_S3, O_S4.1, O_S4.2, O_S5, and X_S6 have been deposited at DDBJ/ENA/GenBank under the accessions JAHZIM000000000, JAHZIN000000000, JAHZIO000000000, JAHZIP000000000, JAHZIS000000000, JAHZIQ000000000, and JAHZIR000000000, respectively. The versions described in this paper are versions JAHZIM010000000, JAHZIN010000000, JAHZIO010000000, JAHZIP010000000, JAHZIS010000000, JAHZIQ010000000, and JAHZIR010000000. Sequence alignments and phylogenetic trees in newick format are provided at the Figshare data repository DOI: 10.6084/m9.figshare.14939475.

## ACKNOWLEDGEMENTS

We thank Klaske Schippers and Rene Wijffels for obtaining the sponge samples examined in this study. We thank D. Tamarit for useful advice and discussions. Sequencing was performed at the Uppsala Genome Centre, a sequencing platform that is hosted at Uppsala University as part of the National Genomics Infrastructure at the Science for Life Laboratory. This national infrastructure is supported by the Swedish Research Council (VR-RFI) and the Knut and Alice Wallenberg Foundation. Computational resources were provided by the Swedish National Infrastructure for Computing (SNIC) at UPPMAX with the project number SNIC 2020/15-158, PDC with the project numbers SNIC 2019/3-474 and SNIC 2020/5-473, and HPC2N with the project number SNIC 2019/5-114. This work was supported by grants from the European Research Council (ERC Starting and Consolidator grants 310039 and 817834, respectively), and the Swedish Research Council (VR grant 2015-04959).

## SUPPLEMENTARY FIGURES

**Figure S1.**
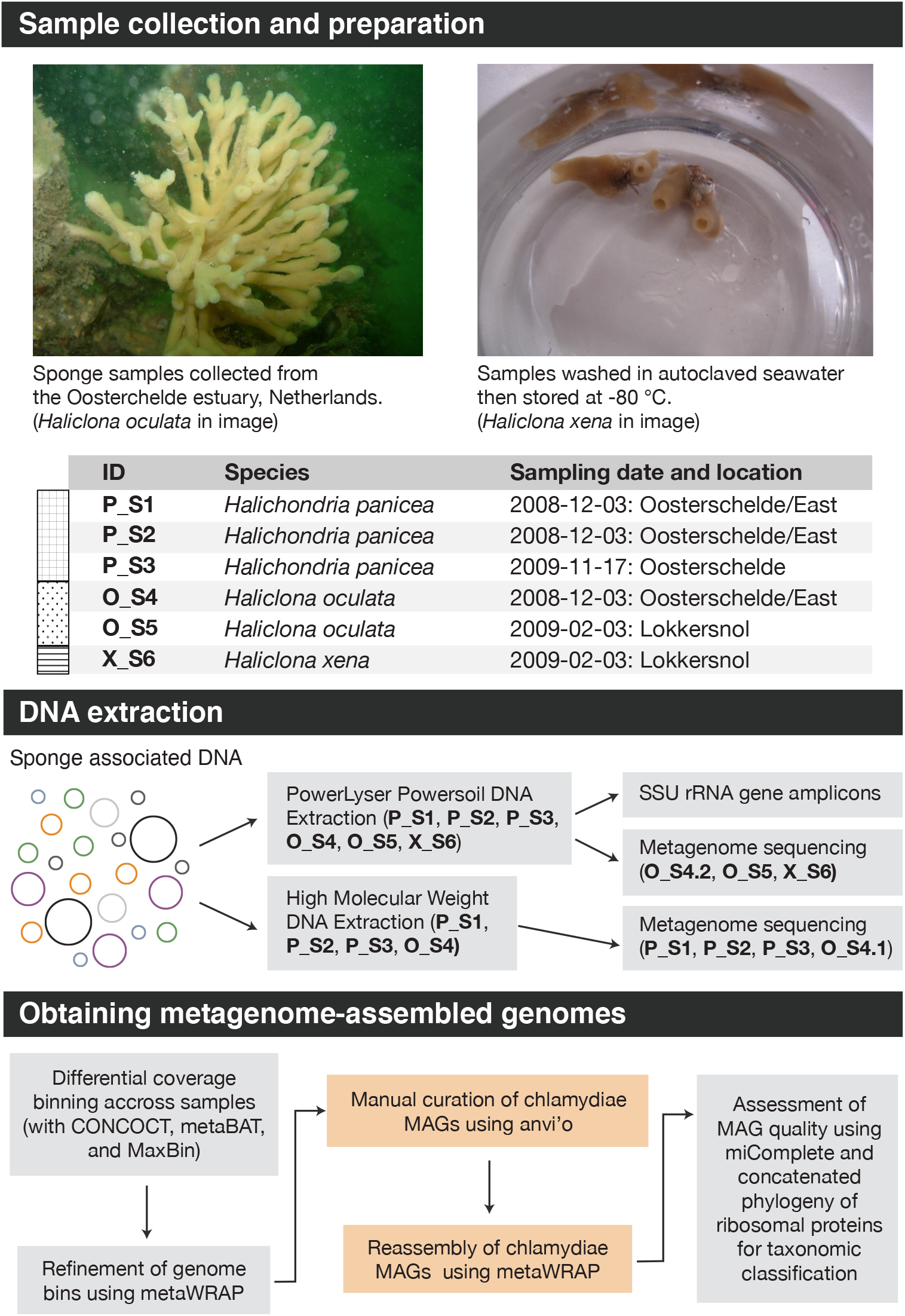
Methods overview of sample collection, sponge specimens collected, DNA extraction methods, amplicon sequencing, metagenome sequencing and assembly, and MAG binning and refinement.

**Figure S2.**
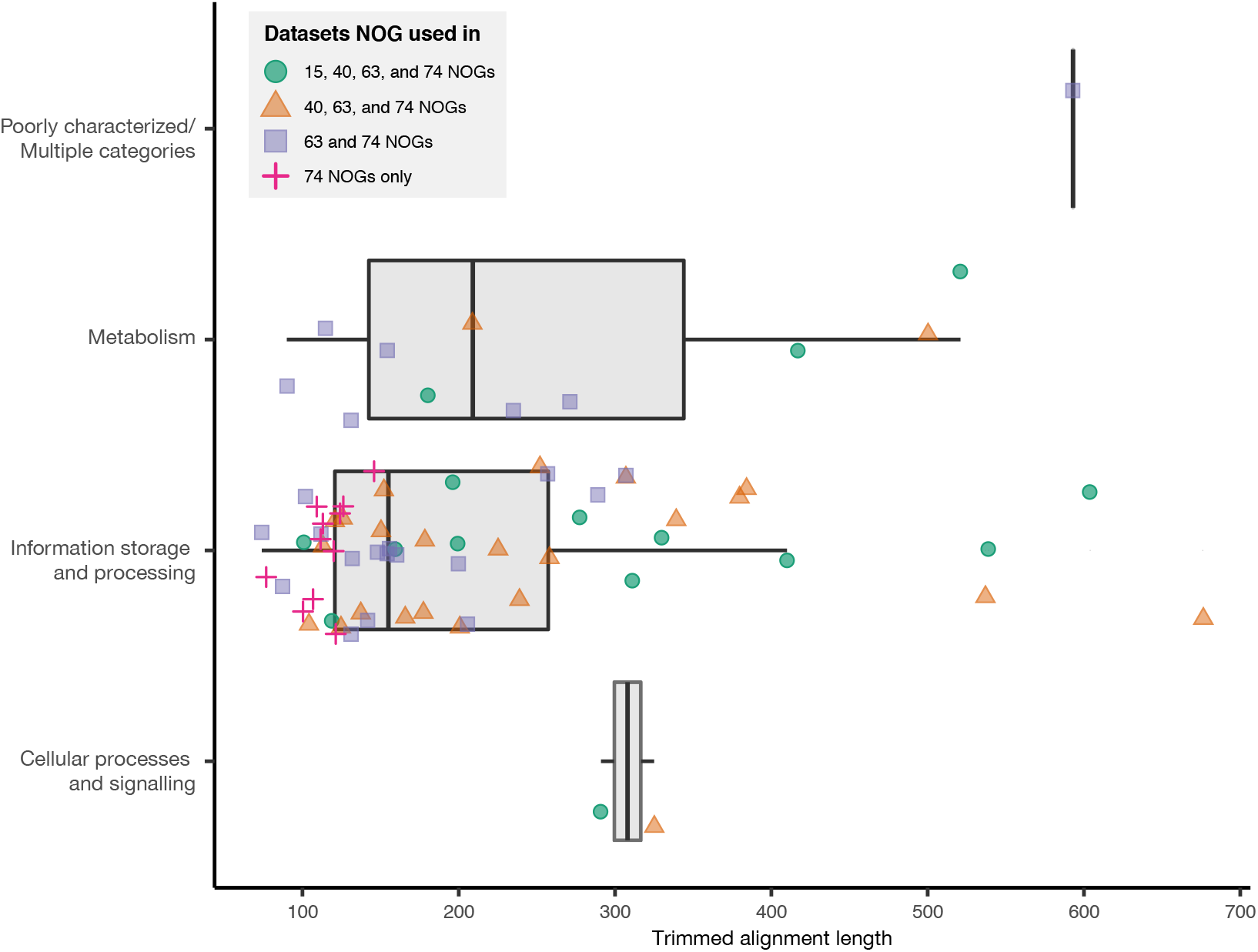
COG category membership of the 74 marker gene NOGs included in Chlamydiae species tree concatenated datasets alongside their trimmed alignment lengths. NOGs were assigned to datasets based on phylogenetic signal, measured by resolving Chlamydiae and other PVC phyla as monophyletic in single-protein trees. All marker gene NOGs were included in the 74 NOG dataset, those resolving Chlamydiae as monophyletic in the 63 NOG dataset, those resolving Chlamydiae and most PVC phyla as monophyletic in the 40 NOG dataset, and those resolving all PVC phyla as monophyletic in the 15 NOG dataset. See Data S7 for tree refinement and selection details.

**Figure S3.**
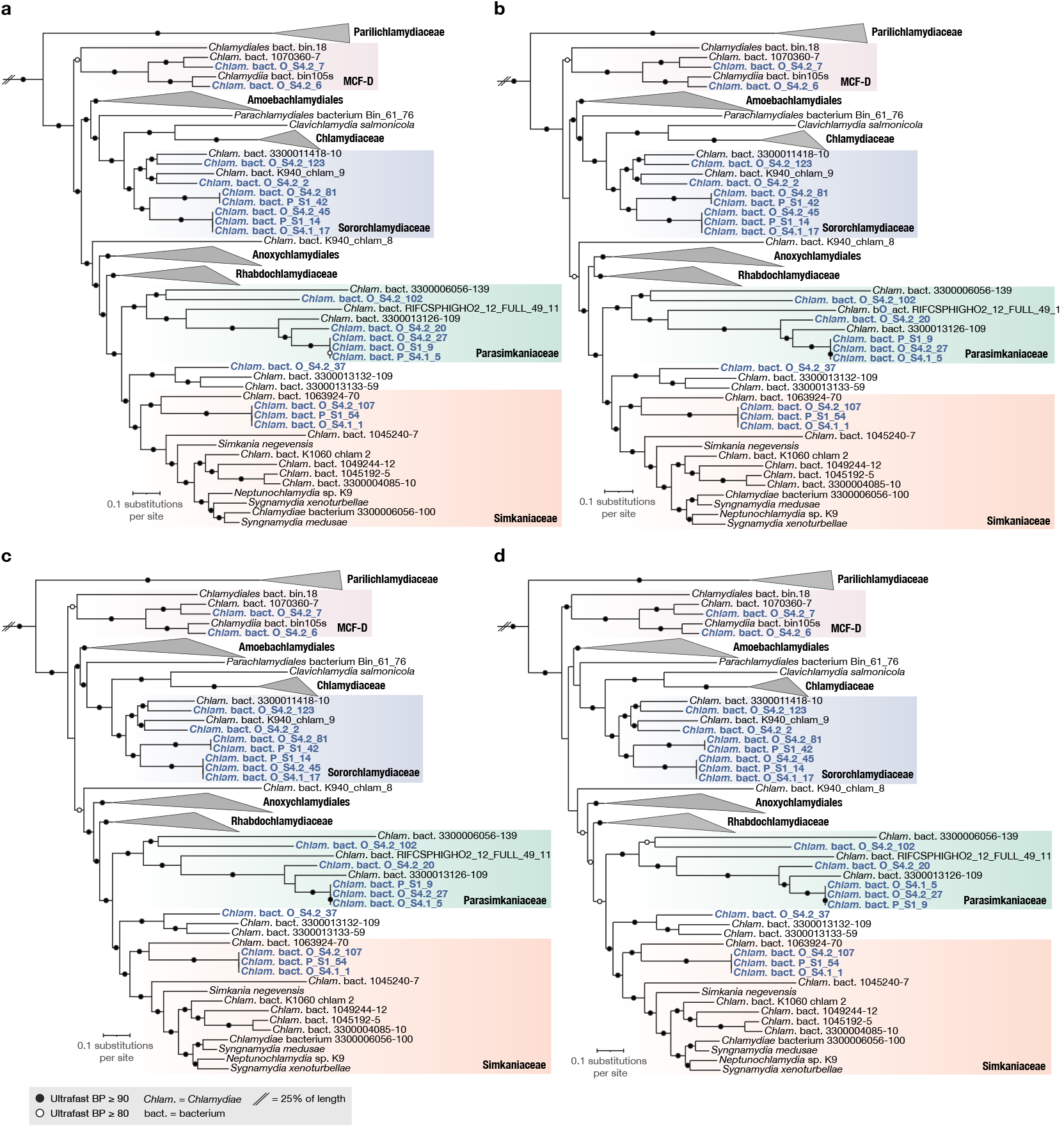
The topology of Chlamydiae species relationships is consistent across reconstructions using subsets of the larger initial dataset of 74 NOGs. Concatenated maximum-likelihood protein phylogenies of Chlamydiae species inferred under the LG+C60+F+R4 model of evolution with 74 (**a**), 63 (**b**), 40 (**c**), and 15 (**d**) single-copy marker NOGs with 16760, 15502, 11211, and 4757 amino acid alignment positions, respectively. Trees are rooted by a PVC outgroup (not shown), with this branch reduced 25 % in length. Sponge chlamydiae MAGs are coloured in blue, with relevant families coloured, and phylogroups with high relative abundance highlighted. Branch support is indicated by circles according to the legend. See Data S5 for uncollapsed species trees and Data S6 for NOGs included in each phylogenetic dataset.

**Figure S4.**
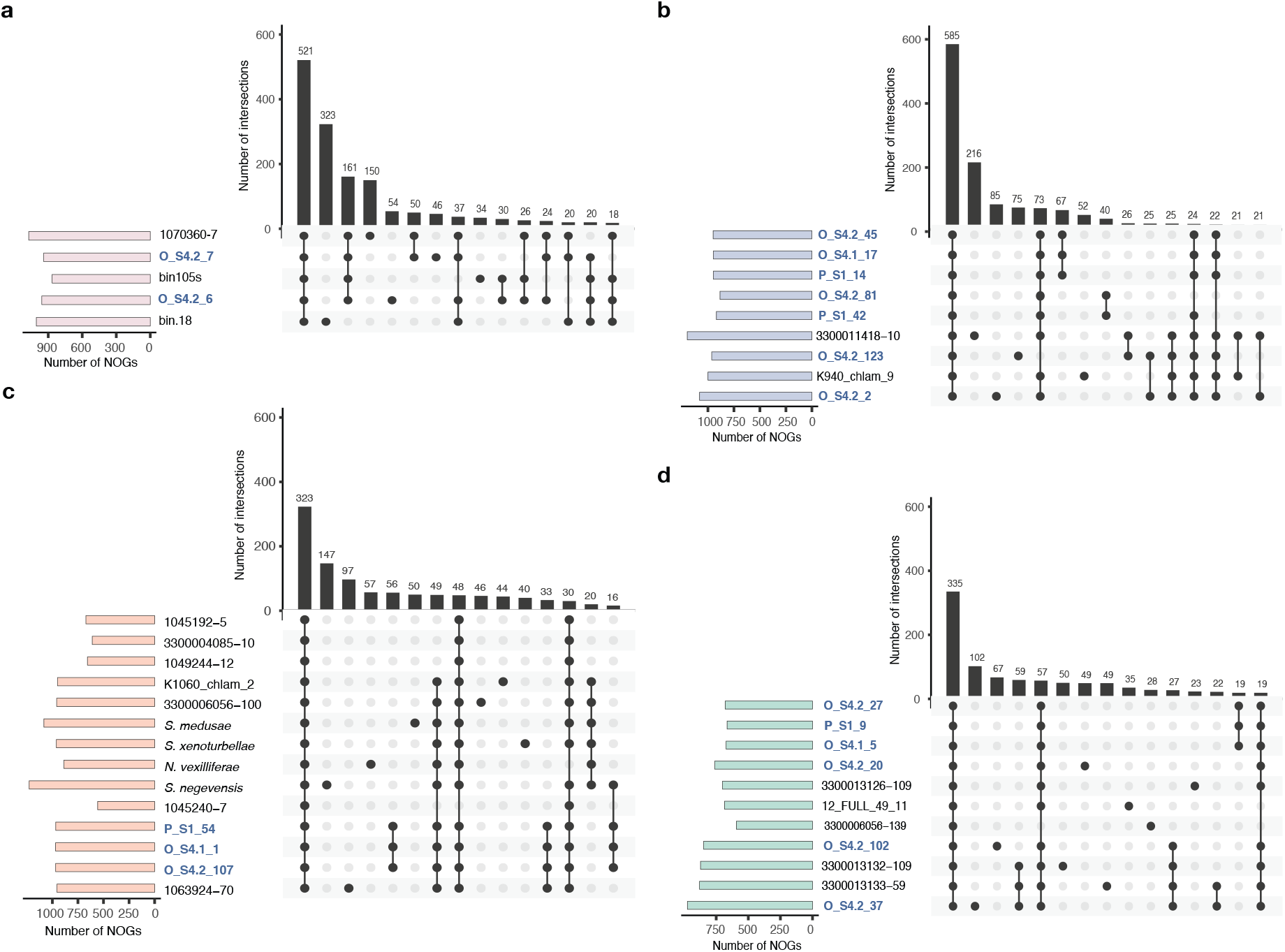
Chlamydiae families share varying levels of core gene content, with smaller sets of genes specific to sub-groups of sponge MAGs. Intersection plots give an overview of shared gene content across chlamydiae genomes from the families MCF-D (**a**), *Ca*. Sororchlamydiaceae (**b**), *Ca*. Parasimkaniaceae (**c**), and Simkaniaceae (**d**). Intersections are based on sets of NOGs found in each genome, with the total number of identified NOGs found in bar charts to the left of each taxon. Each intersection plot shows the number of NOGs shared between the taxa indicated by black circles for the top 15 most abundant intersections.

**Figure S5.**
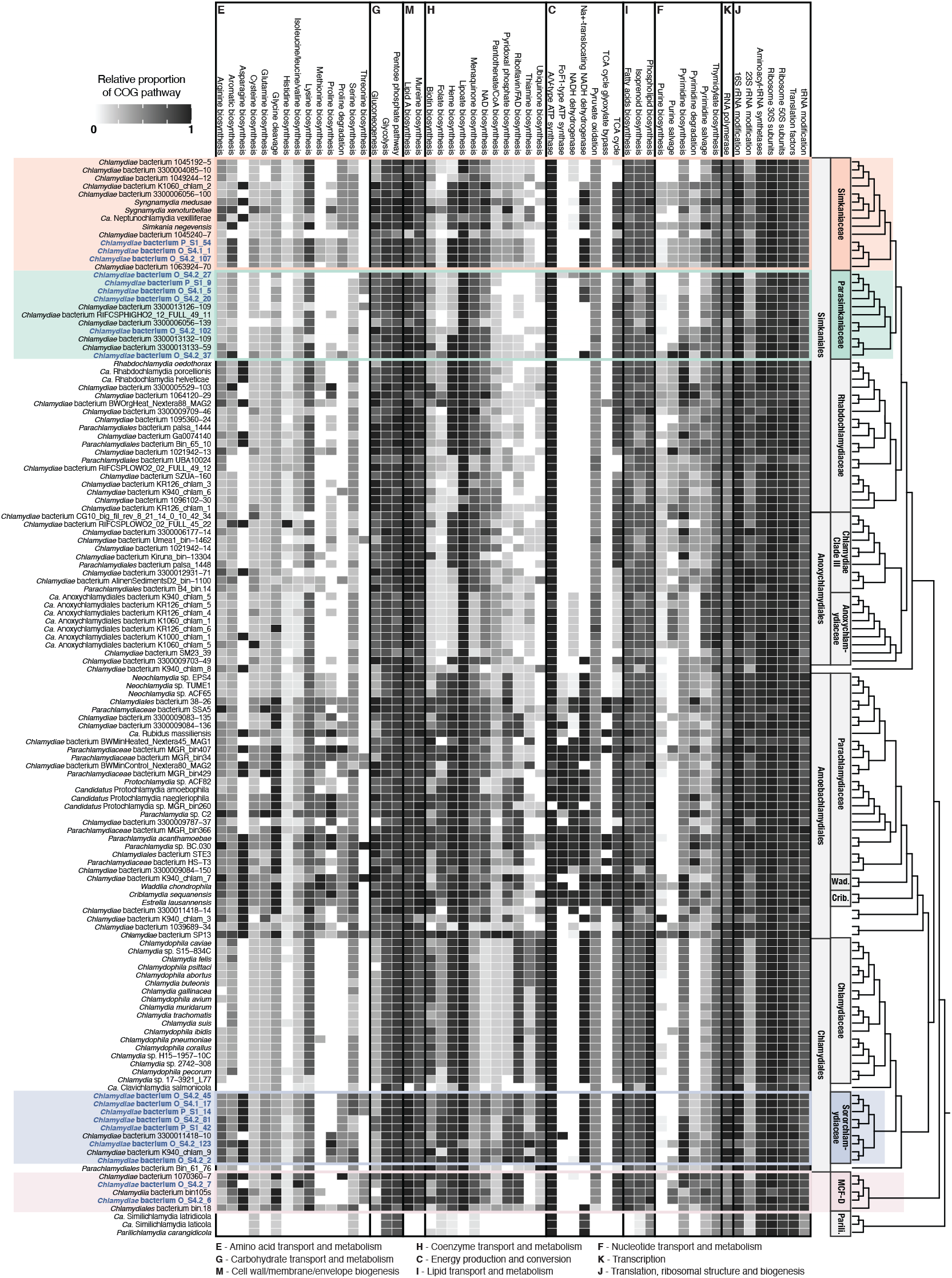
Overview of clusters of orthologous group (COG) pathways across Chlamydiae indicate that sponge-associated chlamydiae have similar metabolic profiles to other chlamydiae from their respective families. The heatmap is organized by COG category and shows the proportion of COGs from a pathway found across taxa, relative to the maximum among chlamydiae. Sponge-associated chlamydiae MAGs are in blue, with relevant families coloured accordingly. Order and family names are indicated to the right alongside a cladogram of species relationships, with the following short forms: Criblamydiaceae (Crib.), Parilichlamydiaceae (Parili.), and Waddliaceae (Wad.). See Data S9 for an overview of each COG across Chlamydiae genomes.

**Figure S6.**
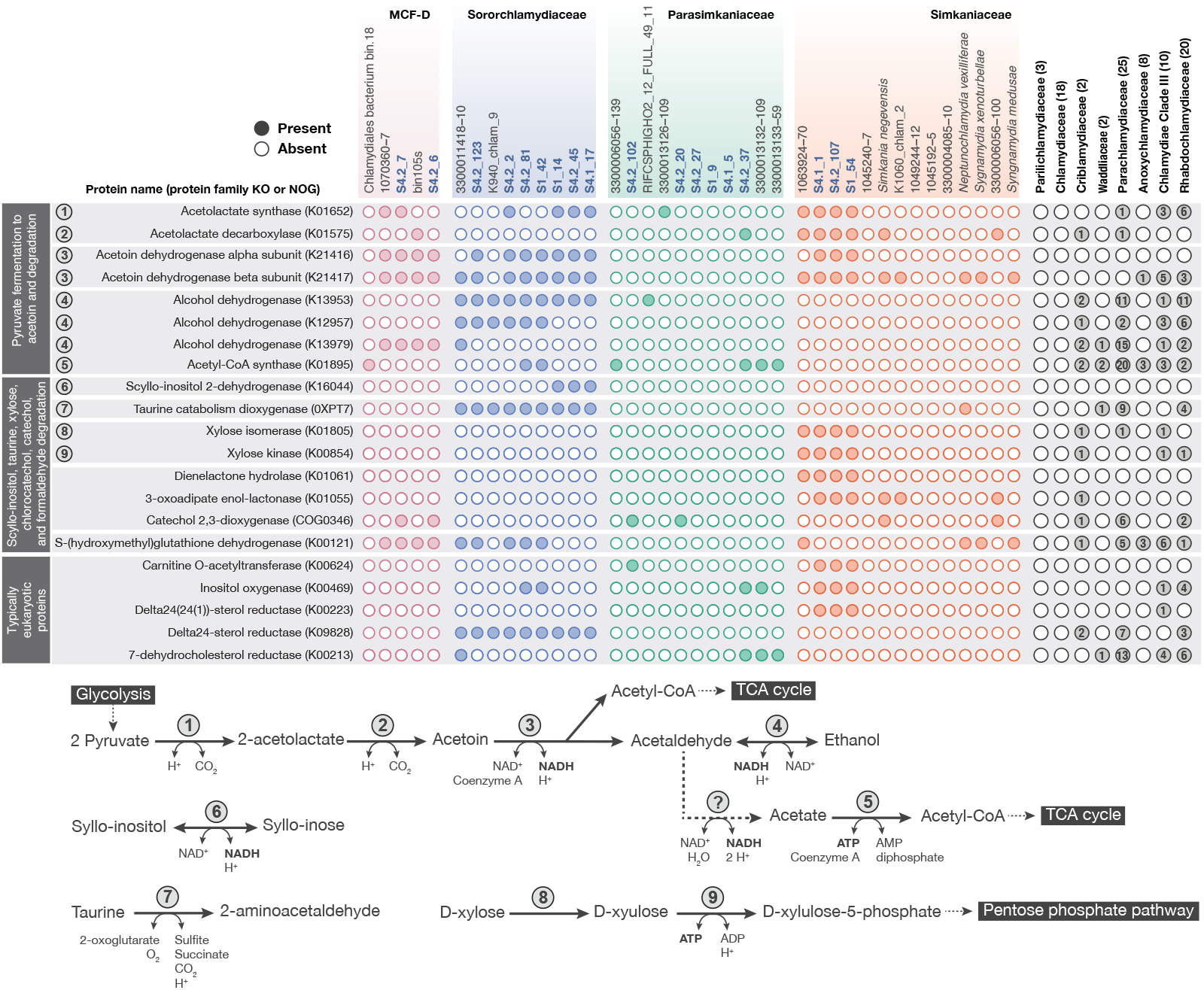
Sponge-associated chlamydiae genomes are enriched in genes related to fermentation, degradation, and genes typical of eukaryotes. The presence and absence of genes of interest across chlamydiae genomes from families with sponge-associated members is shown, alongside a schematic overview of pyruvate to acetoin fermentation, and the degradation of acetoin, syllo-inositol, taurine, and D-xylose. Gene function and protein domain are indicated to the left alongside numbers corresponding to the schematic overview. The number of representative genomes from other chlamydial families encoding the given protein family is indicated to the right. See Data S8 for a full overview of presence and absence across chlamydiae representatives and corresponding gene annotations for sponge-associated chlamydiae.

## SUPPLEMENTARY DATA

**Data S1**. Sponge sample metadata and data accessions, metagenome assembly statistics, and confirmation of sponge identity based on metagenomic SSU rRNA genes.

**Data S2**. Bacteria-specific SSU rRNA gene amplicon OTU sequences.

**Data S3**. Overview of phylum and OTU relative abundances, read counts, and taxonomy for bacteria-specific SSU rRNA gene amplicons across sponge samples.

**Data S4**. Sponge metagenome microbial diversity and MAG information. Contigs encoding SSU rRNA genes and ribosomal proteins identified in metagenomes. Overview of MAGs retrieved from each metagenome, including bin statistics and taxoomic classification.

**Data S5**. Uncollapsed phylogenetic trees, including ribosomal protein phylogeny of metagenomic contigs, species phylogenies of concatenated marker proteins, single protein phylogenies including phosphoenolpyruvate mutase and SnoaL-like polyketide cyclase, and the SSU rRNA gene phylogeny.

**Data S6**. Summary of genome characteristics, IDs, and accessions of sponge chlamydiae MAGs and PVC species representatives.

**Data S7**. Single-copy marker proteins used in concatenated species phylogenies, sequences removed during refinement, and monophyly of PVC phyla in each single-protein tree.

**Data S8**. Annotations of proteins of interest, and the number of respective NOGs and KEGG KOs across chlamydiae.

**Data S9**. Overview of the presence and absence of COGs in each COG pathway across chlamydiae, and annotations of the corresponding proteins from sponge chlamydiae MAGs.

**Data S10**. Overview of antiSMASH results with the type and number of BGCs identified in each chlamydiae species representative. BGCs with top MiBIG hits are indicated alongside top hit information.

**Data S11**. Prevalence of close relatives of sponge chlamydiae (95 % identity) across environmental samples measured by IMNGS and based on SRA SSU rRNA gene amplicon datasets. SRA samples identified with at least 0.1 % relative abundance.

