## Supplementary material for "Genomic diversity and biosynthetic capabilities of sponge-associated chlamydiae": Data S5

**Concatenated ribosomal protein phylogeny**
Metagenomic contigs encoding ribosomal proteins
in context of backbone sequences
**Maximum-likelihood**
**Rapid bootstraps**
**PROTCATLG**

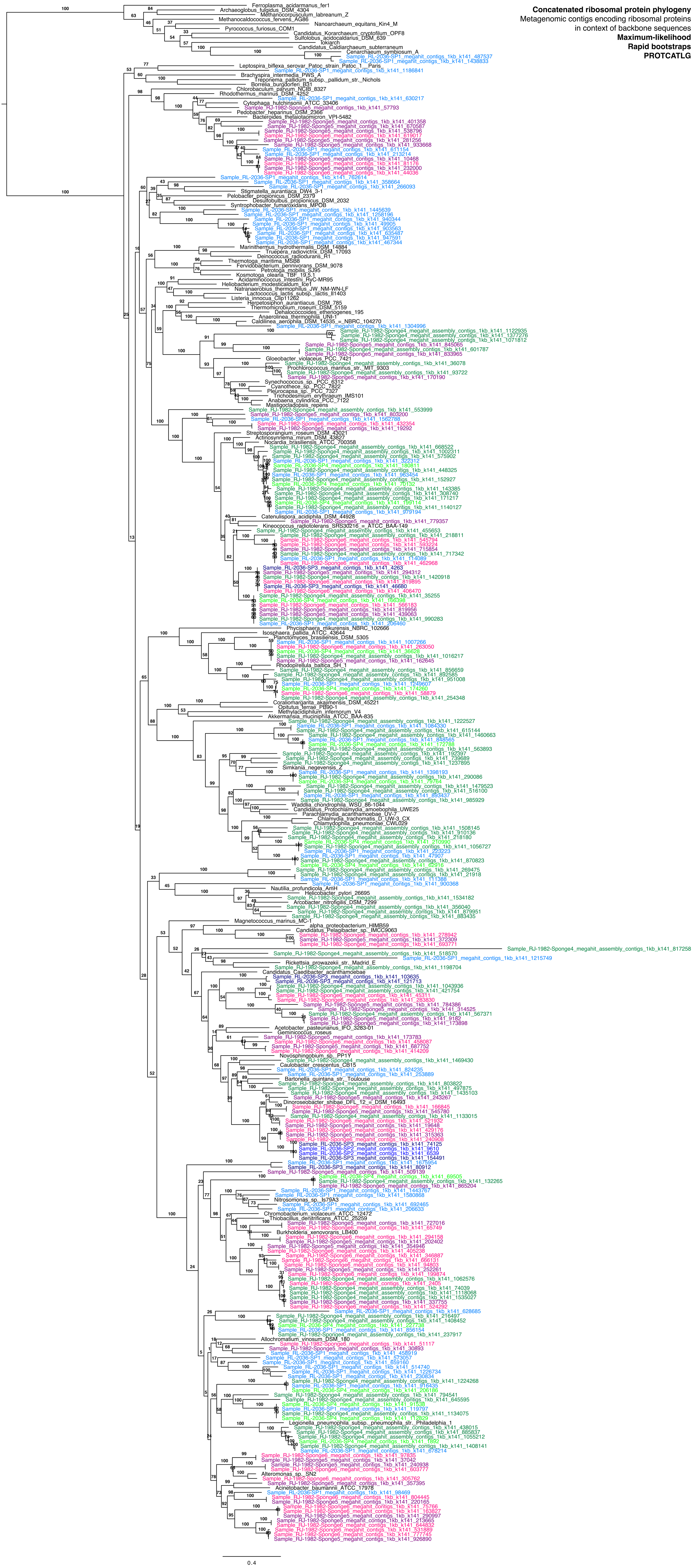

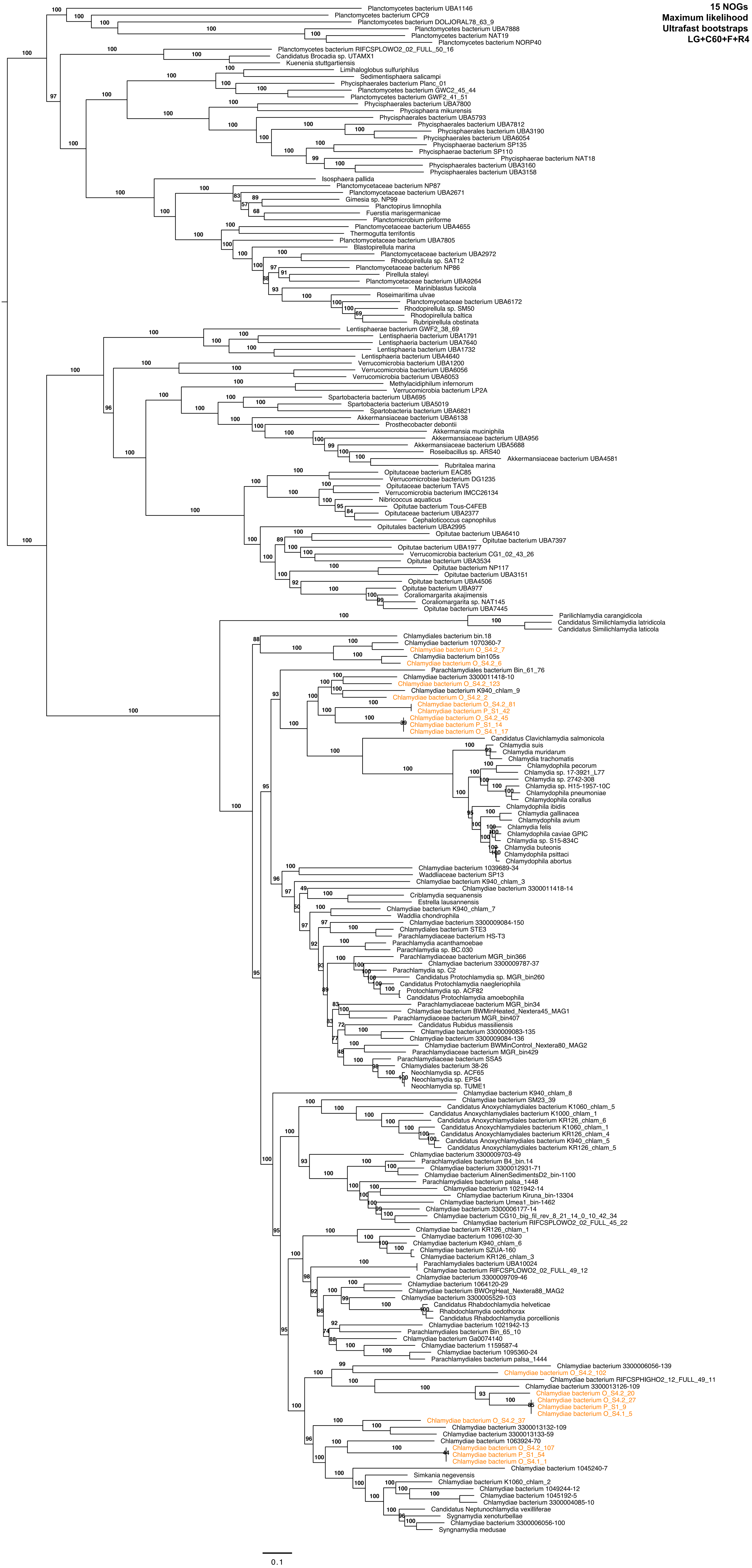
15 NOGs
Maximum likelihood
Ultrafast bootstraps
LG+C60+F+R4
Planctomycetes bacterium UBA1146
Planctomycetes bacterium CPC9
Planctomycetes bacterium DOLJORAL78_63_9
Planctomycetes bacterium UBA7888
Planctomycetes bacterium NAT19
Planctomycetes bacterium NORP40
Planctomycetes bacterium RIFCSPLOWO2_02_FULL_50_16
Candidatus Brocadia sp. UTAMX1
Kuenenia stuttgartiensis
Limihaloglobus sulfuriphilus
Sedimentisphaera salicampi
Phycisphaerales bacterium Planc_01
Planctomycetes bacterium GWC2_45_44
Planctomycetes bacterium GWF2_41_51
Phycisphaerales bacterium UBA7800
Phycisphaera mikurensis
Phycisphaerales bacterium UBA5793
Phycisphaerales bacterium UBA7812
Phycisphaerales bacterium UBA3190
Phycisphaerales bacterium UBA6054
Phycisphaerae bacterium SP135
Phycisphaerae bacterium SP110
Phycisphaerae bacterium NAT18
Phycisphaerales bacterium UBA3160
Phycisphaerales bacterium UBA3158
Isosphaera pallida
Planctomycetaceae bacterium NP87
Planctomycetaceae bacterium UBA2671
Gimesia sp. NP99
Planctopirus limnophila
Fuerstia marisgermanicae
Planctomicrobium piriforme
Planctomycetaceae bacterium UBA4655
Thermogutta terrifontis
Planctomycetaceae bacterium UBA7805
Blastopirellula marina
Planctomycetaceae bacterium UBA2972
Rhodopirellula sp. SAT12
Planctomycetaceae bacterium NP86
Pirellula staleyi
Planctomycetaceae bacterium UBA9264
Mariniblastus fucicola
Roseimaritima ulvae
Planctomycetaceae bacterium UBA6172
Rhodopirellula sp. SM50
Rhodopirellula baltica
Rubripirellula obstinata
Lentisphaerae bacterium GWF2_38_69
Lentisphaeria bacterium UBA1791
Lentisphaeria bacterium UBA7640
Lentisphaeria bacterium UBA1732
Lentisphaeria bacterium UBA4640
Verrucomicrobia bacterium UBA1200
Verrucomicrobia bacterium UBA6056
Verrucomicrobia bacterium UBA6053
Methylacidiphilum infernorum
Verrucomicrobia bacterium LP2A
Spartobacteria bacterium UBA695
Spartobacteria bacterium UBA5019
Spartobacteria bacterium UBA6821
Akkermansiaceae bacterium UBA6138
Prosthecobacter debontii
Akkermansia muciniphila
Akkermansiaceae bacterium UBA956
Akkermansiaceae bacterium UBA5688
Roseibacillus sp. ARS40
Akkermansiaceae bacterium UBA4581
Rubritalea marina
Opitutaceae bacterium EAC85
Verrucomicrobiae bacterium DG1235
Opitutaceae bacterium TAV5
Verrucomicrobia bacterium IMCC26134
Nibricoccus aquaticus
Opitutae bacterium Tous-C4FEB
Opitutaceae bacterium UBA2377
Cephaloticoccus capnophilus
Opitutales bacterium UBA2995
Opitutae bacterium UBA6410
Opitutae bacterium UBA7397
Opitutae bacterium UBA1977
Verrucomicrobia bacterium CG1_02_43_26
Opitutae bacterium UBA3534
Opitutae bacterium NP117
Opitutae bacterium UBA3151
Opitutae bacterium UBA4506
Opitutae bacterium UBA977
Coraliomargarita akajimensis
Coraliomargarita sp. NAT145
Opitutae bacterium UBA7445
Parilichlamydia carangidicola
Candidatus Similichlamydia latridicola
Candidatus Similichlamydia laticola
Chlamydiales bacterium bin.18
Chlamydiae bacterium 1070360-7
Chlamydiae bacterium O_S4.2_7
Chlamydiia bacterium bin105s
Chlamydiae bacterium O_S4.2_6
Parachlamydiales bacterium Bin_61_76
Chlamydiae bacterium 3300011418-10
Chlamydiae bacterium O_S4.2_123
Chlamydiae bacterium K940_chlam_9
Chlamydiae bacterium O_S4.2_2
Chlamydiae bacterium O_S4.2_81
Chlamydiae bacterium P_S1_42
Chlamydiae bacterium O_S4.2_45
Chlamydiae bacterium P_S1_14
Chlamydiae bacterium O_S4.1_17
Candidatus Clavichlamydia salmonicola
Chlamydia suis
Chlamydia muridarum
Chlamydia trachomatis
Chlamydophila pecorum
Chlamydia sp. 17-3921_L77
Chlamydia sp. 2742-308
Chlamydia sp. H15-1957-10C
Chlamydophila pneumoniae
Chlamydophila corallus
Chlamydophila ibidis
Chlamydia gallinacea
Chlamydia avium
Chlamydia felis
Chlamydophila caviae GPIC
Chlamydia sp. S15-834C
Chlamydia buteonis
Chlamydophila psittaci
Chlamydophila abortus
Chlamydiae bacterium 1039689-34
Waddliaceae bacterium SP13
Chlamydiae bacterium K940_chlam_3
Chlamydiae bacterium 3300011418-14
Criblamydia sequanensis
Estrella lausannensis
Chlamydiae bacterium K940_chlam_7
Waddlia chondrophila
Chlamydiae bacterium 3300009084-150
Chlamydiales bacterium STE3
Parachlamydiaceae bacterium HS-T3
Parachlamydia acanthamoebae
Parachlamydia sp. BC.030
Parachlamydiaceae bacterium MGR_bin366
Chlamydiae bacterium 3300009787-37
Parachlamydia sp. C2
Candidatus Protochlamydia sp. MGR_bin260
Candidatus Protochlamydia naegleriophila
Protochlamydia sp. ACF82
Candidatus Protochlamydia amoebophila
Parachlamydiaceae bacterium MGR_bin34
Chlamydiae bacterium BWMinHeated_Nextera45_MAG1
Parachlamydiaceae bacterium MGR_bin407
Candidatus Rubidus massiliensis
Chlamydiae bacterium 3300009083-135
Chlamydiae bacterium 3300009084-136
Chlamydiae bacterium BWMinControl_Nextera80_MAG2
Parachlamydiaceae bacterium MGR_bin429
Parachlamydiaceae bacterium SSA5
Chlamydiales bacterium 38-26
Neochlamydia sp. ACF65
Neochlamydia sp. EPS4
Neochlamydia sp. TUME1
Chlamydiae bacterium K940_chlam_8
Chlamydiae bacterium SM23_39
Candidatus Anoxychlamydiales bacterium K1060_chlam_5
Candidatus Anoxychlamydiales bacterium K1000_chlam_1
Candidatus Anoxychlamydiales bacterium KR126_chlam_6
Candidatus Anoxychlamydiales bacterium K1060_chlam_1
Candidatus Anoxychlamydiales bacterium KR126_chlam_4
Candidatus Anoxychlamydiales bacterium K940_chlam_5
Candidatus Anoxychlamydiales bacterium KR126_chlam_5
Chlamydiae bacterium 3300009703-49
Parachlamydiales bacterium B4_bin.14
Chlamydiae bacterium 3300012931-71
Chlamydiae bacterium AlinenSedimentsD2_bin-1100
Parachlamydiales bacterium palsa_1448
Chlamydiae bacterium 1021942-14
Chlamydiae bacterium Kiruna_bin-13304
Chlamydiae bacterium Umea1_bin-1462
Chlamydiae bacterium 3300006177-14
Chlamydiae bacterium CG10_big_fil_rev_8_21_14_0_10_42_34
Chlamydiae bacterium RIFCSPLOWO2_02_FULL_45_22
Chlamydiae bacterium KR126_chlam_1
Chlamydiae bacterium 1096102-30
Chlamydiae bacterium K940_chlam_6
Chlamydiae bacterium SZUA-160
Chlamydiae bacterium KR126_chlam_3
Parachlamydiales bacterium UBA10024
Chlamydiae bacterium RIFCSPLOWO2_02_FULL_49_12
Chlamydiae bacterium 3300009709-46
Chlamydiae bacterium 1064120-29
Chlamydiae bacterium BWOrgHeat_Nextera88_MAG2
Chlamydiae bacterium 3300005529-103
Candidatus Rhabdochlamydia helveticae
Rhabdochlamydia oedothorax
Candidatus Rhabdochlamydia porcellionis
Chlamydiae bacterium 1021942-13
Parachlamydiales bacterium Bin_65_10
Chlamydiae bacterium Ga0074140
Chlamydiae bacterium 1159587-4
Chlamydiae bacterium 1095360-24
Parachlamydiales bacterium palsa_1444
Chlamydiae bacterium 3300006056-139
Chlamydiae bacterium O_S4.2_102
Chlamydiae bacterium RIFCSPHIGHO2_12_FULL_49_11
Chlamydiae bacterium 3300013126-109
Chlamydiae bacterium O_S4.2_20
Chlamydiae bacterium O_S4.2_27
Chlamydiae bacterium P_S1_9
Chlamydiae bacterium O_S4.1_5
Chlamydiae bacterium O_S4.2_37
Chlamydiae bacterium 3300013132-109
Chlamydiae bacterium 3300013133-59
Chlamydiae bacterium 1063924-70
Chlamydiae bacterium O_S4.2_107
Chlamydiae bacterium P_S1_54
Chlamydiae bacterium O_S4.1_1
Chlamydiae bacterium 1045240-7
Simkania negevensis
Chlamydiae bacterium K1060_chlam_2
Chlamydiae bacterium 1049244-12
Chlamydiae bacterium 1045192-5
Chlamydiae bacterium 3300004085-10
Candidatus Neptunochlamydia vexilliferae
Sygnamydia xenoturbellae
Chlamydiae bacterium 3300006056-100
Syngnamydia medusae
0.1

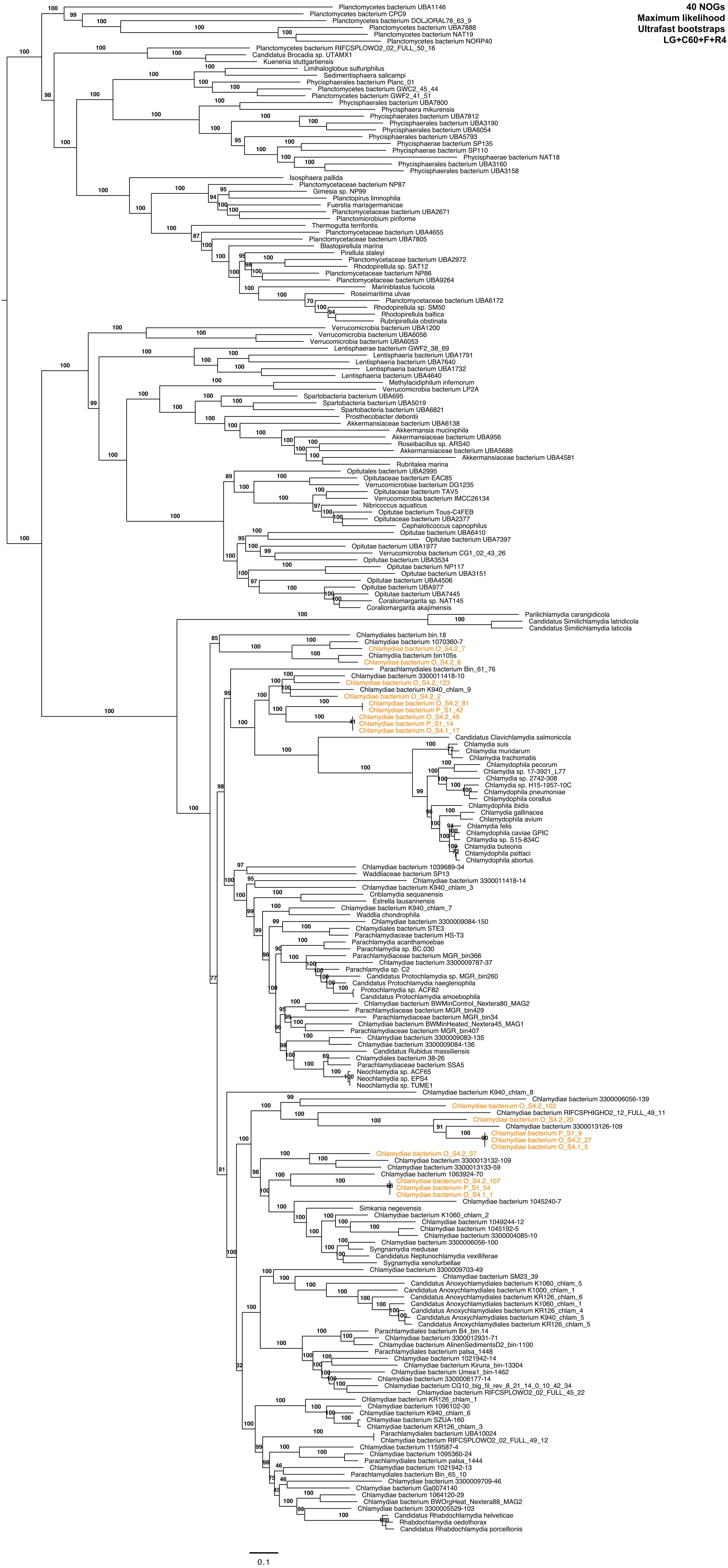
40 NOGs
Maximum likelihood
Ultrafast bootstraps
LG+C60+F+R4
Planctomycetes bacterium UBA1146
Planctomycetes bacterium CPC9
Planctomycetes bacterium DOLJORAL78_63_9
Planctomycetes bacterium UBA7888
Planctomycetes bacterium NAT19
Planctomycetes bacterium NORP40
Planctomycetes bacterium RIFCSPLOWO2_02_FULL_50_16
Candidatus Brocadia sp. UTAMX1
Kuenenia stuttgartiensis
Limihaloglobus sulfuriphilus
Sedimentisphaera salicampi
Phycisphaerales bacterium Planc_01
Planctomycetes bacterium GWC2_45_44
Planctomycetes bacterium GWF2_41_51
Phycisphaerales bacterium UBA7800
Phycisphaera mikurensis
Phycisphaerales bacterium UBA7812
Phycisphaerales bacterium UBA3190
Phycisphaerales bacterium UBA6054
Phycisphaerales bacterium UBA5793
Phycisphaerae bacterium SP135
Phycisphaerae bacterium SP110
Phycisphaerae bacterium NAT18
Phycisphaerales bacterium UBA3160
Phycisphaerales bacterium UBA3158
Isosphaera pallida
Planctomycetaceae bacterium NP87
Gimesia sp. NP99
Planctopirus limnophila
Fuerstia marisgermanicae
Planctomycetaceae bacterium UBA2671
Planctomicrobium piriforme
Thermogutta terrifontis
Planctomycetaceae bacterium UBA4655
Planctomycetaceae bacterium UBA7805
Blastopirellula marina
Pirellula staleyi
Planctomycetaceae bacterium UBA2972
Rhodopirellula sp. SAT12
Planctomycetaceae bacterium NP86
Planctomycetaceae bacterium UBA9264
Mariniblastus fucicola
Roseimaritima ulvae
Planctomycetaceae bacterium UBA6172
Rhodopirellula sp. SM50
Rhodopirellula baltica
Rubripirellula obstinata
Verrucomicrobia bacterium UBA1200
Verrucomicrobia bacterium UBA6056
Verrucomicrobia bacterium UBA6053
Lentisphaerae bacterium GWF2_38_69
Lentisphaeria bacterium UBA1791
Lentisphaeria bacterium UBA7640
Lentisphaeria bacterium UBA1732
Lentisphaeria bacterium UBA4640
Methylacidiphilum infernorum
Verrucomicrobia bacterium LP2A
Spartobacteria bacterium UBA695
Spartobacteria bacterium UBA5019
Spartobacteria bacterium UBA6821
Prosthecobacter debontii
Akkermansiaceae bacterium UBA6138
Akkermansia muciniphila
Akkermansiaceae bacterium UBA956
Roseibacillus sp. ARS40
Akkermansiaceae bacterium UBA5688
Akkermansiaceae bacterium UBA4581
Rubritalea marina
Opitutales bacterium UBA2995
Opitutaceae bacterium EAC85
Verrucomicrobiae bacterium DG1235
Opitutaceae bacterium TAV5
Verrucomicrobia bacterium IMCC26134
Nibricoccus aquaticus
Opitutae bacterium Tous-C4FEB
Opitutaceae bacterium UBA2377
Cephaloticoccus capnophilus
Opitutae bacterium UBA6410
Opitutae bacterium UBA7397
Opitutae bacterium UBA1977
Verrucomicrobia bacterium CG1_02_43_26
Opitutae bacterium UBA3534
Opitutae bacterium NP117
Opitutae bacterium UBA3151
Opitutae bacterium UBA4506
Opitutae bacterium UBA977
Opitutae bacterium UBA7445
Coraliomargarita sp. NAT145
Coraliomargarita akajimensis
Parilichlamydia carangidicola
Candidatus Similichlamydia latridicola
Candidatus Similichlamydia laticola
Chlamydiales bacterium bin.18
Chlamydiae bacterium 1070360-7
Chlamydiae bacterium O_S4.2_7
Chlamydiia bacterium bin105s
Chlamydiae bacterium O_S4.2_6
Parachlamydiales bacterium Bin_61_76
Chlamydiae bacterium 3300011418-10
Chlamydiae bacterium O_S4.2_123
Chlamydiae bacterium K940_chlam_9
Chlamydiae bacterium O_S4.2_2
Chlamydiae bacterium O_S4.2_81
Chlamydiae bacterium P_S1_42
Chlamydiae bacterium O_S4.2_45
Chlamydiae bacterium P_S1_14
Chlamydiae bacterium O_S4.1_17
Candidatus Clavichlamydia salmonicola
Chlamydia suis
Chlamydia muridarum
Chlamydia trachomatis
Chlamydophila pecorum
Chlamydia sp. 17-3921_L77
Chlamydia sp. 2742-308
Chlamydia sp. H15-1957-10C
Chlamydophila pneumoniae
Chlamydophila corallus
Chlamydophila ibidis
Chlamydia gallinacea
Chlamydophila avium
Chlamydia felis
Chlamydophila caviae GPIC
Chlamydia sp. S15-834C
Chlamydia buteonis
Chlamydophila psittaci
Chlamydophila abortus
Chlamydiae bacterium 1039689-34
Waddliaceae bacterium SP13
Chlamydiae bacterium 3300011418-14
Chlamydiae bacterium K940_chlam_3
Criblamydia sequanensis
Estrella lausannensis
Chlamydiae bacterium K940_chlam_7
Waddlia chondrophila
Chlamydiae bacterium 3300009084-150
Chlamydiales bacterium STE3
Parachlamydiaceae bacterium HS-T3
Parachlamydia acanthamoebae
Parachlamydia sp. BC.030
Parachlamydiaceae bacterium MGR_bin366
Chlamydiae bacterium 3300009787-37
Parachlamydia sp. C2
Candidatus Protochlamydia sp. MGR_bin260
Candidatus Protochlamydia naegleriophila
Protochlamydia sp. ACF82
Candidatus Protochlamydia amoebophila
Chlamydiae bacterium BWMinControl_Nextera80_MAG2
Parachlamydiaceae bacterium MGR_bin429
Parachlamydiaceae bacterium MGR_bin34
Chlamydiae bacterium BWMinHeated_Nextera45_MAG1
Parachlamydiaceae bacterium MGR_bin407
Chlamydiae bacterium 3300009083-135
Chlamydiae bacterium 3300009084-136
Candidatus Rubidus massiliensis
Chlamydiales bacterium 38-26
Parachlamydiaceae bacterium SSA5
Neochlamydia sp. ACF65
Neochlamydia sp. EPS4
Neochlamydia sp. TUME1
Chlamydiae bacterium K940_chlam_8
Chlamydiae bacterium 3300006056-139
Chlamydiae bacterium O_S4.2_102
Chlamydiae bacterium RIFCSPHIGHO2_12_FULL_49_11
Chlamydiae bacterium O_S4.2_20
Chlamydiae bacterium 3300013126-109
Chlamydiae bacterium P_S1_9
Chlamydiae bacterium O_S4.2_27
Chlamydiae bacterium O_S4.1_5
Chlamydiae bacterium O_S4.2_37
Chlamydiae bacterium 3300013132-109
Chlamydiae bacterium 3300013133-59
Chlamydiae bacterium 1063924-70
Chlamydiae bacterium O_S4.2_107
Chlamydiae bacterium P_S1_54
Chlamydiae bacterium O_S4.1_1
Chlamydiae bacterium 1045240-7
Simkania negevensis
Chlamydiae bacterium K1060_chlam_2
Chlamydiae bacterium 1049244-12
Chlamydiae bacterium 1045192-5
Chlamydiae bacterium 3300004085-10
Chlamydiae bacterium 3300006056-100
Syngnamydia medusae
Candidatus Neptunochlamydia vexilliferae
Sygnamydia xenoturbellae
Chlamydiae bacterium 3300009703-49
Chlamydiae bacterium SM23_39
Candidatus Anoxychlamydiales bacterium K1060_chlam_5
Candidatus Anoxychlamydiales bacterium K1000_chlam_1
Candidatus Anoxychlamydiales bacterium KR126_chlam_6
Candidatus Anoxychlamydiales bacterium K1060_chlam_1
Candidatus Anoxychlamydiales bacterium KR126_chlam_4
Candidatus Anoxychlamydiales bacterium K940_chlam_5
Candidatus Anoxychlamydiales bacterium KR126_chlam_5
Parachlamydiales bacterium B4_bin.14
Chlamydiae bacterium 3300012931-71
Chlamydiae bacterium AlinenSedimentsD2_bin-1100
Parachlamydiales bacterium palsa_1448
Chlamydiae bacterium 1021942-14
Chlamydiae bacterium Kiruna_bin-13304
Chlamydiae bacterium Umea1_bin-1462
Chlamydiae bacterium 3300006177-14
Chlamydiae bacterium CG10_big_fil_rev_8_21_14_0_10_42_34
Chlamydiae bacterium RIFCSPLOWO2_02_FULL_45_22
Chlamydiae bacterium KR126_chlam_1
Chlamydiae bacterium 1096102-30
Chlamydiae bacterium K940_chlam_6
Chlamydiae bacterium SZUA-160
Chlamydiae bacterium KR126_chlam_3
Parachlamydiales bacterium UBA10024
Chlamydiae bacterium RIFCSPLOWO2_02_FULL_49_12
Chlamydiae bacterium 1159587-4
Chlamydiae bacterium 1095360-24
Parachlamydiales bacterium palsa_1444
Chlamydiae bacterium 1021942-13
Parachlamydiales bacterium Bin_65_10
Chlamydiae bacterium 3300009709-46
Chlamydiae bacterium Ga0074140
Chlamydiae bacterium 1064120-29
Chlamydiae bacterium BWOrgHeat_Nextera88_MAG2
Chlamydiae bacterium 3300005529-103
Candidatus Rhabdochlamydia helveticae
Rhabdochlamydia oedothorax
Candidatus Rhabdochlamydia porcellionis
0.1

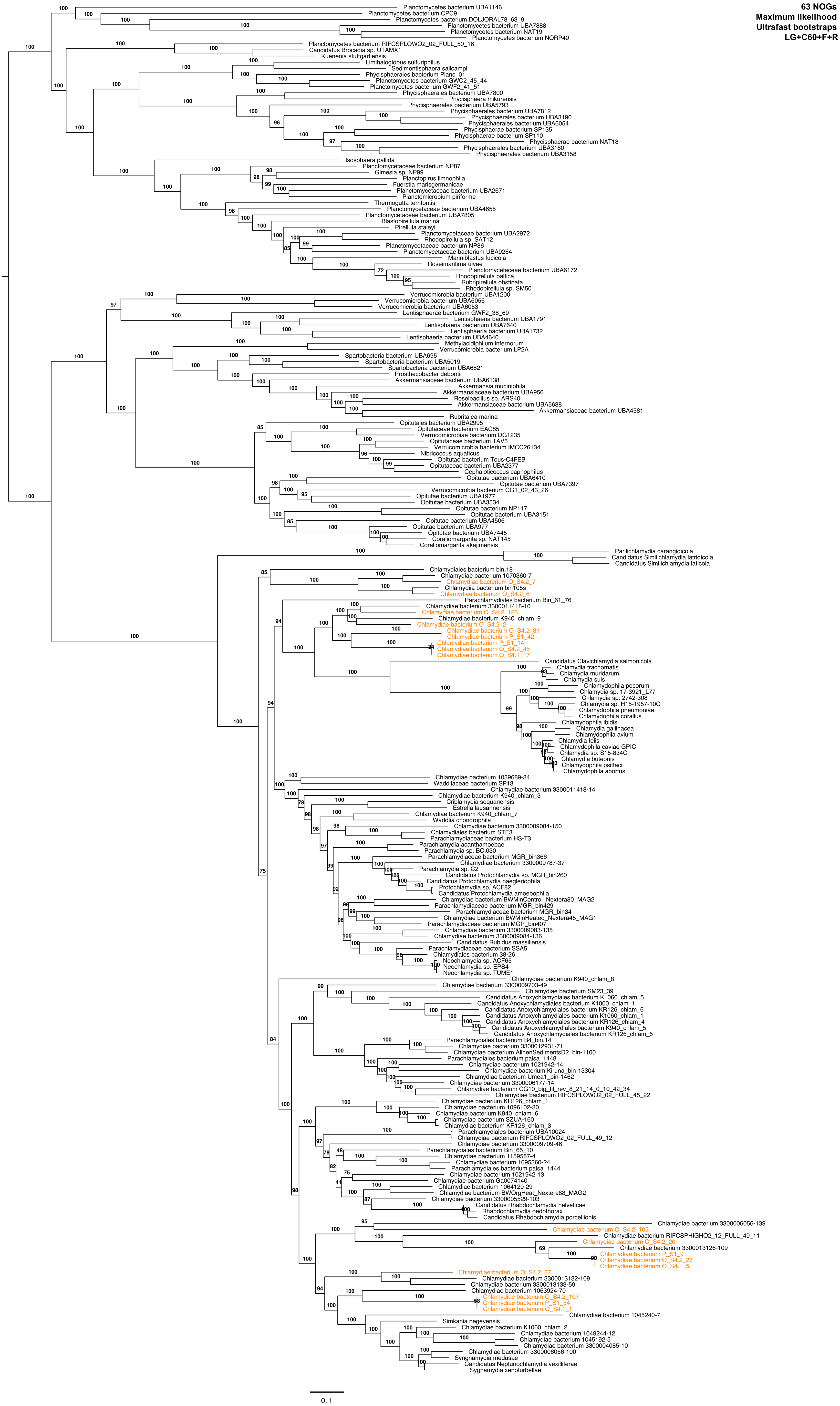
63 NOGs
Maximum likelihood
Ultrafast bootstraps
LG+C60+F+R
Planctomycetes bacterium UBA1146
Planctomycetes bacterium CPC9
Planctomycetes bacterium DOLJORAL78_63_9
Planctomycetes bacterium UBA7888
Planctomycetes bacterium NAT19
Planctomycetes bacterium NORP40
Planctomycetes bacterium RIFCSPLOWO2_02_FULL_50_16
Candidatus Brocadia sp. UTAMX1
Kuenenia stuttgartiensis
Limihaloglobus sulfuriphilus
Sedimentisphaera salicampi
Phycisphaerales bacterium Planc_01
Planctomycetes bacterium GWC2_45_44
Planctomycetes bacterium GWF2_41_51
Phycisphaerales bacterium UBA7800
Phycisphaera mikurensis
Phycisphaerales bacterium UBA5793
Phycisphaerales bacterium UBA7812
Phycisphaerales bacterium UBA3190
Phycisphaerales bacterium UBA6054
Phycisphaerae bacterium SP135
Phycisphaerae bacterium SP110
Phycisphaerae bacterium NAT18
Phycisphaerales bacterium UBA3160
Phycisphaerales bacterium UBA3158
Isosphaera pallida
Planctomycetaceae bacterium NP87
Gimesia sp. NP99
Planctopirus limnophila
Fuerstia marisgermanicae
Planctomycetaceae bacterium UBA2671
Planctomicrobium piriforme
Thermogutta terrifontis
Planctomycetaceae bacterium UBA4655
Planctomycetaceae bacterium UBA7805
Blastopirellula marina
Pirellula staleyi
Planctomycetaceae bacterium UBA2972
Rhodopirellula sp. SAT12
Planctomycetaceae bacterium NP86
Planctomycetaceae bacterium UBA9264
Mariniblastus fucicola
Roseimaritima ulvae
Planctomycetaceae bacterium UBA6172
Rhodopirellula baltica
Rubripirellula obstinata
Rhodopirellula sp. SM50
Verrucomicrobia bacterium UBA1200
Verrucomicrobia bacterium UBA6056
Verrucomicrobia bacterium UBA6053
Lentisphaerae bacterium GWF2_38_69
Lentisphaeria bacterium UBA1791
Lentisphaeria bacterium UBA7640
Lentisphaeria bacterium UBA1732
Lentisphaeria bacterium UBA4640
Methylacidiphilum infernorum
Verrucomicrobia bacterium LP2A
Spartobacteria bacterium UBA695
Spartobacteria bacterium UBA5019
Spartobacteria bacterium UBA6821
Prosthecobacter debontii
Akkermansiaceae bacterium UBA6138
Akkermansia muciniphila
Akkermansiaceae bacterium UBA956
Roseibacillus sp. ARS40
Akkermansiaceae bacterium UBA5688
Akkermansiaceae bacterium UBA4581
Rubritalea marina
Opitutales bacterium UBA2995
Opitutaceae bacterium EAC85
Verrucomicrobiae bacterium DG1235
Opitutaceae bacterium TAV5
Verrucomicrobia bacterium IMCC26134
Nibricoccus aquaticus
Opitutae bacterium Tous-C4FEB
Opitutaceae bacterium UBA2377
Cephaloticoccus capnophilus
Opitutae bacterium UBA6410
Opitutae bacterium UBA7397
Verrucomicrobia bacterium CG1_02_43_26
Opitutae bacterium UBA1977
Opitutae bacterium UBA3534
Opitutae bacterium NP117
Opitutae bacterium UBA3151
Opitutae bacterium UBA4506
Opitutae bacterium UBA977
Opitutae bacterium UBA7445
Coraliomargarita sp. NAT145
Coraliomargarita akajimensis
Parilichlamydia carangidicola
Candidatus Similichlamydia latridicola
Candidatus Similichlamydia laticola
Chlamydiales bacterium bin.18
Chlamydiae bacterium 1070360-7
Chlamydiae bacterium O_S4.2_7
Chlamydiia bacterium bin105s
Chlamydiae bacterium O_S4.2_6
Parachlamydiales bacterium Bin_61_76
Chlamydiae bacterium 3300011418-10
Chlamydiae bacterium O_S4.2_123
Chlamydiae bacterium K940_chlam_9
Chlamydiae bacterium O_S4.2_2
Chlamydiae bacterium O_S4.2_81
Chlamydiae bacterium P_S1_42
Chlamydiae bacterium P_S1_14
Chlamydiae bacterium O_S4.2_45
Chlamydiae bacterium O_S4.1_17
Candidatus Clavichlamydia salmonicola
Chlamydia trachomatis
Chlamydia muridarum
Chlamydia suis
Chlamydophila pecorum
Chlamydia sp. 17-3921_L77
Chlamydia sp. 2742-308
Chlamydia sp. H15-1957-10C
Chlamydophila pneumoniae
Chlamydophila corallus
Chlamydophila ibidis
Chlamydia gallinacea
Chlamydophila avium
Chlamydia felis
Chlamydophila caviae GPIC
Chlamydia sp. S15-834C
Chlamydia buteonis
Chlamydophila psittaci
Chlamydophila abortus
Chlamydiae bacterium 1039689-34
Waddliaceae bacterium SP13
Chlamydiae bacterium 3300011418-14
Chlamydiae bacterium K940_chlam_3
Criblamydia sequanensis
Estrella lausannensis
Chlamydiae bacterium K940_chlam_7
Waddlia chondrophila
Chlamydiae bacterium 3300009084-150
Chlamydiales bacterium STE3
Parachlamydiaceae bacterium HS-T3
Parachlamydia acanthamoebae
Parachlamydia sp. BC.030
Parachlamydiaceae bacterium MGR_bin366
Chlamydiae bacterium 3300009787-37
Parachlamydia sp. C2
Candidatus Protochlamydia sp. MGR_bin260
Candidatus Protochlamydia naegleriophila
Protochlamydia sp. ACF82
Candidatus Protochlamydia amoebophila
Chlamydiae bacterium BWMinControl_Nextera80_MAG2
Parachlamydiaceae bacterium MGR_bin429
Parachlamydiaceae bacterium MGR_bin34
Chlamydiae bacterium BWMinHeated_Nextera45_MAG1
Parachlamydiaceae bacterium MGR_bin407
Chlamydiae bacterium 3300009083-135
Chlamydiae bacterium 3300009084-136
Candidatus Rubidus massiliensis
Parachlamydiaceae bacterium SSA5
Chlamydiales bacterium 38-26
Neochlamydia sp. ACF65
Neochlamydia sp. EPS4
Neochlamydia sp. TUME1
Chlamydiae bacterium K940_chlam_8
Chlamydiae bacterium 3300009703-49
Chlamydiae bacterium SM23_39
Candidatus Anoxychlamydiales bacterium K1060_chlam_5
Candidatus Anoxychlamydiales bacterium K1000_chlam_1
Candidatus Anoxychlamydiales bacterium KR126_chlam_6
Candidatus Anoxychlamydiales bacterium K1060_chlam_1
Candidatus Anoxychlamydiales bacterium KR126_chlam_4
Candidatus Anoxychlamydiales bacterium K940_chlam_5
Candidatus Anoxychlamydiales bacterium KR126_chlam_5
Parachlamydiales bacterium B4_bin.14
Chlamydiae bacterium 3300012931-71
Chlamydiae bacterium AlinenSedimentsD2_bin-1100
Parachlamydiales bacterium palsa_1448
Chlamydiae bacterium 1021942-14
Chlamydiae bacterium Kiruna_bin-13304
Chlamydiae bacterium Umea1_bin-1462
Chlamydiae bacterium 3300006177-14
Chlamydiae bacterium CG10_big_fil_rev_8_21_14_0_10_42_34
Chlamydiae bacterium RIFCSPLOWO2_02_FULL_45_22
Chlamydiae bacterium KR126_chlam_1
Chlamydiae bacterium 1096102-30
Chlamydiae bacterium K940_chlam_6
Chlamydiae bacterium SZUA-160
Chlamydiae bacterium KR126_chlam_3
Parachlamydiales bacterium UBA10024
Chlamydiae bacterium RIFCSPLOWO2_02_FULL_49_12
Chlamydiae bacterium 3300009709-46
Parachlamydiales bacterium Bin_65_10
Chlamydiae bacterium 1159587-4
Chlamydiae bacterium 1095360-24
Parachlamydiales bacterium palsa_1444
Chlamydiae bacterium 1021942-13
Chlamydiae bacterium Ga0074140
Chlamydiae bacterium 1064120-29
Chlamydiae bacterium BWOrgHeat_Nextera88_MAG2
Chlamydiae bacterium 3300005529-103
Candidatus Rhabdochlamydia helveticae
Rhabdochlamydia oedothorax
Candidatus Rhabdochlamydia porcellionis
Chlamydiae bacterium 3300006056-139
Chlamydiae bacterium O_S4.2_102
Chlamydiae bacterium RIFCSPHIGHO2_12_FULL_49_11
Chlamydiae bacterium O_S4.2_20
Chlamydiae bacterium 3300013126-109
Chlamydiae bacterium P_S1_9
Chlamydiae bacterium O_S4.2_27
Chlamydiae bacterium O_S4.1_5
Chlamydiae bacterium O_S4.2_37
Chlamydiae bacterium 3300013132-109
Chlamydiae bacterium 3300013133-59
Chlamydiae bacterium 1063924-70
Chlamydiae bacterium O_S4.2_107
Chlamydiae bacterium P_S1_54
Chlamydiae bacterium O_S4.1_1
Chlamydiae bacterium 1045240-7
Simkania negevensis
Chlamydiae bacterium K1060_chlam_2
Chlamydiae bacterium 1049244-12
Chlamydiae bacterium 1045192-5
Chlamydiae bacterium 3300004085-10
Chlamydiae bacterium 3300006056-100
Syngnamydia medusae
Candidatus Neptunochlamydia vexilliferae
Sygnamydia xenoturbellae
0.1

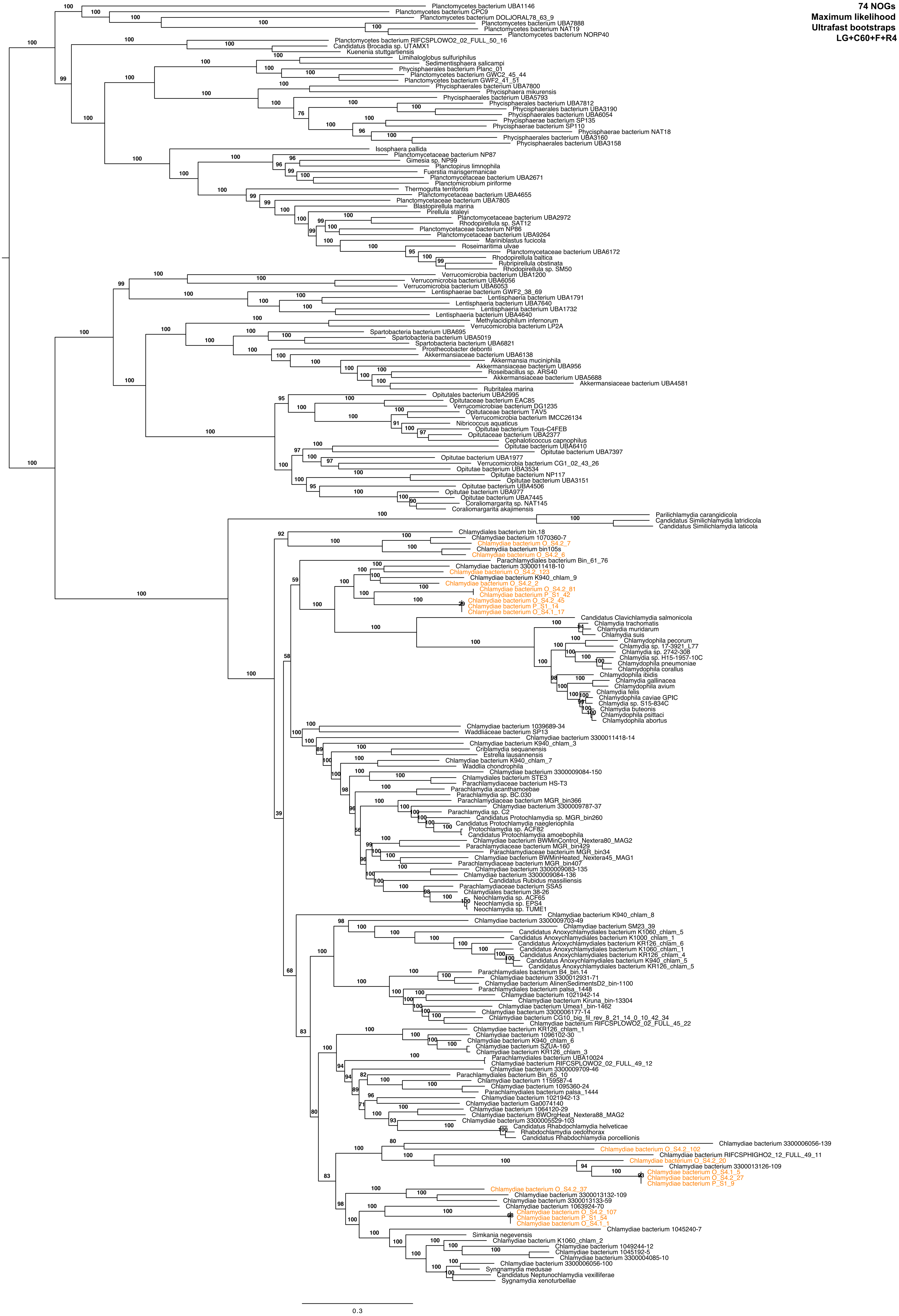
74 NOGs
Maximum likelihood
Ultrafast bootstraps
LG+C60+F+R4
Planctomycetes bacterium UBA1146
Planctomycetes bacterium CPC9
Planctomycetes bacterium DOLJORAL78_63_9
Planctomycetes bacterium UBA7888
Planctomycetes bacterium NAT19
Planctomycetes bacterium NORP40
Planctomycetes bacterium RIFCSPLOWO2_02_FULL_50_16
Candidatus Brocadia sp. UTAMX1
Kuenenia stuttgartiensis
Limihaloglobus sulfuriphilus
Sedimentisphaera salicampi
Phycisphaerales bacterium Planc_01
Planctomycetes bacterium GWC2_45_44
Planctomycetes bacterium GWF2_41_51
Phycisphaerales bacterium UBA7800
Phycisphaera mikurensis
Phycisphaerales bacterium UBA5793
Phycisphaerales bacterium UBA7812
Phycisphaerales bacterium UBA3190
Phycisphaerales bacterium UBA6054
Phycisphaerae bacterium SP135
Phycisphaerae bacterium SP110
Phycisphaerae bacterium NAT18
Phycisphaerales bacterium UBA3160
Phycisphaerales bacterium UBA3158
Isosphaera pallida
Planctomycetaceae bacterium NP87
Gimesia sp. NP99
Planctopirus limnophila
Fuerstia marisgermanicae
Planctomycetaceae bacterium UBA2671
Planctomicrobium piriforme
Thermogutta terrifontis
Planctomycetaceae bacterium UBA4655
Planctomycetaceae bacterium UBA7805
Blastopirellula marina
Pirellula staleyi
Planctomycetaceae bacterium UBA2972
Rhodopirellula sp. SAT12
Planctomycetaceae bacterium NP86
Planctomycetaceae bacterium UBA9264
Mariniblastus fucicola
Roseimaritima ulvae
Planctomycetaceae bacterium UBA6172
Rhodopirellula baltica
Rubripirellula obstinata
Rhodopirellula sp. SM50
Verrucomicrobia bacterium UBA1200
Verrucomicrobia bacterium UBA6056
Verrucomicrobia bacterium UBA6053
Lentisphaerae bacterium GWF2_38_69
Lentisphaeria bacterium UBA1791
Lentisphaeria bacterium UBA7640
Lentisphaeria bacterium UBA1732
Lentisphaeria bacterium UBA4640
Methylacidiphilum infernorum
Verrucomicrobia bacterium LP2A
Spartobacteria bacterium UBA695
Spartobacteria bacterium UBA5019
Spartobacteria bacterium UBA6821
Prosthecobacter debontii
Akkermansiaceae bacterium UBA6138
Akkermansia muciniphila
Akkermansiaceae bacterium UBA956
Roseibacillus sp. ARS40
Akkermansiaceae bacterium UBA5688
Akkermansiaceae bacterium UBA4581
Rubritalea marina
Opitutales bacterium UBA2995
Opitutaceae bacterium EAC85
Verrucomicrobiae bacterium DG1235
Opitutaceae bacterium TAV5
Verrucomicrobia bacterium IMCC26134
Nibricoccus aquaticus
Opitutae bacterium Tous-C4FEB
Opitutaceae bacterium UBA2377
Cephaloticoccus capnophilus
Opitutae bacterium UBA6410
Opitutae bacterium UBA7397
Opitutae bacterium UBA1977
Verrucomicrobia bacterium CG1_02_43_26
Opitutae bacterium UBA3534
Opitutae bacterium NP117
Opitutae bacterium UBA3151
Opitutae bacterium UBA4506
Opitutae bacterium UBA977
Opitutae bacterium UBA7445
Coraliomargarita sp. NAT145
Coraliomargarita akajimensis
Parilichlamydia carangidicola
Candidatus Similichlamydia latridicola
Candidatus Similichlamydia laticola
Chlamydiales bacterium bin.18
Chlamydiae bacterium 1070360-7
Chlamydiae bacterium O_S4.2_7
Chlamydiia bacterium bin105s
Chlamydiae bacterium O_S4.2_6
Parachlamydiales bacterium Bin_61_76
Chlamydiae bacterium 3300011418-10
Chlamydiae bacterium O_S4.2_123
Chlamydiae bacterium K940_chlam_9
Chlamydiae bacterium O_S4.2_2
Chlamydiae bacterium O_S4.2_81
Chlamydiae bacterium P_S1_42
Chlamydiae bacterium O_S4.2_45
Chlamydiae bacterium P_S1_14
Chlamydiae bacterium O_S4.1_17
Candidatus Clavichlamydia salmonicola
Chlamydia trachomatis
Chlamydia muridarum
Chlamydia suis
Chlamydophila pecorum
Chlamydia sp. 17-3921_L77
Chlamydia sp. 2742-308
Chlamydia sp. H15-1957-10C
Chlamydophila pneumoniae
Chlamydophila corallus
Chlamydophila ibidis
Chlamydia gallinacea
Chlamydophila avium
Chlamydia felis
Chlamydophila caviae GPIC
Chlamydia sp. S15-834C
Chlamydia buteonis
Chlamydophila psittaci
Chlamydophila abortus
Chlamydiae bacterium 1039689-34
Waddliaceae bacterium SP13
Chlamydiae bacterium 3300011418-14
Chlamydiae bacterium K940_chlam_3
Criblamydia sequanensis
Estrella lausannensis
Chlamydiae bacterium K940_chlam_7
Waddlia chondrophila
Chlamydiae bacterium 3300009084-150
Chlamydiales bacterium STE3
Parachlamydiaceae bacterium HS-T3
Parachlamydia acanthamoebae
Parachlamydia sp. BC.030
Parachlamydiaceae bacterium MGR_bin366
Chlamydiae bacterium 3300009787-37
Parachlamydia sp. C2
Candidatus Protochlamydia sp. MGR_bin260
Candidatus Protochlamydia naegleriophila
Protochlamydia sp. ACF82
Candidatus Protochlamydia amoebophila
Chlamydiae bacterium BWMinControl_Nextera80_MAG2
Parachlamydiaceae bacterium MGR_bin429
Parachlamydiaceae bacterium MGR_bin34
Chlamydiae bacterium BWMinHeated_Nextera45_MAG1
Parachlamydiaceae bacterium MGR_bin407
Chlamydiae bacterium 3300009083-135
Chlamydiae bacterium 3300009084-136
Candidatus Rubidus massiliensis
Parachlamydiaceae bacterium SSA5
Chlamydiales bacterium 38-26
Neochlamydia sp. ACF65
Neochlamydia sp. EPS4
Neochlamydia sp. TUME1
Chlamydiae bacterium K940_chlam_8
Chlamydiae bacterium 3300009703-49
Chlamydiae bacterium SM23_39
Candidatus Anoxychlamydiales bacterium K1060_chlam_5
Candidatus Anoxychlamydiales bacterium K1000_chlam_1
Candidatus Anoxychlamydiales bacterium KR126_chlam_6
Candidatus Anoxychlamydiales bacterium K1060_chlam_1
Candidatus Anoxychlamydiales bacterium KR126_chlam_4
Candidatus Anoxychlamydiales bacterium K940_chlam_5
Candidatus Anoxychlamydiales bacterium KR126_chlam_5
Parachlamydiales bacterium B4_bin.14
Chlamydiae bacterium 3300012931-71
Chlamydiae bacterium AlinenSedimentsD2_bin-1100
Parachlamydiales bacterium palsa_1448
Chlamydiae bacterium 1021942-14
Chlamydiae bacterium Kiruna_bin-13304
Chlamydiae bacterium Umea1_bin-1462
Chlamydiae bacterium 3300006177-14
Chlamydiae bacterium CG10_big_fil_rev_8_21_14_0_10_42_34
Chlamydiae bacterium RIFCSPLOWO2_02_FULL_45_22
Chlamydiae bacterium KR126_chlam_1
Chlamydiae bacterium 1096102-30
Chlamydiae bacterium K940_chlam_6
Chlamydiae bacterium SZUA-160
Chlamydiae bacterium KR126_chlam_3
Parachlamydiales bacterium UBA10024
Chlamydiae bacterium RIFCSPLOWO2_02_FULL_49_12
Chlamydiae bacterium 3300009709-46
Parachlamydiales bacterium Bin_65_10
Chlamydiae bacterium 1159587-4
Chlamydiae bacterium 1095360-24
Parachlamydiales bacterium palsa_1444
Chlamydiae bacterium 1021942-13
Chlamydiae bacterium Ga0074140
Chlamydiae bacterium 1064120-29
Chlamydiae bacterium BWOrgHeat_Nextera88_MAG2
Chlamydiae bacterium 3300005529-103
Candidatus Rhabdochlamydia helveticae
Rhabdochlamydia oedothorax
Candidatus Rhabdochlamydia porcellionis
Chlamydiae bacterium 3300006056-139
Chlamydiae bacterium O_S4.2_102
Chlamydiae bacterium RIFCSPHIGHO2_12_FULL_49_11
Chlamydiae bacterium O_S4.2_20
Chlamydiae bacterium 3300013126-109
Chlamydiae bacterium O_S4.1_5
Chlamydiae bacterium O_S4.2_27
Chlamydiae bacterium P_S1_9
Chlamydiae bacterium O_S4.2_37
Chlamydiae bacterium 3300013132-109
Chlamydiae bacterium 3300013133-59
Chlamydiae bacterium 1063924-70
Chlamydiae bacterium O_S4.2_107
Chlamydiae bacterium P_S1_54
Chlamydiae bacterium O_S4.1_1
Chlamydiae bacterium 1045240-7
Simkania negevensis
Chlamydiae bacterium K1060_chlam_2
Chlamydiae bacterium 1049244-12
Chlamydiae bacterium 1045192-5
Chlamydiae bacterium 3300004085-10
Chlamydiae bacterium 3300006056-100
Syngnamydia medusae
Candidatus Neptunochlamydia vexilliferae
Sygnamydia xenoturbellae
0.3

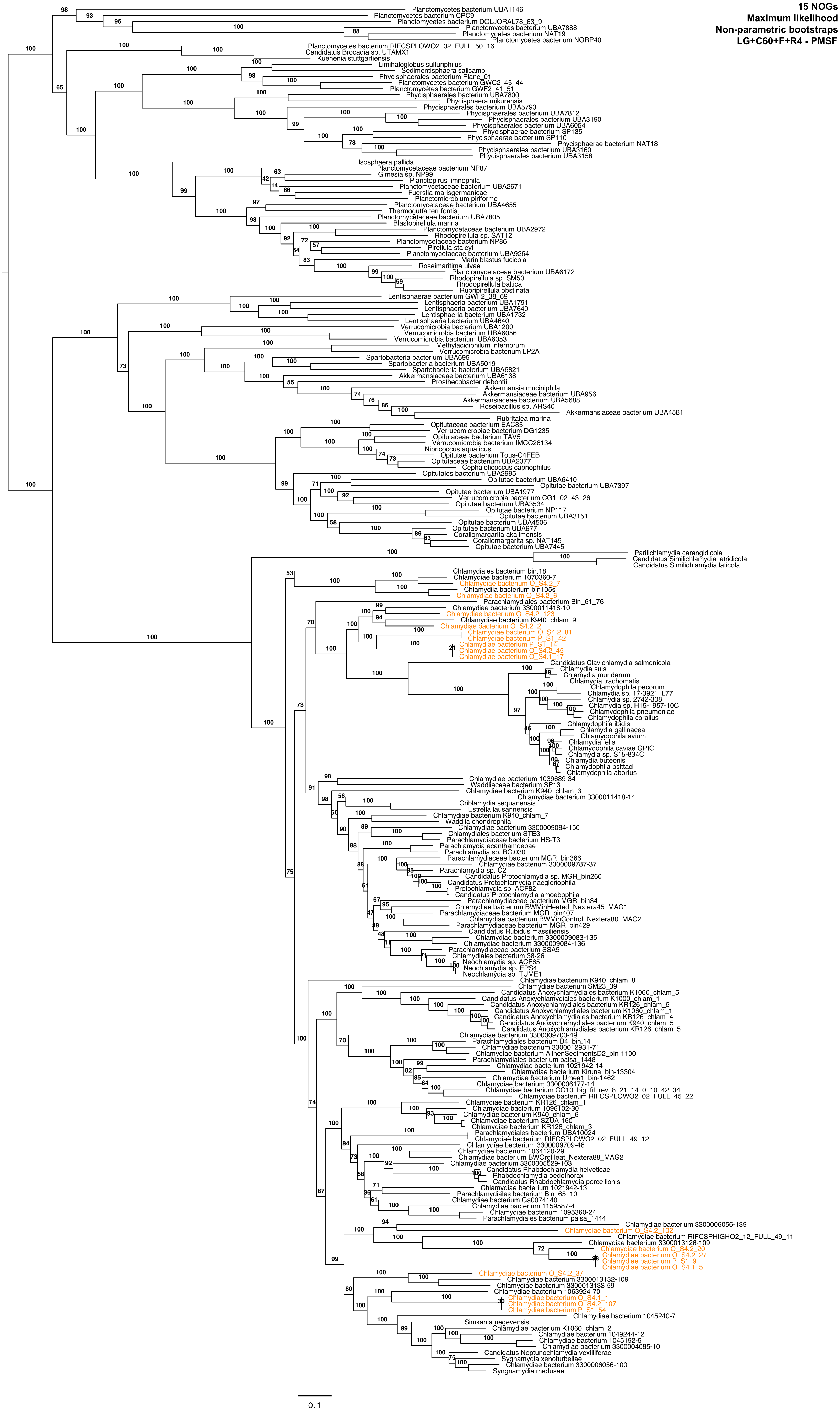
15 NOGs
Maximum likelihood
Non-parametric bootstraps
LG+C60+F+R4 - PMSF
Planctomycetes bacterium UBA1146
Planctomycetes bacterium CPC9
Planctomycetes bacterium DOLJORAL78_63_9
Planctomycetes bacterium UBA7888
Planctomycetes bacterium NAT19
Planctomycetes bacterium NORP40
Planctomycetes bacterium RIFCSPLOWO2_02_FULL_50_16
Candidatus Brocadia sp. UTAMX1
Kuenenia stuttgartiensis
Limihaloglobus sulfuriphilus
Sedimentisphaera salicampi
Phycisphaerales bacterium Planc_01
Planctomycetes bacterium GWC2_45_44
Planctomycetes bacterium GWF2_41_51
Phycisphaerales bacterium UBA7800
Phycisphaera mikurensis
Phycisphaerales bacterium UBA5793
Phycisphaerales bacterium UBA7812
Phycisphaerales bacterium UBA3190
Phycisphaerales bacterium UBA6054
Phycisphaerae bacterium SP135
Phycisphaerae bacterium SP110
Phycisphaerae bacterium NAT18
Phycisphaerales bacterium UBA3160
Phycisphaerales bacterium UBA3158
Isosphaera pallida
Planctomycetaceae bacterium NP87
Gimesia sp. NP99
Planctopirus limnophila
Planctomycetaceae bacterium UBA2671
Fuerstia marisgermanicae
Planctomicrobium piriforme
Planctomycetaceae bacterium UBA4655
Thermogutta terrifontis
Planctomycetaceae bacterium UBA7805
Blastopirellula marina
Planctomycetaceae bacterium UBA2972
Rhodopirellula sp. SAT12
Planctomycetaceae bacterium NP86
Pirellula staleyi
Planctomycetaceae bacterium UBA9264
Mariniblastus fucicola
Roseimaritima ulvae
Planctomycetaceae bacterium UBA6172
Rhodopirellula sp. SM50
Rhodopirellula baltica
Rubripirellula obstinata
Lentisphaerae bacterium GWF2_38_69
Lentisphaeria bacterium UBA1791
Lentisphaeria bacterium UBA7640
Lentisphaeria bacterium UBA1732
Lentisphaeria bacterium UBA4640
Verrucomicrobia bacterium UBA1200
Verrucomicrobia bacterium UBA6056
Verrucomicrobia bacterium UBA6053
Methylacidiphilum infernorum
Verrucomicrobia bacterium LP2A
Spartobacteria bacterium UBA695
Spartobacteria bacterium UBA5019
Spartobacteria bacterium UBA6821
Akkermansiaceae bacterium UBA6138
Prosthecobacter debontii
Akkermansia muciniphila
Akkermansiaceae bacterium UBA956
Akkermansiaceae bacterium UBA5688
Roseibacillus sp. ARS40
Akkermansiaceae bacterium UBA4581
Rubritalea marina
Opitutaceae bacterium EAC85
Verrucomicrobiae bacterium DG1235
Opitutaceae bacterium TAV5
Verrucomicrobia bacterium IMCC26134
Nibricoccus aquaticus
Opitutae bacterium Tous-C4FEB
Opitutaceae bacterium UBA2377
Cephaloticoccus capnophilus
Opitutales bacterium UBA2995
Opitutae bacterium UBA6410
Opitutae bacterium UBA7397
Opitutae bacterium UBA1977
Verrucomicrobia bacterium CG1_02_43_26
Opitutae bacterium UBA3534
Opitutae bacterium NP117
Opitutae bacterium UBA3151
Opitutae bacterium UBA4506
Opitutae bacterium UBA977
Coraliomargarita akajimensis
Coraliomargarita sp. NAT145
Opitutae bacterium UBA7445
Parilichlamydia carangidicola
Candidatus Similichlamydia latridicola
Candidatus Similichlamydia laticola
Chlamydiales bacterium bin.18
Chlamydiae bacterium 1070360-7
Chlamydiae bacterium O_S4.2_7
Chlamydiia bacterium bin105s
Chlamydiae bacterium O_S4.2_6
Parachlamydiales bacterium Bin_61_76
Chlamydiae bacterium 3300011418-10
Chlamydiae bacterium O_S4.2_123
Chlamydiae bacterium K940_chlam_9
Chlamydiae bacterium O_S4.2_2
Chlamydiae bacterium O_S4.2_81
Chlamydiae bacterium P_S1_42
Chlamydiae bacterium P_S1_14
Chlamydiae bacterium O_S4.2_45
Chlamydiae bacterium O_S4.1_17
Candidatus Clavichlamydia salmonicola
Chlamydia suis
Chlamydia muridarum
Chlamydia trachomatis
Chlamydophila pecorum
Chlamydia sp. 17-3921_L77
Chlamydia sp. 2742-308
Chlamydia sp. H15-1957-10C
Chlamydophila pneumoniae
Chlamydophila corallus
Chlamydophila ibidis
Chlamydia gallinacea
Chlamydophila avium
Chlamydia felis
Chlamydophila caviae GPIC
Chlamydia sp. S15-834C
Chlamydia buteonis
Chlamydophila psittaci
Chlamydophila abortus
Chlamydiae bacterium 1039689-34
Waddliaceae bacterium SP13
Chlamydiae bacterium K940_chlam_3
Chlamydiae bacterium 3300011418-14
Criblamydia sequanensis
Estrella lausannensis
Chlamydiae bacterium K940_chlam_7
Waddlia chondrophila
Chlamydiae bacterium 3300009084-150
Chlamydiales bacterium STE3
Parachlamydiaceae bacterium HS-T3
Parachlamydia acanthamoebae
Parachlamydia sp. BC.030
Parachlamydiaceae bacterium MGR_bin366
Chlamydiae bacterium 3300009787-37
Parachlamydia sp. C2
Candidatus Protochlamydia sp. MGR_bin260
Candidatus Protochlamydia naegleriophila
Protochlamydia sp. ACF82
Candidatus Protochlamydia amoebophila
Parachlamydiaceae bacterium MGR_bin34
Chlamydiae bacterium BWMinHeated_Nextera45_MAG1
Parachlamydiaceae bacterium MGR_bin407
Chlamydiae bacterium BWMinControl_Nextera80_MAG2
Parachlamydiaceae bacterium MGR_bin429
Candidatus Rubidus massiliensis
Chlamydiae bacterium 3300009083-135
Chlamydiae bacterium 3300009084-136
Parachlamydiaceae bacterium SSA5
Chlamydiales bacterium 38-26
Neochlamydia sp. ACF65
Neochlamydia sp. EPS4
Neochlamydia sp. TUME1
Chlamydiae bacterium K940_chlam_8
Chlamydiae bacterium SM23_39
Candidatus Anoxychlamydiales bacterium K1060_chlam_5
Candidatus Anoxychlamydiales bacterium K1000_chlam_1
Candidatus Anoxychlamydiales bacterium KR126_chlam_6
Candidatus Anoxychlamydiales bacterium K1060_chlam_1
Candidatus Anoxychlamydiales bacterium KR126_chlam_4
Candidatus Anoxychlamydiales bacterium K940_chlam_5
Candidatus Anoxychlamydiales bacterium KR126_chlam_5
Chlamydiae bacterium 3300009703-49
Parachlamydiales bacterium B4_bin.14
Chlamydiae bacterium 3300012931-71
Chlamydiae bacterium AlinenSedimentsD2_bin-1100
Parachlamydiales bacterium palsa_1448
Chlamydiae bacterium 1021942-14
Chlamydiae bacterium Kiruna_bin-13304
Chlamydiae bacterium Umea1_bin-1462
Chlamydiae bacterium 3300006177-14
Chlamydiae bacterium CG10_big_fil_rev_8_21_14_0_10_42_34
Chlamydiae bacterium RIFCSPLOWO2_02_FULL_45_22
Chlamydiae bacterium KR126_chlam_1
Chlamydiae bacterium 1096102-30
Chlamydiae bacterium K940_chlam_6
Chlamydiae bacterium SZUA-160
Chlamydiae bacterium KR126_chlam_3
Parachlamydiales bacterium UBA10024
Chlamydiae bacterium RIFCSPLOWO2_02_FULL_49_12
Chlamydiae bacterium 3300009703-46
Chlamydiae bacterium 1064120-29
Chlamydiae bacterium BWOrgHeat_Nextera88_MAG2
Chlamydiae bacterium 3300005529-103
Candidatus Rhabdochlamydia helveticae
Rhabdochlamydia oedothorax
Candidatus Rhabdochlamydia porcellionis
Chlamydiae bacterium 1021942-13
Parachlamydiales bacterium Bin_65_10
Chlamydiae bacterium Ga0074140
Chlamydiae bacterium 1159587-4
Chlamydiae bacterium 1095360-24
Parachlamydiales bacterium palsa_1444
Chlamydiae bacterium 3300006056-139
Chlamydiae bacterium O_S4.2_102
Chlamydiae bacterium RIFCSPHIGHO2_12_FULL_49_11
Chlamydiae bacterium 3300013126-109
Chlamydiae bacterium O_S4.2_20
Chlamydiae bacterium O_S4.2_27
Chlamydiae bacterium P_S1_9
Chlamydiae bacterium O_S4.1_5
Chlamydiae bacterium O_S4.2_37
Chlamydiae bacterium 3300013132-109
Chlamydiae bacterium 3300013133-59
Chlamydiae bacterium 1063924-70
Chlamydiae bacterium O_S4.1_1
Chlamydiae bacterium O_S4.2_107
Chlamydiae bacterium P_S1_54
Chlamydiae bacterium 1045240-7
Simkania negevensis
Chlamydiae bacterium K1060_chlam_2
Chlamydiae bacterium 1049244-12
Chlamydiae bacterium 1045192-5
Chlamydiae bacterium 3300004085-10
Candidatus Neptunochlamydia vexilliferae
Syngnamydia xenoturbellae
Chlamydiae bacterium 3300006056-100
Syngnamydia medusae
0.1

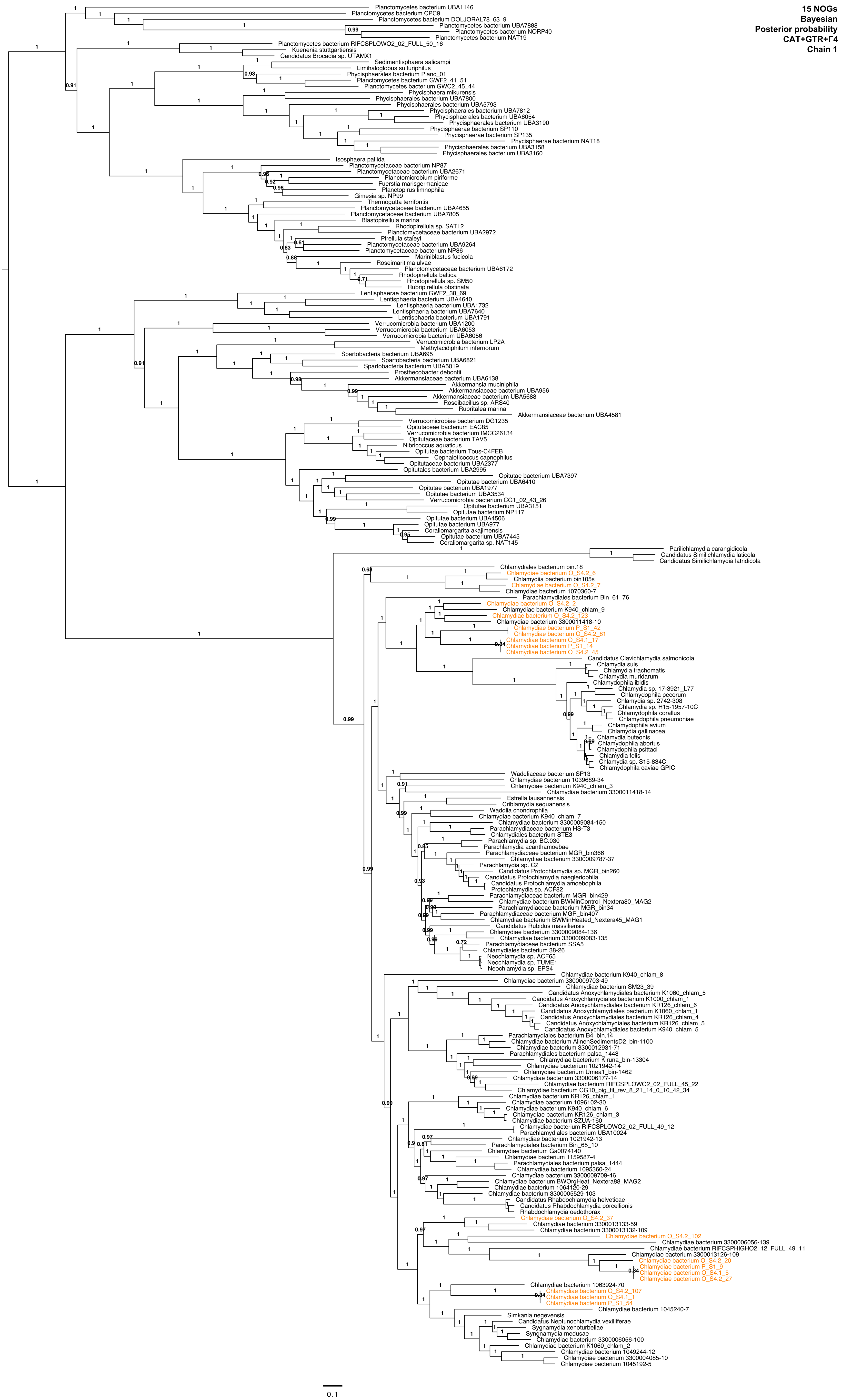
15 NOGs
Bayesian
Posterior probability
CAT+GTR+Γ4
Chain 1
Planctomycetes bacterium UBA1146
Planctomycetes bacterium CPC9
Planctomycetes bacterium DOLJORAL78_63_9
Planctomycetes bacterium UBA7888
Planctomycetes bacterium NORP40
Planctomycetes bacterium NAT19
Planctomycetes bacterium RIFCSPLOWO2_02_FULL_50_16
Kuenenia stuttgartiensis
Candidatus Brocadia sp. UTAMX1
Sedimentisphaera salicampi
Limihaloglobus sulfuriphilus
Phycisphaerales bacterium Planc_01
Planctomycetes bacterium GWF2_41_51
Planctomycetes bacterium GWC2_45_44
Phycisphaera mikurensis
Phycisphaerales bacterium UBA7800
Phycisphaerales bacterium UBA5793
Phycisphaerales bacterium UBA7812
Phycisphaerales bacterium UBA6054
Phycisphaerales bacterium UBA3190
Phycisphaerae bacterium SP110
Phycisphaerae bacterium SP135
Phycisphaerae bacterium NAT18
Phycisphaerales bacterium UBA3158
Phycisphaerales bacterium UBA3160
Isosphaera pallida
Planctomycetaceae bacterium NP87
Planctomycetaceae bacterium UBA2671
Planctomicrobium piriforme
Fuerstia marisgermanicae
Planctopirus limnophila
Gimesia sp. NP99
Thermogutta terrifontis
Planctomycetaceae bacterium UBA4655
Planctomycetaceae bacterium UBA7805
Blastopirellula marina
Rhodopirellula sp. SAT12
Planctomycetaceae bacterium UBA2972
Pirellula staleyi
Planctomycetaceae bacterium UBA9264
Planctomycetaceae bacterium NP86
Mariniblastus fucicola
Roseimaritima ulvae
Planctomycetaceae bacterium UBA6172
Rhodopirellula baltica
Rhodopirellula sp. SM50
Rubripirellula obstinata
Lentisphaerae bacterium GWF2_38_69
Lentisphaeria bacterium UBA4640
Lentisphaeria bacterium UBA1732
Lentisphaeria bacterium UBA7640
Lentisphaeria bacterium UBA1791
Verrucomicrobia bacterium UBA1200
Verrucomicrobia bacterium UBA6053
Verrucomicrobia bacterium UBA6056
Verrucomicrobia bacterium LP2A
Methylacidiphilum infernorum
Spartobacteria bacterium UBA695
Spartobacteria bacterium UBA6821
Spartobacteria bacterium UBA5019
Prosthecobacter debontii
Akkermansiaceae bacterium UBA6138
Akkermansia muciniphila
Akkermansiaceae bacterium UBA956
Akkermansiaceae bacterium UBA5688
Roseibacillus sp. ARS40
Rubritalea marina
Akkermansiaceae bacterium UBA4581
Verrucomicrobiae bacterium DG1235
Opitutaceae bacterium EAC85
Verrucomicrobia bacterium IMCC26134
Opitutaceae bacterium TAV5
Nibricoccus aquaticus
Opitutae bacterium Tous-C4FEB
Cephaloticoccus capnophilus
Opitutaceae bacterium UBA2377
Opitutales bacterium UBA2995
Opitutae bacterium UBA7397
Opitutae bacterium UBA6410
Opitutae bacterium UBA1977
Opitutae bacterium UBA3534
Verrucomicrobia bacterium CG1_02_43_26
Opitutae bacterium UBA3151
Opitutae bacterium NP117
Opitutae bacterium UBA4506
Opitutae bacterium UBA977
Coraliomargarita akajimensis
Opitutae bacterium UBA7445
Coraliomargarita sp. NAT145
Parilichlamydia carangidicola
Candidatus Similichlamydia laticola
Candidatus Similichlamydia latridicola
Chlamydiales bacterium bin.18
Chlamydiae bacterium O_S4.2_6
Chlamydia bacterium bin105s
Chlamydiae bacterium O_S4.2_7
Chlamydiae bacterium 1070360-7
Parachlamydiales bacterium Bin_61_76
Chlamydiae bacterium O_S4.2_2
Chlamydiae bacterium K940_chlam_9
Chlamydiae bacterium O_S4.2_123
Chlamydiae bacterium 3300011418-10
Chlamydiae bacterium P_S1_42
Chlamydiae bacterium O_S4.2_81
Chlamydiae bacterium O_S4.1_17
Chlamydiae bacterium P_S1_14
Chlamydiae bacterium O_S4.2_45
Candidatus Clavichlamydia salmonicola
Chlamydia suis
Chlamydia trachomatis
Chlamydia muridarum
Chlamydophila ibidis
Chlamydia sp. 17-3921_L77
Chlamydophila pecorum
Chlamydia sp. 2742-308
Chlamydia sp. H15-1957-10C
Chlamydophila corallus
Chlamydophila pneumoniae
Chlamydophila avium
Chlamydia gallinacea
Chlamydia buteonis
Chlamydophila abortus
Chlamydophila psittaci
Chlamydia felis
Chlamydia sp. S15-834C
Chlamydophila caviae GPIC
Waddliaceae bacterium SP13
Chlamydiae bacterium 1039689-34
Chlamydiae bacterium K940_chlam_3
Chlamydiae bacterium 3300011418-14
Estrella lausannensis
Criblamydia sequanensis
Waddlia chondrophila
Chlamydiae bacterium K940_chlam_7
Chlamydiae bacterium 3300009084-150
Parachlamydiaceae bacterium HS-T3
Chlamydiales bacterium STE3
Parachlamydia sp. BC.030
Parachlamydia acanthamoebae
Parachlamydiaceae bacterium MGR_bin366
Chlamydiae bacterium 3300009787-37
Parachlamydia sp. C2
Candidatus Protochlamydia sp. MGR_bin260
Candidatus Protochlamydia naegleriophila
Candidatus Protochlamydia amoebophila
Protochlamydia sp. ACF82
Parachlamydiaceae bacterium MGR_bin429
Chlamydiae bacterium BWMinControl_Nextera80_MAG2
Parachlamydiaceae bacterium MGR_bin34
Parachlamydiaceae bacterium MGR_bin407
Chlamydiae bacterium BWMinHeated_Nextera45_MAG1
Candidatus Rubidus massiliensis
Chlamydiae bacterium 3300009084-136
Chlamydiae bacterium 3300009083-135
Parachlamydiaceae bacterium SSA5
Chlamydiales bacterium 38-26
Neochlamydia sp. ACF65
Neochlamydia sp. TUME1
Neochlamydia sp. EPS4
Chlamydiae bacterium K940_chlam_8
Chlamydiae bacterium 3300009703-49
Chlamydiae bacterium SM23_39
Candidatus Anoxychlamydiales bacterium K1060_chlam_5
Candidatus Anoxychlamydiales bacterium K1000_chlam_1
Candidatus Anoxychlamydiales bacterium KR126_chlam_6
Candidatus Anoxychlamydiales bacterium K1060_chlam_1
Candidatus Anoxychlamydiales bacterium KR126_chlam_4
Candidatus Anoxychlamydiales bacterium KR126_chlam_5
Candidatus Anoxychlamydiales bacterium K940_chlam_5
Parachlamydiales bacterium B4_bin.14
Chlamydiae bacterium AlinenSedimentsD2_bin-1100
Chlamydiae bacterium 3300012931-71
Parachlamydiales bacterium palsa_1448
Chlamydiae bacterium Kiruna_bin-13304
Chlamydiae bacterium 1021942-14
Chlamydiae bacterium Umea1_bin-1462
Chlamydiae bacterium 3300006177-14
Chlamydiae bacterium RIFCSPLOWO2_02_FULL_45_22
Chlamydiae bacterium CG10_big_fil_rev_8_21_14_0_10_42_34
Chlamydiae bacterium KR126_chlam_1
Chlamydiae bacterium 1096102-30
Chlamydiae bacterium K940_chlam_6
Chlamydiae bacterium KR126_chlam_3
Chlamydiae bacterium SZUA-160
Chlamydiae bacterium RIFCSPLOWO2_02_FULL_49_12
Parachlamydiales bacterium UBA10024
Chlamydiae bacterium 1021942-13
Parachlamydiales bacterium Bin_65_10
Chlamydiae bacterium Ga0074140
Chlamydiae bacterium 1159587-4
Parachlamydiales bacterium palsa_1444
Chlamydiae bacterium 1095360-24
Chlamydiae bacterium 3300009709-46
Chlamydiae bacterium BWOrgHeat_Nextera88_MAG2
Chlamydiae bacterium 1064120-29
Chlamydiae bacterium 3300005529-103
Candidatus Rhabdochlamydia helveticae
Candidatus Rhabdochlamydia porcellionis
Rhabdochlamydia oedothorax
Chlamydiae bacterium O_S4.2_37
Chlamydiae bacterium 3300013133-59
Chlamydiae bacterium 3300013132-109
Chlamydiae bacterium O_S4.2_102
Chlamydiae bacterium 3300006056-139
Chlamydiae bacterium RIFCSPHIGHO2_12_FULL_49_11
Chlamydiae bacterium 3300013126-109
Chlamydiae bacterium O_S4.2_20
Chlamydiae bacterium P_S1_9
Chlamydiae bacterium O_S4.1_5
Chlamydiae bacterium O_S4.2_27
Chlamydiae bacterium 1063924-70
Chlamydiae bacterium O_S4.2_107
Chlamydiae bacterium O_S4.1_1
Chlamydiae bacterium P_S1_54
Chlamydiae bacterium 1045240-7
Simkania negevensis
Candidatus Neptunochlamydia vexilliferae
Sygnamydia xenoturbellae
Syngnamydia medusae
Chlamydiae bacterium 3300006056-100
Chlamydiae bacterium K1060_chlam_2
Chlamydiae bacterium 1049244-12
Chlamydiae bacterium 3300004085-10
Chlamydiae bacterium 1045192-5
0.1

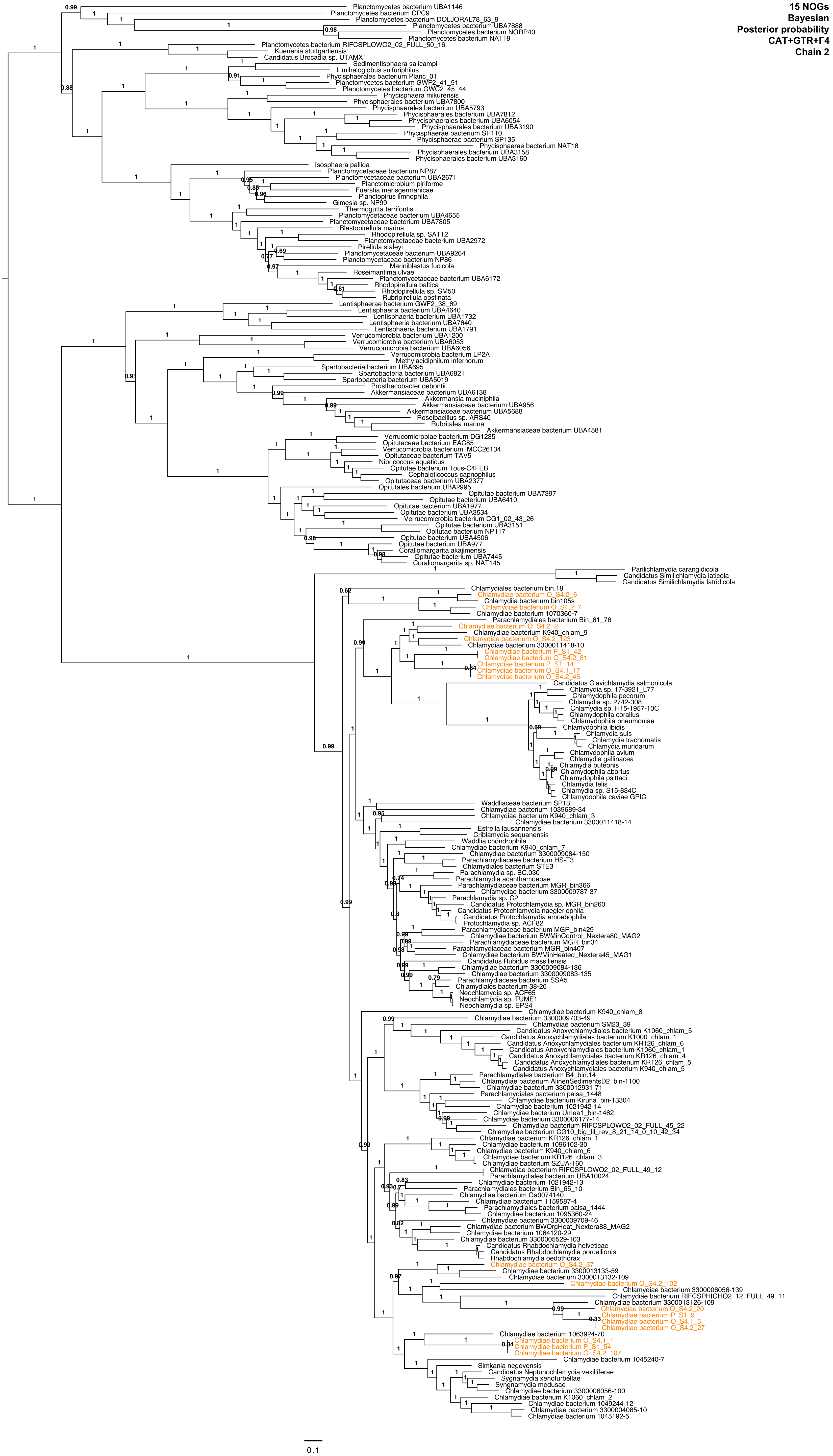
15 NOGs
Bayesian
Posterior probability
CAT+GTR+Γ4
Chain 2
Planctomycetes bacterium UBA1146
Planctomycetes bacterium CPC9
Planctomycetes bacterium DOLJORAL78_63_9
Planctomycetes bacterium UBA7888
Planctomycetes bacterium NORP40
Planctomycetes bacterium NAT19
Planctomycetes bacterium RIFCSPLOWO2_02_FULL_50_16
Kuenenia stuttgartiensis
Candidatus Brocadia sp. UTAMX1
Sedimentisphaera salicampi
Limihaloglobus sulfuriphilus
Phycisphaerales bacterium Planc_01
Planctomycetes bacterium GWF2_41_51
Planctomycetes bacterium GWC2_45_44
Phycisphaera mikurensis
Phycisphaerales bacterium UBA7800
Phycisphaerales bacterium UBA5793
Phycisphaerales bacterium UBA7812
Phycisphaerales bacterium UBA6054
Phycisphaerales bacterium UBA3190
Phycisphaerae bacterium SP110
Phycisphaerae bacterium SP135
Phycisphaerae bacterium NAT18
Phycisphaerales bacterium UBA3158
Phycisphaerales bacterium UBA3160
Isosphaera pallida
Planctomycetaceae bacterium NP87
Planctomycetaceae bacterium UBA2671
Planctomicrobium piriforme
Fuerstia marisgermanicae
Planctopirus limnophila
Gimesia sp. NP99
Thermogutta terrifontis
Planctomycetaceae bacterium UBA4655
Planctomycetaceae bacterium UBA7805
Blastopirellula marina
Rhodopirellula sp. SAT12
Planctomycetaceae bacterium UBA2972
Pirellula staleyi
Planctomycetaceae bacterium UBA9264
Planctomycetaceae bacterium NP86
Mariniblastus fucicola
Roseimaritima ulvae
Planctomycetaceae bacterium UBA6172
Rhodopirellula baltica
Rhodopirellula sp. SM50
Rubripirellula obstinata
Lentisphaerae bacterium GWF2_38_69
Lentisphaeria bacterium UBA4640
Lentisphaeria bacterium UBA1732
Lentisphaeria bacterium UBA7640
Lentisphaeria bacterium UBA1791
Verrucomicrobia bacterium UBA1200
Verrucomicrobia bacterium UBA6053
Verrucomicrobia bacterium UBA6056
Verrucomicrobia bacterium LP2A
Methylacidiphilum infernorum
Spartobacteria bacterium UBA695
Spartobacteria bacterium UBA6821
Spartobacteria bacterium UBA5019
Prosthecobacter debontii
Akkermansiaceae bacterium UBA6138
Akkermansia muciniphila
Akkermansiaceae bacterium UBA956
Akkermansiaceae bacterium UBA5688
Roseibacillus sp. ARS40
Rubritalea marina
Akkermansiaceae bacterium UBA4581
Verrucomicrobiae bacterium DG1235
Opitutaceae bacterium EAC85
Verrucomicrobia bacterium IMCC26134
Opitutaceae bacterium TAV5
Nibricoccus aquaticus
Opitutae bacterium Tous-C4FEB
Cephaloticoccus capnophilus
Opitutaceae bacterium UBA2377
Opitutales bacterium UBA2995
Opitutae bacterium UBA7397
Opitutae bacterium UBA6410
Opitutae bacterium UBA1977
Opitutae bacterium UBA3534
Verrucomicrobia bacterium CG1_02_43_26
Opitutae bacterium UBA3151
Opitutae bacterium NP117
Opitutae bacterium UBA4506
Opitutae bacterium UBA977
Coraliomargarita akajimensis
Opitutae bacterium UBA7445
Coraliomargarita sp. NAT145
Parilichlamydia carangidicola
Candidatus Similichlamydia laticola
Candidatus Similichlamydia latridicola
Chlamydiales bacterium bin.18
Chlamydiae bacterium O_S4.2_6
Chlamydia bacterium bin105s
Chlamydiae bacterium O_S4.2_7
Chlamydiae bacterium 1070360-7
Parachlamydiales bacterium Bin_61_76
Chlamydiae bacterium O_S4.2_2
Chlamydiae bacterium K940_chlam_9
Chlamydiae bacterium O_S4.2_123
Chlamydiae bacterium 3300011418-10
Chlamydiae bacterium P_S1_42
Chlamydiae bacterium O_S4.2_81
Chlamydiae bacterium P_S1_14
Chlamydiae bacterium O_S4.1_17
Chlamydiae bacterium O_S4.2_45
Candidatus Clavichlamydia salmonicola
Chlamydia sp. 17-3921_L77
Chlamydophila pecorum
Chlamydia sp. 2742-308
Chlamydia sp. H15-1957-10C
Chlamydophila corallus
Chlamydophila pneumoniae
Chlamydophila ibidis
Chlamydia suis
Chlamydia trachomatis
Chlamydia muridarum
Chlamydophila avium
Chlamydia gallinacea
Chlamydia buteonis
Chlamydophila abortus
Chlamydophila psittaci
Chlamydia felis
Chlamydia sp. S15-834C
Chlamydophila caviae GPIC
Waddliaceae bacterium SP13
Chlamydiae bacterium 1039689-34
Chlamydiae bacterium K940_chlam_3
Chlamydiae bacterium 3300011418-14
Estrella lausannensis
Criblamydia sequanensis
Waddlia chondrophila
Chlamydiae bacterium K940_chlam_7
Chlamydiae bacterium 3300009084-150
Parachlamydiaceae bacterium HS-T3
Chlamydiales bacterium STE3
Parachlamydia sp. BC.030
Parachlamydia acanthamoebae
Parachlamydiaceae bacterium MGR_bin366
Chlamydiae bacterium 3300009787-37
Parachlamydia sp. C2
Candidatus Protochlamydia sp. MGR_bin260
Candidatus Protochlamydia naegleriophila
Candidatus Protochlamydia amoebophila
Protochlamydia sp. ACF82
Parachlamydiaceae bacterium MGR_bin429
Chlamydiae bacterium BWMinControl_Nextera80_MAG2
Parachlamydiaceae bacterium MGR_bin34
Parachlamydiaceae bacterium MGR_bin407
Chlamydiae bacterium BWMinHeated_Nextera45_MAG1
Candidatus Rubidus massiliensis
Chlamydiae bacterium 3300009084-136
Chlamydiae bacterium 3300009083-135
Parachlamydiaceae bacterium SSA5
Chlamydiales bacterium 38-26
Neochlamydia sp. ACF65
Neochlamydia sp. TUME1
Neochlamydia sp. EPS4
Chlamydiae bacterium K940_chlam_8
Chlamydiae bacterium 3300009703-49
Chlamydiae bacterium SM23_39
Candidatus Anoxychlamydiales bacterium K1060_chlam_5
Candidatus Anoxychlamydiales bacterium K1000_chlam_1
Candidatus Anoxychlamydiales bacterium KR126_chlam_6
Candidatus Anoxychlamydiales bacterium K1060_chlam_1
Candidatus Anoxychlamydiales bacterium KR126_chlam_4
Candidatus Anoxychlamydiales bacterium KR126_chlam_5
Candidatus Anoxychlamydiales bacterium K940_chlam_5
Parachlamydiales bacterium B4_bin.14
Chlamydiae bacterium AlinenSedimentsD2_bin-1100
Chlamydiae bacterium 3300012931-71
Parachlamydiales bacterium palsa_1448
Chlamydiae bacterium Kiruna_bin-13304
Chlamydiae bacterium 1021942-14
Chlamydiae bacterium Umea1_bin-1462
Chlamydiae bacterium 3300006177-14
Chlamydiae bacterium RIFCSPLOWO2_02_FULL_45_22
Chlamydiae bacterium CG10_big_fil_rev_8_21_14_0_10_42_34
Chlamydiae bacterium KR126_chlam_1
Chlamydiae bacterium 1096102-30
Chlamydiae bacterium K940_chlam_6
Chlamydiae bacterium KR126_chlam_3
Chlamydiae bacterium SZUA-160
Chlamydiae bacterium RIFCSPLOWO2_02_FULL_49_12
Parachlamydiales bacterium UBA10024
Chlamydiae bacterium 1021942-13
Parachlamydiales bacterium Bin_65_10
Chlamydiae bacterium Ga0074140
Chlamydiae bacterium 1159587-4
Parachlamydiales bacterium palsa_1444
Chlamydiae bacterium 1095360-24
Chlamydiae bacterium 3300009709-46
Chlamydiae bacterium BWOrgHeat_Nextera88_MAG2
Chlamydiae bacterium 1064120-29
Chlamydiae bacterium 3300005529-103
Candidatus Rhabdochlamydia helveticae
Candidatus Rhabdochlamydia porcellionis
Rhabdochlamydia oedothorax
Chlamydiae bacterium O_S4.2_37
Chlamydiae bacterium 3300013133-59
Chlamydiae bacterium 3300013132-109
Chlamydiae bacterium O_S4.2_102
Chlamydiae bacterium 3300006056-139
Chlamydiae bacterium RIFCSPHIGHO2_12_FULL_49_11
Chlamydiae bacterium 3300013126-109
Chlamydiae bacterium O_S4.2_20
Chlamydiae bacterium P_S1_9
Chlamydiae bacterium O_S4.1_5
Chlamydiae bacterium O_S4.2_27
Chlamydiae bacterium 1063924-70
Chlamydiae bacterium O_S4.1_1
Chlamydiae bacterium P_S1_54
Chlamydiae bacterium O_S4.2_107
Chlamydiae bacterium 1045240-7
Simkania negevensis
Candidatus Neptunochlamydia vexilliferae
Sygnamydia xenoturbellae
Syngnamydia medusae
Chlamydiae bacterium 3300006056-100
Chlamydiae bacterium K1060_chlam_2
Chlamydiae bacterium 1049244-12
Chlamydiae bacterium 3300004085-10
Chlamydiae bacterium 1045192-5
0.1

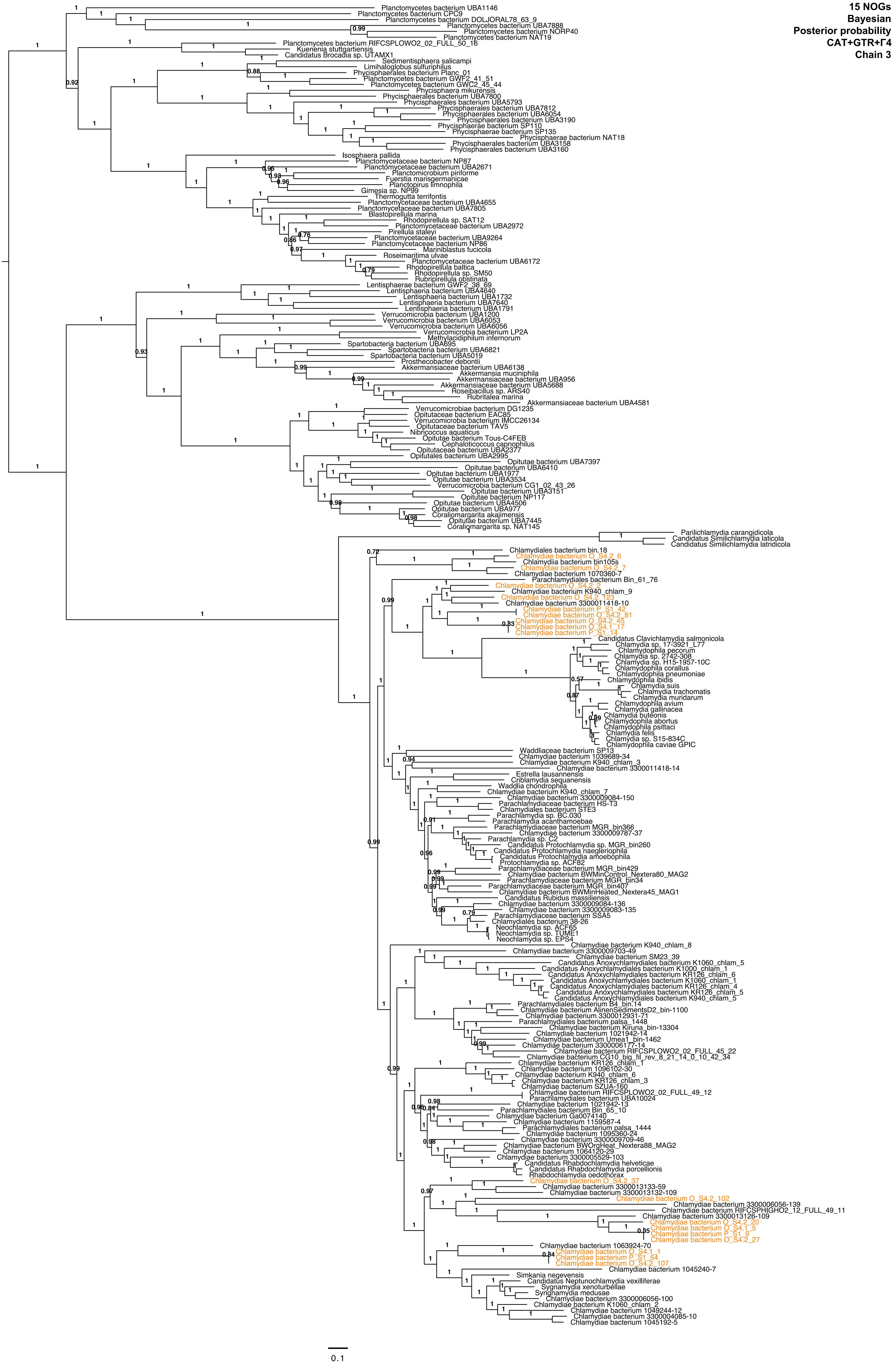

15 NOGs
Bayesian
Posterior probability
CAT+GTR+Γ4
Chain 3
Planctomycetes bacterium UBA1146
Planctomycetes bacterium CPC9
Planctomycetes bacterium DOLJORAL78_63_9
Planctomycetes bacterium UBA7888
Planctomycetes bacterium NORP40
Planctomycetes bacterium NAT19
Planctomycetes bacterium RIFCSPLOWO2_02_FULL_50_16
Kuenenia stuttgartiensis
Candidatus Brocadia sp. UTAMX1
Sedimentisphaera salicampi
Limihaloglobus sulfuriphilus
Phycisphaerales bacterium Planc_01
Planctomycetes bacterium GWF2_41_51
Planctomycetes bacterium GWC2_45_44
Phycisphaera mikurensis
Phycisphaerales bacterium UBA7800
Phycisphaerales bacterium UBA5793
Phycisphaerales bacterium UBA7812
Phycisphaerales bacterium UBA6054
Phycisphaerales bacterium UBA3190
Phycisphaerae bacterium SP110
Phycisphaerae bacterium SP135
Phycisphaerae bacterium NAT18
Phycisphaerales bacterium UBA3158
Phycisphaerales bacterium UBA3160
Isosphaera pallida
Planctomycetaceae bacterium NP87
Planctomycetaceae bacterium UBA2671
Planctomicrobium piriforme
Fuerstia marisgermanicae
Planctopirus limnophila
Gimesia sp. NP99
Thermogutta terrifontis
Planctomycetaceae bacterium UBA4655
Planctomycetaceae bacterium UBA7805
Blastopirellula marina
Rhodopirellula sp. SAT12
Planctomycetaceae bacterium UBA2972
Pirellula staleyi
Planctomycetaceae bacterium UBA9264
Planctomycetaceae bacterium NP86
Mariniblastus fucicola
Roseimaritima ulvae
Planctomycetaceae bacterium UBA6172
Rhodopirellula baltica
Rhodopirellula sp. SM50
Rubripirellula obstinata
Lentisphaerae bacterium GWF2_38_69
Lentisphaeria bacterium UBA4640
Lentisphaeria bacterium UBA1732
Lentisphaeria bacterium UBA7640
Lentisphaeria bacterium UBA1791
Verrucomicrobia bacterium UBA1200
Verrucomicrobia bacterium UBA6053
Verrucomicrobia bacterium UBA6056
Verrucomicrobia bacterium LP2A
Methylacidiphilum infernorum
Spartobacteria bacterium UBA695
Spartobacteria bacterium UBA6821
Spartobacteria bacterium UBA5019
Prosthecobacter debontii
Akkermansiaceae bacterium UBA6138
Akkermansia muciniphila
Akkermansiaceae bacterium UBA956
Akkermansiaceae bacterium UBA5688
Roseibacillus sp. ARS40
Rubritalea marina
Akkermansiaceae bacterium UBA4581
Verrucomicrobiae bacterium DG1235
Opitutaceae bacterium EAC85
Verrucomicrobia bacterium IMCC26134
Opitutaceae bacterium TAV5
Nibricoccus aquaticus
Opitutae bacterium Tous-C4FEB
Cephaloticoccus capnophilus
Opitutaceae bacterium UBA2377
Opitutales bacterium UBA2995
Opitutae bacterium UBA7397
Opitutae bacterium UBA6410
Opitutae bacterium UBA1977
Opitutae bacterium UBA3534
Verrucomicrobia bacterium CG1_02_43_26
Opitutae bacterium UBA3151
Opitutae bacterium NP117
Opitutae bacterium UBA4506
Opitutae bacterium UBA977
Coraliomargarita akajimensis
Opitutae bacterium UBA7445
Coraliomargarita sp. NAT145
Parilichlamydia carangidicola
Candidatus Similichlamydia laticola
Candidatus Similichlamydia latridicola
Chlamydiales bacterium bin.18
Chlamydiae bacterium O_S4.2_6
Chlamydia bacterium bin105s
Chlamydiae bacterium O_S4.2_7
Chlamydiae bacterium 1070360-7
Parachlamydiales bacterium Bin_61_76
Chlamydiae bacterium O_S4.2_2
Chlamydiae bacterium K940_chlam_9
Chlamydiae bacterium O_S4.2_123
Chlamydiae bacterium 3300011418-10
Chlamydiae bacterium P_S1_42
Chlamydiae bacterium O_S4.2_81
Chlamydiae bacterium O_S4.2_45
Chlamydiae bacterium O_S4.1_17
Chlamydiae bacterium P_S1_14
Candidatus Clavichlamydia salmonicola
Chlamydia sp. 17-3921_L77
Chlamydophila pecorum
Chlamydia sp. 2742-308
Chlamydia sp. H15-1957-10C
Chlamydophila corallus
Chlamydophila pneumoniae
Chlamydophila ibidis
Chlamydia suis
Chlamydia trachomatis
Chlamydia muridarum
Chlamydophila avium
Chlamydia gallinacea
Chlamydia buteonis
Chlamydophila abortus
Chlamydophila psittaci
Chlamydia felis
Chlamydia sp. S15-834C
Chlamydophila caviae GPIC
Waddliaceae bacterium SP13
Chlamydiae bacterium 1039689-34
Chlamydiae bacterium K940_chlam_3
Chlamydiae bacterium 3300011418-14
Estrella lausannensis
Criblamydia sequanensis
Waddlia chondrophila
Chlamydiae bacterium K940_chlam_7
Chlamydiae bacterium 3300009084-150
Parachlamydiaceae bacterium HS-T3
Chlamydiales bacterium STE3
Parachlamydia sp. BC.030
Parachlamydia acanthamoebae
Parachlamydiaceae bacterium MGR_bin366
Chlamydiae bacterium 3300009787-37
Parachlamydia sp. C2
Candidatus Protochlamydia sp. MGR_bin260
Candidatus Protochlamydia naegleriophila
Candidatus Protochlamydia amoebophila
Protochlamydia sp. ACF82
Parachlamydiaceae bacterium MGR_bin429
Chlamydiae bacterium BWMinControl_Nextera80_MAG2
Parachlamydiaceae bacterium MGR_bin34
Parachlamydiaceae bacterium MGR_bin407
Chlamydiae bacterium BWMinHeated_Nextera45_MAG1
Candidatus Rubidus massiliensis
Chlamydiae bacterium 3300009084-136
Chlamydiae bacterium 3300009083-135
Parachlamydiaceae bacterium SSA5
Chlamydiales bacterium 38-26
Neochlamydia sp. ACF65
Neochlamydia sp. TUME1
Neochlamydia sp. EPS4
Chlamydiae bacterium K940_chlam_8
Chlamydiae bacterium 3300009703-49
Chlamydiae bacterium SM23_39
Candidatus Anoxychlamydiales bacterium K1060_chlam_5
Candidatus Anoxychlamydiales bacterium K1000_chlam_1
Candidatus Anoxychlamydiales bacterium KR126_chlam_6
Candidatus Anoxychlamydiales bacterium K1060_chlam_1
Candidatus Anoxychlamydiales bacterium KR126_chlam_4
Candidatus Anoxychlamydiales bacterium KR126_chlam_5
Candidatus Anoxychlamydiales bacterium K940_chlam_5
Parachlamydiales bacterium B4_bin.14
Chlamydiae bacterium AlinenSedimentsD2_bin-1100
Chlamydiae bacterium 3300012931-71
Parachlamydiales bacterium palsa_1448
Chlamydiae bacterium Kiruna_bin-13304
Chlamydiae bacterium 1021942-14
Chlamydiae bacterium Umea1_bin-1462
Chlamydiae bacterium 3300006177-14
Chlamydiae bacterium RIFCSPLOWO2_02_FULL_45_22
Chlamydiae bacterium CG10_big_fil_rev_8_21_14_0_10_42_34
Chlamydiae bacterium KR126_chlam_1
Chlamydiae bacterium 1096102-30
Chlamydiae bacterium K940_chlam_6
Chlamydiae bacterium KR126_chlam_3
Chlamydiae bacterium SZUA-160
Chlamydiae bacterium RIFCSPLOWO2_02_FULL_49_12
Parachlamydiales bacterium UBA10024
Chlamydiae bacterium 1021942-13
Parachlamydiales bacterium Bin_65_10
Chlamydiae bacterium Ga0074140
Chlamydiae bacterium 1159587-4
Parachlamydiales bacterium palsa_1444
Chlamydiae bacterium 1095360-24
Chlamydiae bacterium 3300009709-46
Chlamydiae bacterium BWOrgHeat_Nextera88_MAG2
Chlamydiae bacterium 1064120-29
Chlamydiae bacterium 3300005529-103
Candidatus Rhabdochlamydia helveticae
Candidatus Rhabdochlamydia porcellionis
Rhabdochlamydia oedothorax
Chlamydiae bacterium O_S4.2_37
Chlamydiae bacterium 3300013133-59
Chlamydiae bacterium 3300013132-109
Chlamydiae bacterium O_S4.2_102
Chlamydiae bacterium 3300006056-139
Chlamydiae bacterium RIFCSPHIGHO2_12_FULL_49_11
Chlamydiae bacterium 3300013126-109
Chlamydiae bacterium O_S4.2_20
Chlamydiae bacterium O_S4.1_5
Chlamydiae bacterium P_S1_9
Chlamydiae bacterium O_S4.2_27
Chlamydiae bacterium 1063924-70
Chlamydiae bacterium O_S4.1_1
Chlamydiae bacterium P_S1_54
Chlamydiae bacterium O_S4.2_107
Chlamydiae bacterium 1045240-7
Simkania negevensis
Candidatus Neptunochlamydia vexilliferae
Sygnamydia xenoturbellae
Syngnamydia medusae
Chlamydiae bacterium 3300006056-100
Chlamydiae bacterium K1060_chlam_2
Chlamydiae bacterium 1049244-12
Chlamydiae bacterium 3300004085-10
Chlamydiae bacterium 1045192-5
0.1

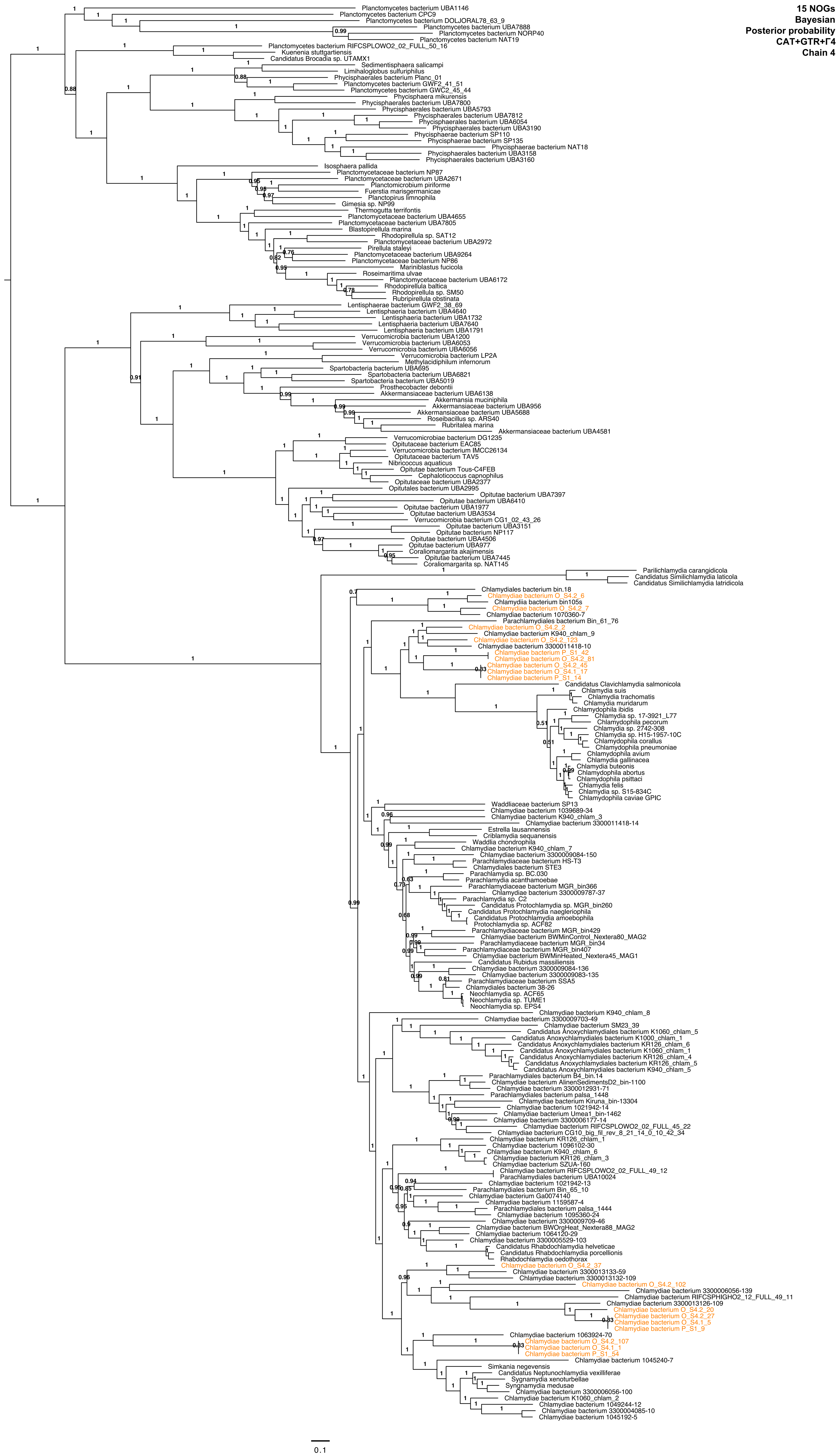
15 NOGs
Bayesian
Posterior probability
CAT+GTR+Γ4
Chain 4
Planctomycetes bacterium UBA1146
Planctomycetes bacterium CPC9
Planctomycetes bacterium DOLJORAL78_63_9
Planctomycetes bacterium UBA7888
Planctomycetes bacterium NORP40
Planctomycetes bacterium NAT19
Planctomycetes bacterium RIFCSPLOWO2_02_FULL_50_16
Kuenenia stuttgartiensis
Candidatus Brocadia sp. UTAMX1
Sedimentisphaera salicampi
Limihaloglobus sulfuriphilus
Phycisphaerales bacterium Planc_01
Planctomycetes bacterium GWF2_41_51
Planctomycetes bacterium GWC2_45_44
Phycisphaera mikurensis
Phycisphaerales bacterium UBA7800
Phycisphaerales bacterium UBA5793
Phycisphaerales bacterium UBA7812
Phycisphaerales bacterium UBA6054
Phycisphaerales bacterium UBA3190
Phycisphaerae bacterium SP110
Phycisphaerae bacterium SP135
Phycisphaerae bacterium NAT18
Phycisphaerales bacterium UBA3158
Phycisphaerales bacterium UBA3160
Isosphaera pallida
Planctomycetaceae bacterium NP87
Planctomycetaceae bacterium UBA2671
Planctomicrobium piriforme
Fuerstia marisgermanicae
Planctopirus limnophila
Gimesia sp. NP99
Thermogutta terrifontis
Planctomycetaceae bacterium UBA4655
Planctomycetaceae bacterium UBA7805
Blastopirellula marina
Rhodopirellula sp. SAT12
Planctomycetaceae bacterium UBA2972
Pirellula staleyi
Planctomycetaceae bacterium UBA9264
Planctomycetaceae bacterium NP86
Mariniblastus fucicola
Roseimaritima ulvae
Planctomycetaceae bacterium UBA6172
Rhodopirellula baltica
Rhodopirellula sp. SM50
Rubripirellula obstinata
Lentisphaerae bacterium GWF2_38_69
Lentisphaeria bacterium UBA4640
Lentisphaeria bacterium UBA1732
Lentisphaeria bacterium UBA7640
Lentisphaeria bacterium UBA1791
Verrucomicrobia bacterium UBA1200
Verrucomicrobia bacterium UBA6053
Verrucomicrobia bacterium UBA6056
Verrucomicrobia bacterium LP2A
Methylacidiphilum infernorum
Spartobacteria bacterium UBA695
Spartobacteria bacterium UBA6821
Spartobacteria bacterium UBA5019
Prosthecobacter debontii
Akkermansiaceae bacterium UBA6138
Akkermansia muciniphila
Akkermansiaceae bacterium UBA956
Akkermansiaceae bacterium UBA5688
Roseibacillus sp. ARS40
Rubritalea marina
Akkermansiaceae bacterium UBA4581
Verrucomicrobiae bacterium DG1235
Opitutaceae bacterium EAC85
Verrucomicrobia bacterium IMCC26134
Opitutaceae bacterium TAV5
Nibricoccus aquaticus
Opitutae bacterium Tous-C4FEB
Cephaloticoccus capnophilus
Opitutaceae bacterium UBA2377
Opitutales bacterium UBA2995
Opitutae bacterium UBA7397
Opitutae bacterium UBA6410
Opitutae bacterium UBA1977
Opitutae bacterium UBA3534
Verrucomicrobia bacterium CG1_02_43_26
Opitutae bacterium UBA3151
Opitutae bacterium NP117
Opitutae bacterium UBA4506
Opitutae bacterium UBA977
Coraliomargarita akajimensis
Opitutae bacterium UBA7445
Coraliomargarita sp. NAT145
Parilichlamydia carangidicola
Candidatus Similichlamydia laticola
Candidatus Similichlamydia latridicola
Chlamydiales bacterium bin.18
Chlamydiae bacterium O_S4.2_6
Chlamydiia bacterium bin105s
Chlamydiae bacterium O_S4.2_7
Chlamydiae bacterium 1070360-7
Parachlamydiales bacterium Bin_61_76
Chlamydiae bacterium O_S4.2_2
Chlamydiae bacterium K940_chlam_9
Chlamydiae bacterium O_S4.2_123
Chlamydiae bacterium 3300011418-10
Chlamydiae bacterium P_S1_42
Chlamydiae bacterium O_S4.2_81
Chlamydiae bacterium O_S4.2_45
Chlamydiae bacterium O_S4.1_17
Chlamydiae bacterium P_S1_14
Candidatus Clavichlamydia salmonicola
Chlamydia suis
Chlamydia trachomatis
Chlamydia muridarum
Chlamydophila ibidis
Chlamydia sp. 17-3921_L77
Chlamydophila pecorum
Chlamydia sp. 2742-308
Chlamydia sp. H15-1957-10C
Chlamydophila corallus
Chlamydophila pneumoniae
Chlamydophila avium
Chlamydia gallinacea
Chlamydia buteonis
Chlamydophila abortus
Chlamydophila psittaci
Chlamydia felis
Chlamydia sp. S15-834C
Chlamydophila caviae GPIC
Waddliaceae bacterium SP13
Chlamydiae bacterium 1039689-34
Chlamydiae bacterium K940_chlam_3
Chlamydiae bacterium 3300011418-14
Estrella lausannensis
Criblamydia sequanensis
Waddlia chondrophila
Chlamydiae bacterium K940_chlam_7
Chlamydiae bacterium 3300009084-150
Parachlamydiaceae bacterium HS-T3
Chlamydiales bacterium STE3
Parachlamydia sp. BC.030
Parachlamydia acanthamoebae
Parachlamydiaceae bacterium MGR_bin366
Chlamydiae bacterium 3300009787-37
Parachlamydia sp. C2
Candidatus Protochlamydia sp. MGR_bin260
Candidatus Protochlamydia naegleriophila
Candidatus Protochlamydia amoebophila
Protochlamydia sp. ACF82
Parachlamydiaceae bacterium MGR_bin429
Chlamydiae bacterium BWMinControl_Nextera80_MAG2
Parachlamydiaceae bacterium MGR_bin34
Parachlamydiaceae bacterium MGR_bin407
Chlamydiae bacterium BWMinHeated_Nextera45_MAG1
Candidatus Rubidus massiliensis
Chlamydiae bacterium 3300009084-136
Chlamydiae bacterium 3300009083-135
Parachlamydiaceae bacterium SSA5
Chlamydiales bacterium 38-26
Neochlamydia sp. ACF65
Neochlamydia sp. TUME1
Neochlamydia sp. EPS4
Chlamydiae bacterium K940_chlam_8
Chlamydiae bacterium 3300009703-49
Chlamydiae bacterium SM23_39
Candidatus Anoxychlamydiales bacterium K1060_chlam_5
Candidatus Anoxychlamydiales bacterium K1000_chlam_1
Candidatus Anoxychlamydiales bacterium KR126_chlam_6
Candidatus Anoxychlamydiales bacterium K1060_chlam_1
Candidatus Anoxychlamydiales bacterium KR126_chlam_4
Candidatus Anoxychlamydiales bacterium KR126_chlam_5
Candidatus Anoxychlamydiales bacterium K940_chlam_5
Parachlamydiales bacterium B4_bin.14
Chlamydiae bacterium AlinenSedimentsD2_bin-1100
Chlamydiae bacterium 3300012931-71
Parachlamydiales bacterium palsa_1448
Chlamydiae bacterium Kiruna_bin-13304
Chlamydiae bacterium 1021942-14
Chlamydiae bacterium Umea1_bin-1462
Chlamydiae bacterium 3300006177-14
Chlamydiae bacterium RIFCSPLOWO2_02_FULL_45_22
Chlamydiae bacterium CG10_big_fil_rev_8_21_14_0_10_42_34
Chlamydiae bacterium KR126_chlam_1
Chlamydiae bacterium 1096102-30
Chlamydiae bacterium K940_chlam_6
Chlamydiae bacterium KR126_chlam_3
Chlamydiae bacterium SZUA-160
Chlamydiae bacterium RIFCSPLOWO2_02_FULL_49_12
Parachlamydiales bacterium UBA10024
Chlamydiae bacterium 1021942-13
Parachlamydiales bacterium Bin_65_10
Chlamydiae bacterium Ga0074140
Chlamydiae bacterium 1159587-4
Parachlamydiales bacterium palsa_1444
Chlamydiae bacterium 1095360-24
Chlamydiae bacterium 3300009709-46
Chlamydiae bacterium BWOrgHeat_Nextera88_MAG2
Chlamydiae bacterium 1064120-29
Chlamydiae bacterium 3300005529-103
Candidatus Rhabdochlamydia helveticae
Candidatus Rhabdochlamydia porcellionis
Rhabdochlamydia oedothorax
Chlamydiae bacterium O_S4.2_37
Chlamydiae bacterium 3300013133-59
Chlamydiae bacterium 3300013132-109
Chlamydiae bacterium O_S4.2_102
Chlamydiae bacterium 3300006056-139
Chlamydiae bacterium RIFCSPHIGHO2_12_FULL_49_11
Chlamydiae bacterium 3300013126-109
Chlamydiae bacterium O_S4.2_20
Chlamydiae bacterium O_S4.2_27
Chlamydiae bacterium O_S4.1_5
Chlamydiae bacterium P_S1_9
Chlamydiae bacterium 1063924-70
Chlamydiae bacterium O_S4.2_107
Chlamydiae bacterium O_S4.1_1
Chlamydiae bacterium P_S1_54
Chlamydiae bacterium 1045240-7
Simkania negevensis
Candidatus Neptunochlamydia vexilliferae
Sygnamydia xenoturbellae
Syngnamydia medusae
Chlamydiae bacterium 3300006056-100
Chlamydiae bacterium K1060_chlam_2
Chlamydiae bacterium 1049244-12
Chlamydiae bacterium 3300004085-10
Chlamydiae bacterium 1045192-5
0.1

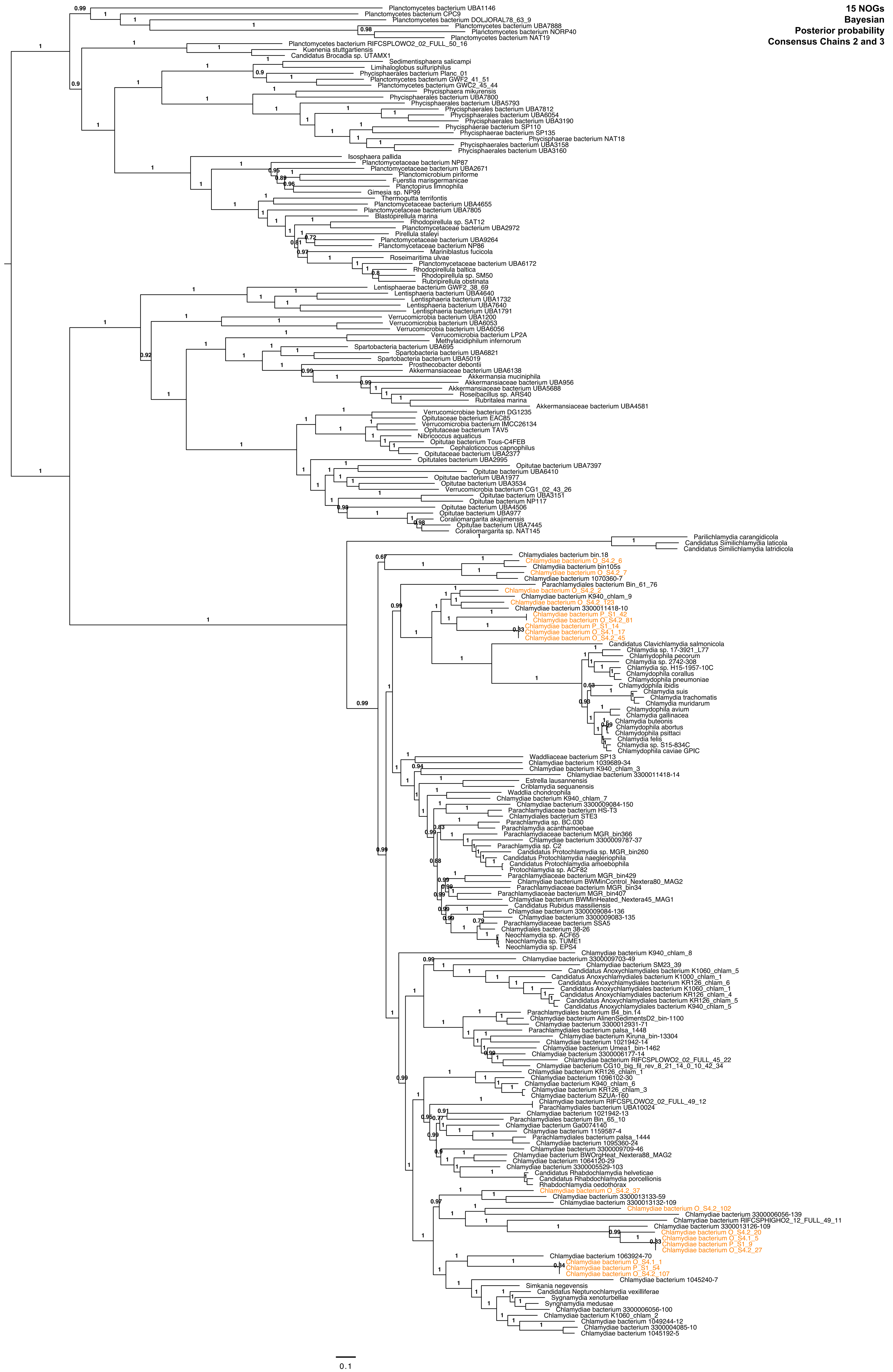
15 NOGs
Bayesian
Posterior probability
Consensus Chains 2 and 3
Planctomycetes bacterium UBA1146
Planctomycetes bacterium CPC9
Planctomycetes bacterium DOLJORAL78_63_9
Planctomycetes bacterium UBA7888
Planctomycetes bacterium NORP40
Planctomycetes bacterium NAT19
Planctomycetes bacterium RIFCSPLOWO2_02_FULL_50_16
Kuenenia stuttgartiensis
Candidatus Brocadia sp. UTAMX1
Sedimentisphaera salicampi
Limihaloglobus sulfuriphilus
Phycisphaerales bacterium Planc_01
Planctomycetes bacterium GWF2_41_51
Planctomycetes bacterium GWC2_45_44
Phycisphaera mikurensis
Phycisphaerales bacterium UBA7800
Phycisphaerales bacterium UBA5793
Phycisphaerales bacterium UBA7812
Phycisphaerales bacterium UBA6054
Phycisphaerales bacterium UBA3190
Phycisphaerae bacterium SP110
Phycisphaerae bacterium SP135
Phycisphaerae bacterium NAT18
Phycisphaerales bacterium UBA3158
Phycisphaerales bacterium UBA3160
Isosphaera pallida
Planctomycetaceae bacterium NP87
Planctomycetaceae bacterium UBA2671
Planctomicrobium piriforme
Fuerstia marisgermanicae
Planctopirus limnophila
Gimesia sp. NP99
Thermogutta terrifontis
Planctomycetaceae bacterium UBA4655
Planctomycetaceae bacterium UBA7805
Blastopirellula marina
Rhodopirellula sp. SAT12
Planctomycetaceae bacterium UBA2972
Pirellula staleyi
Planctomycetaceae bacterium UBA9264
Planctomycetaceae bacterium NP86
Mariniblastus fucicola
Roseimaritima ulvae
Planctomycetaceae bacterium UBA6172
Rhodopirellula baltica
Rhodopirellula sp. SM50
Rubripirellula obstinata
Lentisphaerae bacterium GWF2_38_69
Lentisphaeria bacterium UBA4640
Lentisphaeria bacterium UBA1732
Lentisphaeria bacterium UBA7640
Lentisphaeria bacterium UBA1791
Verrucomicrobia bacterium UBA1200
Verrucomicrobia bacterium UBA6053
Verrucomicrobia bacterium UBA6056
Verrucomicrobia bacterium LP2A
Methylacidiphilum infernorum
Spartobacteria bacterium UBA695
Spartobacteria bacterium UBA6821
Spartobacteria bacterium UBA5019
Prosthecobacter debontii
Akkermansiaceae bacterium UBA6138
Akkermansia muciniphila
Akkermansiaceae bacterium UBA956
Akkermansiaceae bacterium UBA5688
Roseibacillus sp. ARS40
Rubritalea marina
Akkermansiaceae bacterium UBA4581
Verrucomicrobiae bacterium DG1235
Opitutaceae bacterium EAC85
Verrucomicrobia bacterium IMCC26134
Opitutaceae bacterium TAV5
Nibricoccus aquaticus
Opitutae bacterium Tous-C4FEB
Cephaloticoccus capnophilus
Opitutaceae bacterium UBA2377
Opitutales bacterium UBA2995
Opitutae bacterium UBA7397
Opitutae bacterium UBA6410
Opitutae bacterium UBA1977
Opitutae bacterium UBA3534
Verrucomicrobia bacterium CG1_02_43_26
Opitutae bacterium UBA3151
Opitutae bacterium NP117
Opitutae bacterium UBA4506
Opitutae bacterium UBA977
Coraliomargarita akajimensis
Opitutae bacterium UBA7445
Coraliomargarita sp. NAT145
Parilichlamydia carangidicola
Candidatus Similichlamydia laticola
Candidatus Similichlamydia latridicola
Chlamydiales bacterium bin.18
Chlamydiae bacterium O_S4.2_6
Chlamydiia bacterium bin105s
Chlamydiae bacterium O_S4.2_7
Chlamydiae bacterium 1070360-7
Parachlamydiales bacterium Bin_61_76
Chlamydiae bacterium O_S4.2_2
Chlamydiae bacterium K940_chlam_9
Chlamydiae bacterium O_S4.2_T23
Chlamydiae bacterium 3300011418-10
Chlamydiae bacterium P_S1_42
Chlamydiae bacterium O_S4.2_81
Chlamydiae bacterium P_S1_14
Chlamydiae bacterium O_S4.1_17
Chlamydiae bacterium O_S4.2_45
Candidatus Clavichlamydia salmonicola
Chlamydia sp. 17-3921_L77
Chlamydophila pecorum
Chlamydia sp. 2742-308
Chlamydia sp. H15-1957-10C
Chlamydophila corallus
Chlamydophila pneumoniae
Chlamydophila ibidis
Chlamydia suis
Chlamydia trachomatis
Chlamydia muridarum
Chlamydophila avium
Chlamydia gallinacea
Chlamydia buteonis
Chlamydophila abortus
Chlamydophila psittaci
Chlamydia felis
Chlamydia sp. S15-834C
Chlamydophila caviae GPIC
Waddliaceae bacterium SP13
Chlamydiae bacterium 1039689-34
Chlamydiae bacterium K940_chlam_3
Chlamydiae bacterium 3300011418-14
Estrella lausannensis
Criblamydia sequanensis
Waddlia chondrophila
Chlamydiae bacterium K940_chlam_7
Chlamydiae bacterium 3300009084-150
Parachlamydiaceae bacterium HS-T3
Chlamydiales bacterium STE3
Parachlamydia sp. BC.030
Parachlamydia acanthamoebae
Parachlamydiaceae bacterium MGR_bin366
Chlamydiae bacterium 3300009787-37
Parachlamydia sp. C2
Candidatus Protochlamydia sp. MGR_bin260
Candidatus Protochlamydia naegleriophila
Candidatus Protochlamydia amoebophila
Protochlamydia sp. ACF82
Parachlamydiaceae bacterium MGR_bin429
Chlamydiae bacterium BWMinControl_Nextera80_MAG2
Parachlamydiaceae bacterium MGR_bin34
Parachlamydiaceae bacterium MGR_bin407
Chlamydiae bacterium BWMinHeated_Nextera45_MAG1
Candidatus Rubidus massiliensis
Chlamydiae bacterium 3300009084-136
Chlamydiae bacterium 3300009083-135
Parachlamydiaceae bacterium SSA5
Chlamydiales bacterium 38-26
Neochlamydia sp. ACF65
Neochlamydia sp. TUME1
Neochlamydia sp. EPS4
Chlamydiae bacterium K940_chlam_8
Chlamydiae bacterium 3300009703-49
Chlamydiae bacterium SM23_39
Candidatus Anoxychlamydiales bacterium K1060_chlam_5
Candidatus Anoxychlamydiales bacterium K1000_chlam_1
Candidatus Anoxychlamydiales bacterium KR126_chlam_6
Candidatus Anoxychlamydiales bacterium K1060_chlam_1
Candidatus Anoxychlamydiales bacterium KR126_chlam_4
Candidatus Anoxychlamydiales bacterium KR126_chlam_5
Candidatus Anoxychlamydiales bacterium K940_chlam_5
Parachlamydiales bacterium B4_bin.14
Chlamydiae bacterium AlinenSedimentsD2_bin-1100
Chlamydiae bacterium 3300012931-71
Parachlamydiales bacterium palsa_1448
Chlamydiae bacterium Kiruna_bin-13304
Chlamydiae bacterium 1021942-14
Chlamydiae bacterium Umea1_bin-1462
Chlamydiae bacterium 3300006177-14
Chlamydiae bacterium RIFCSPLOWO2_02_FULL_45_22
Chlamydiae bacterium CG10_big_fil_rev_8_21_14_0_10_42_34
Chlamydiae bacterium KR126_chlam_1
Chlamydiae bacterium 1096102-30
Chlamydiae bacterium K940_chlam_6
Chlamydiae bacterium KR126_chlam_3
Chlamydiae bacterium SZUA-160
Chlamydiae bacterium RIFCSPLOWO2_02_FULL_49_12
Parachlamydiales bacterium UBA10024
Chlamydiae bacterium 1021942-13
Parachlamydiales bacterium Bin_65_10
Chlamydiae bacterium Ga0074140
Chlamydiae bacterium 1159587-4
Parachlamydiales bacterium palsa_1444
Chlamydiae bacterium 1095360-24
Chlamydiae bacterium 3300009709-46
Chlamydiae bacterium BWOrgHeat_Nextera88_MAG2
Chlamydiae bacterium 1064120-29
Chlamydiae bacterium 3300005529-103
Candidatus Rhabdochlamydia helveticae
Candidatus Rhabdochlamydia porcellionis
Rhabdochlamydia oedothorax
Chlamydiae bacterium O_S4.2_37
Chlamydiae bacterium 3300013133-59
Chlamydiae bacterium 3300013132-109
Chlamydiae bacterium O_S4.2_102
Chlamydiae bacterium 3300006056-139
Chlamydiae bacterium RIFCSPHIGHO2_12_FULL_49_11
Chlamydiae bacterium 3300013126-109
Chlamydiae bacterium O_S4.2_20
Chlamydiae bacterium O_S4.1_5
Chlamydiae bacterium P_S1_9
Chlamydiae bacterium O_S4.2_27
Chlamydiae bacterium 1063924-70
Chlamydiae bacterium O_S4.1_1
Chlamydiae bacterium P_S1_54
Chlamydiae bacterium O_S4.2_107
Chlamydiae bacterium 1045240-7
Simkania negevensis
Candidatus Neptunochlamydia vexilliferae
Syngnamydia xenoturbellae
Syngnamydia medusae
Chlamydiae bacterium 3300006056-100
Chlamydiae bacterium K1060_chlam_2
Chlamydiae bacterium 1049244-12
Chlamydiae bacterium 3300004085-10
Chlamydiae bacterium 1045192-5
0.1

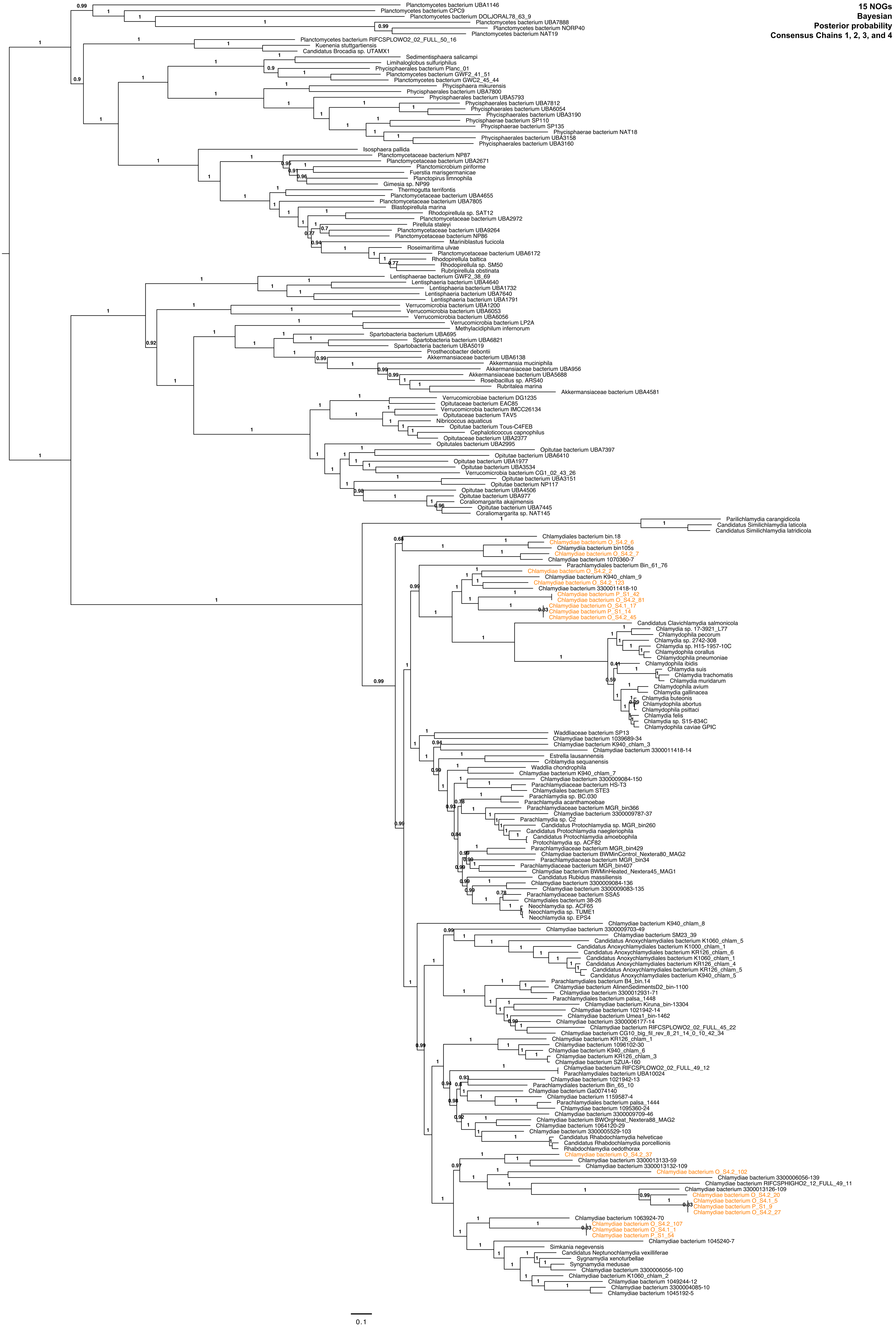
15 NOGs
Bayesian
Posterior probability
Consensus Chains 1, 2, 3, and 4
Planctomycetes bacterium UBA1146
Planctomycetes bacterium CPC9
Planctomycetes bacterium DOLJORAL78_63_9
Planctomycetes bacterium UBA7888
Planctomycetes bacterium NORP40
Planctomycetes bacterium NAT19
Planctomycetes bacterium RIFCSPLOWO2_02_FULL_50_16
Kuenenia stuttgartiensis
Candidatus Brocadia sp. UTAMX1
Sedimentisphaera salicampi
Limihaloglobus sulfuriphilus
Phycisphaerales bacterium Planc_01
Planctomycetes bacterium GWF2_41_51
Planctomycetes bacterium GWC2_45_44
Phycisphaera mikurensis
Phycisphaerales bacterium UBA7800
Phycisphaerales bacterium UBA5793
Phycisphaerales bacterium UBA7812
Phycisphaerales bacterium UBA6054
Phycisphaerales bacterium UBA3190
Phycisphaerae bacterium SP110
Phycisphaerae bacterium SP135
Phycisphaerae bacterium NAT18
Phycisphaerales bacterium UBA3158
Phycisphaerales bacterium UBA3160
Isosphaera pallida
Planctomycetaceae bacterium NP87
Planctomycetaceae bacterium UBA2671
Planctomicrobium piriforme
Fuerstia marisgermanicae
Planctopirus limnophila
Gimesia sp. NP99
Thermogutta terrifontis
Planctomycetaceae bacterium UBA4655
Planctomycetaceae bacterium UBA7805
Blastopirellula marina
Rhodopirellula sp. SAT12
Planctomycetaceae bacterium UBA2972
Pirellula staleyi
Planctomycetaceae bacterium UBA9264
Planctomycetaceae bacterium NP86
Mariniblastus fucicola
Roseimaritima ulvae
Planctomycetaceae bacterium UBA6172
Rhodopirellula baltica
Rhodopirellula sp. SM50
Rubripirellula obstinata
Lentisphaerae bacterium GWF2_38_69
Lentisphaeria bacterium UBA4640
Lentisphaeria bacterium UBA1732
Lentisphaeria bacterium UBA7640
Lentisphaeria bacterium UBA1791
Verrucomicrobia bacterium UBA1200
Verrucomicrobia bacterium UBA6053
Verrucomicrobia bacterium UBA6056
Verrucomicrobia bacterium LP2A
Methylacidiphilum infernorum
Spartobacteria bacterium UBA695
Spartobacteria bacterium UBA6821
Spartobacteria bacterium UBA5019
Prosthecobacter debontii
Akkermansiaceae bacterium UBA6138
Akkermansia muciniphila
Akkermansiaceae bacterium UBA956
Akkermansiaceae bacterium UBA5688
Roseibacillus sp. ARS40
Rubritalea marina
Akkermansiaceae bacterium UBA4581
Verrucomicrobiae bacterium DG1235
Opitutaceae bacterium EAC85
Verrucomicrobia bacterium IMCC26134
Opitutaceae bacterium TAV5
Nibricoccus aquaticus
Opitutae bacterium Tous-C4FEB
Cephaloticoccus capnophilus
Opitutaceae bacterium UBA2377
Opitutales bacterium UBA2995
Opitutae bacterium UBA7397
Opitutae bacterium UBA6410
Opitutae bacterium UBA1977
Opitutae bacterium UBA3534
Verrucomicrobia bacterium CG1_02_43_26
Opitutae bacterium UBA3151
Opitutae bacterium NP117
Opitutae bacterium UBA4506
Opitutae bacterium UBA977
Coraliomargarita akajimensis
Opitutae bacterium UBA7445
Coraliomargarita sp. NAT145
Parilichlamydia carangidicola
Candidatus Similichlamydia laticola
Candidatus Similichlamydia latridicola
Chlamydiales bacterium bin.18
Chlamydiae bacterium O_S4.2_6
Chlamydiia bacterium bin105s
Chlamydiae bacterium O_S4.2_7
Chlamydiae bacterium 1070360-7
Parachlamydiales bacterium Bin_61_76
Chlamydiae bacterium O_S4.2_2
Chlamydiae bacterium K940_chlam_9
Chlamydiae bacterium O_S4.2_123
Chlamydiae bacterium 3300011418-10
Chlamydiae bacterium P_S1_42
Chlamydiae bacterium O_S4.2_81
Chlamydiae bacterium O_S4.1_17
Chlamydiae bacterium P_S1_14
Chlamydiae bacterium O_S4.2_45
Candidatus Clavichlamydia salmonicola
Chlamydia sp. 17-3921_L77
Chlamydophila pecorum
Chlamydia sp. 2742-308
Chlamydia sp. H15-1957-10C
Chlamydophila corallus
Chlamydophila pneumoniae
Chlamydophila ibidis
Chlamydia suis
Chlamydia trachomatis
Chlamydia muridarum
Chlamydophila avium
Chlamydia gallinacea
Chlamydia buteonis
Chlamydophila abortus
Chlamydophila psittaci
Chlamydia felis
Chlamydia sp. S15-834C
Chlamydophila caviae GPIC
Waddliaceae bacterium SP13
Chlamydiae bacterium 1039689-34
Chlamydiae bacterium K940_chlam_3
Chlamydiae bacterium 3300011418-14
Estrella lausannensis
Criblamydia sequanensis
Waddlia chondrophila
Chlamydiae bacterium K940_chlam_7
Chlamydiae bacterium 3300009084-150
Parachlamydiaceae bacterium HS-T3
Chlamydiales bacterium STE3
Parachlamydia sp. BC.030
Parachlamydia acanthamoebae
Parachlamydiaceae bacterium MGR_bin366
Chlamydiae bacterium 3300009787-37
Parachlamydia sp. C2
Candidatus Protochlamydia sp. MGR_bin260
Candidatus Protochlamydia naegleriophila
Candidatus Protochlamydia amoebophila
Protochlamydia sp. ACF82
Parachlamydiaceae bacterium MGR_bin429
Chlamydiae bacterium BWMinControl_Nextera80_MAG2
Parachlamydiaceae bacterium MGR_bin34
Parachlamydiaceae bacterium MGR_bin407
Chlamydiae bacterium BWMinHeated_Nextera45_MAG1
Candidatus Rubidus massiliensis
Chlamydiae bacterium 3300009084-136
Chlamydiae bacterium 3300009083-135
Parachlamydiaceae bacterium SSA5
Chlamydiales bacterium 38-26
Neochlamydia sp. ACF65
Neochlamydia sp. TUME1
Neochlamydia sp. EPS4
Chlamydiae bacterium K940_chlam_8
Chlamydiae bacterium 3300009703-49
Chlamydiae bacterium SM23_39
Candidatus Anoxychlamydiales bacterium K1060_chlam_5
Candidatus Anoxychlamydiales bacterium K1000_chlam_1
Candidatus Anoxychlamydiales bacterium KR126_chlam_6
Candidatus Anoxychlamydiales bacterium K1060_chlam_1
Candidatus Anoxychlamydiales bacterium KR126_chlam_4
Candidatus Anoxychlamydiales bacterium KR126_chlam_5
Candidatus Anoxychlamydiales bacterium K940_chlam_5
Parachlamydiales bacterium B4_bin.14
Chlamydiae bacterium AlinenSedimentsD2_bin-1100
Chlamydiae bacterium 3300012931-71
Parachlamydiales bacterium palsa_1448
Chlamydiae bacterium Kiruna_bin-13304
Chlamydiae bacterium 1021942-14
Chlamydiae bacterium Umea1_bin-1462
Chlamydiae bacterium 3300006177-14
Chlamydiae bacterium RIFCSPLOWO2_02_FULL_45_22
Chlamydiae bacterium CG10_big_fil_rev_8_21_14_0_10_42_34
Chlamydiae bacterium KR126_chlam_1
Chlamydiae bacterium 1096102-30
Chlamydiae bacterium K940_chlam_6
Chlamydiae bacterium KR126_chlam_3
Chlamydiae bacterium SZUA-160
Chlamydiae bacterium RIFCSPLOWO2_02_FULL_49_12
Parachlamydiales bacterium UBA10024
Chlamydiae bacterium 1021942-13
Parachlamydiales bacterium Bin_65_10
Chlamydiae bacterium Ga0074140
Chlamydiae bacterium 1159587-4
Parachlamydiales bacterium palsa_1444
Chlamydiae bacterium 1095360-24
Chlamydiae bacterium 3300009709-46
Chlamydiae bacterium BWOrgHeat_Nextera88_MAG2
Chlamydiae bacterium 1064120-29
Chlamydiae bacterium 3300005529-103
Candidatus Rhabdochlamydia helveticae
Candidatus Rhabdochlamydia porcellionis
Rhabdochlamydia oedothorax
Chlamydiae bacterium O_S4.2_37
Chlamydiae bacterium 3300013133-59
Chlamydiae bacterium 3300013132-109
Chlamydiae bacterium O_S4.2_102
Chlamydiae bacterium 3300006056-139
Chlamydiae bacterium RIFCSPHIGHO2_12_FULL_49_11
Chlamydiae bacterium 3300013126-109
Chlamydiae bacterium O_S4.2_20
Chlamydiae bacterium O_S4.1_5
Chlamydiae bacterium P_S1_9
Chlamydiae bacterium O_S4.2_27
Chlamydiae bacterium 1063924-70
Chlamydiae bacterium O_S4.2_107
Chlamydiae bacterium O_S4.1_1
Chlamydiae bacterium P_S1_54
Chlamydiae bacterium 1045240-7
Simkania negevensis
Candidatus Neptunochlamydia vexilliferae
Sygnamydia xenoturbellae
Syngnamydia medusae
Chlamydiae bacterium 3300006056-100
Chlamydiae bacterium K1060_chlam_2
Chlamydiae bacterium 1049244-12
Chlamydiae bacterium 3300004085-10
Chlamydiae bacterium 1045192-5
0.1

**Phosphoenolpyruvate mutase (PepM) protein phylogeny (PF13714)**

In context of functionally verified proteins (red)

**Maximum-likelihood**

**Ultrafast bootstraps**

**LG+C60+F+R4**

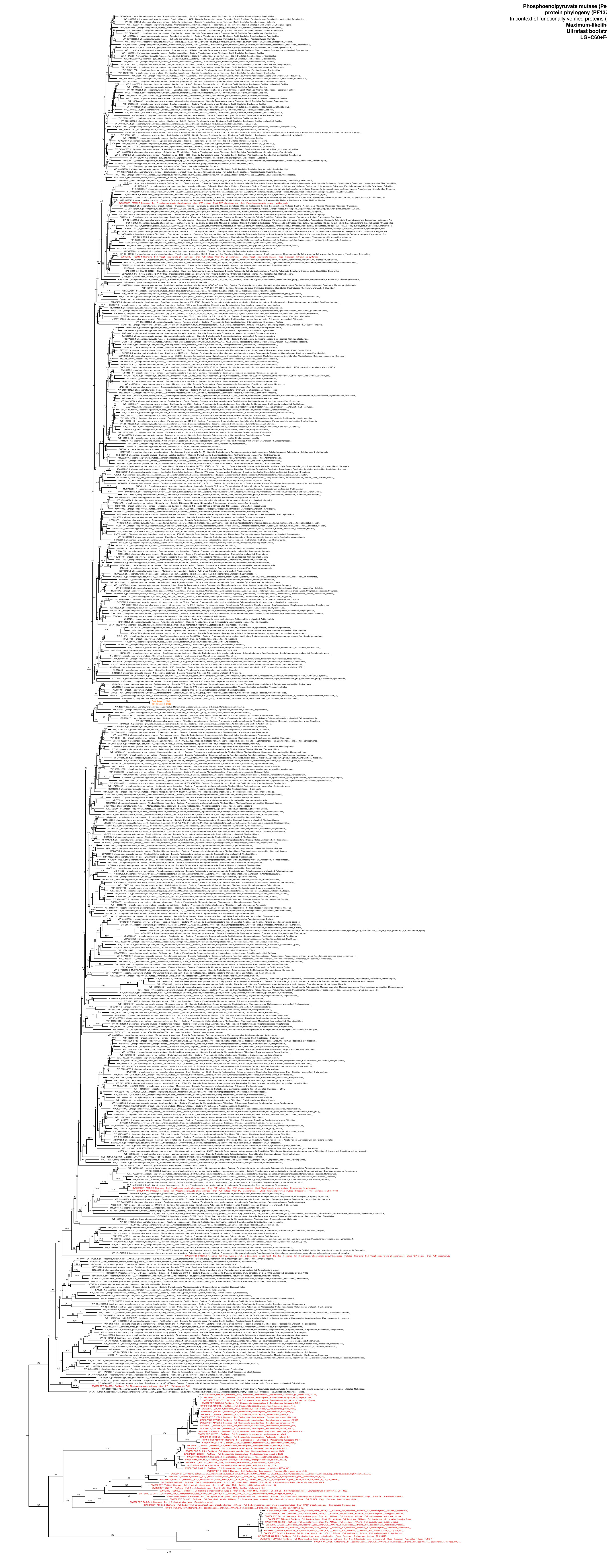

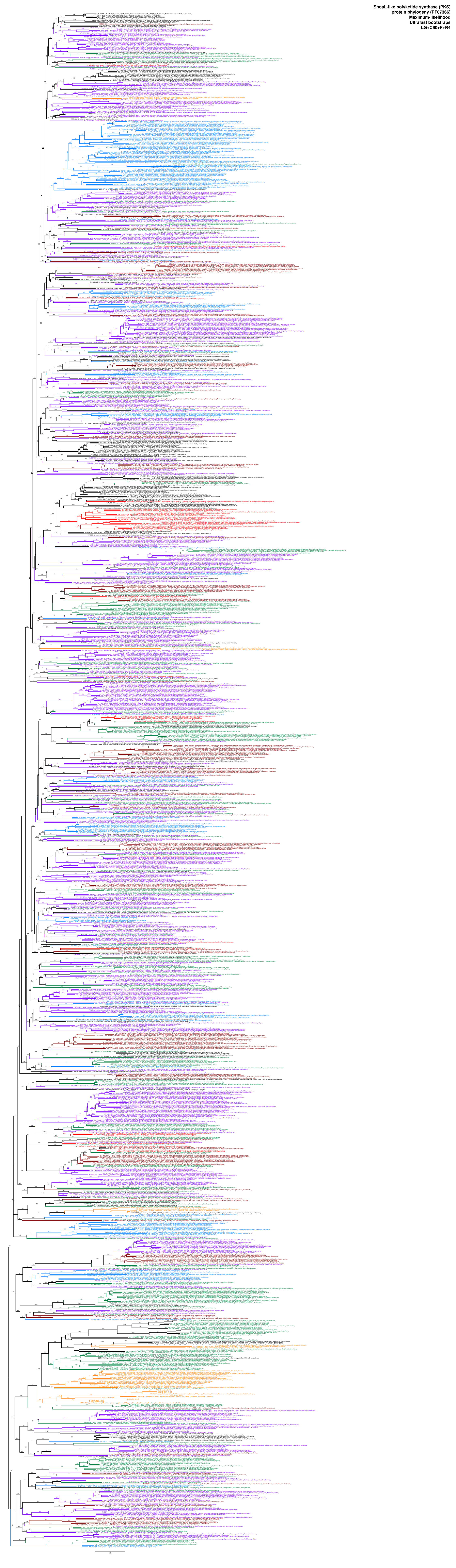
SnoaL-like polyketide synthase (PKS)
protein phylogeny (PF07366)
Maximum-likelihood
Ultrafast bootstraps
LG+C60+F+R4

**Delta24(24(1))-sterol reductase (K00223) and 7-dehydrocholesterol reductase (K00213) with Pfam domain PF01222**

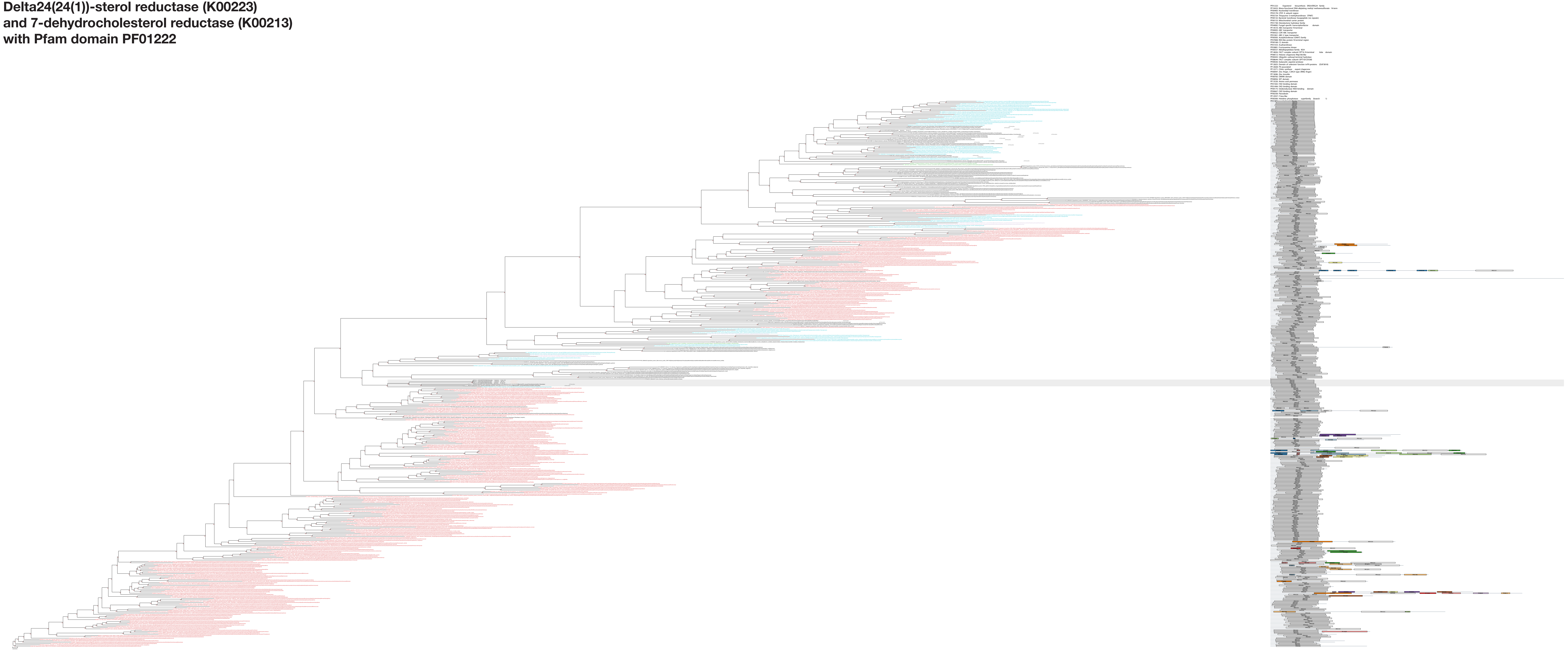

### Inositol oxygenase (K00469) with Pfam domain PF05153

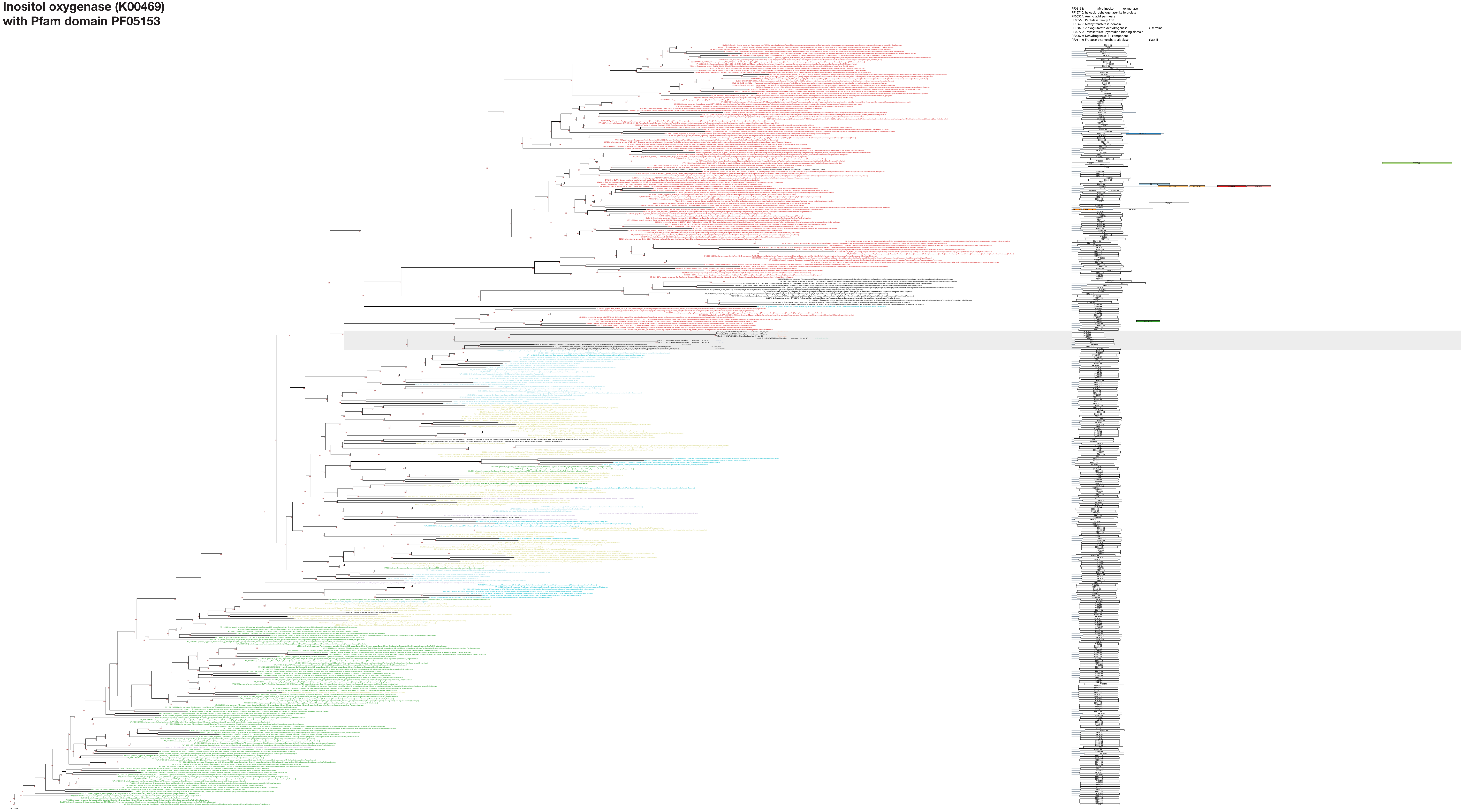

### Carnitine O-acetyltransferase (K00624)

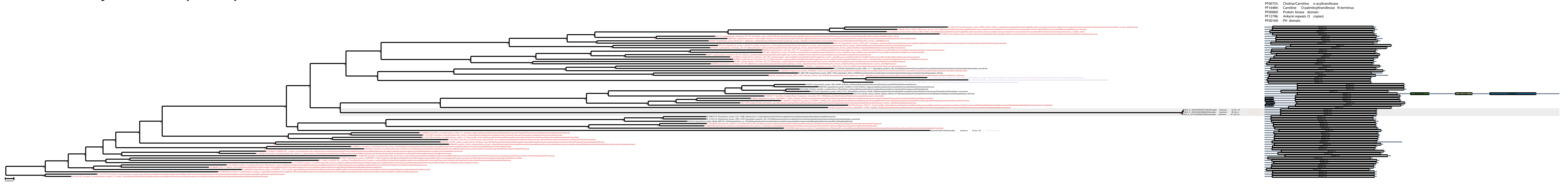

### Delta24-sterol reductase (K09828)

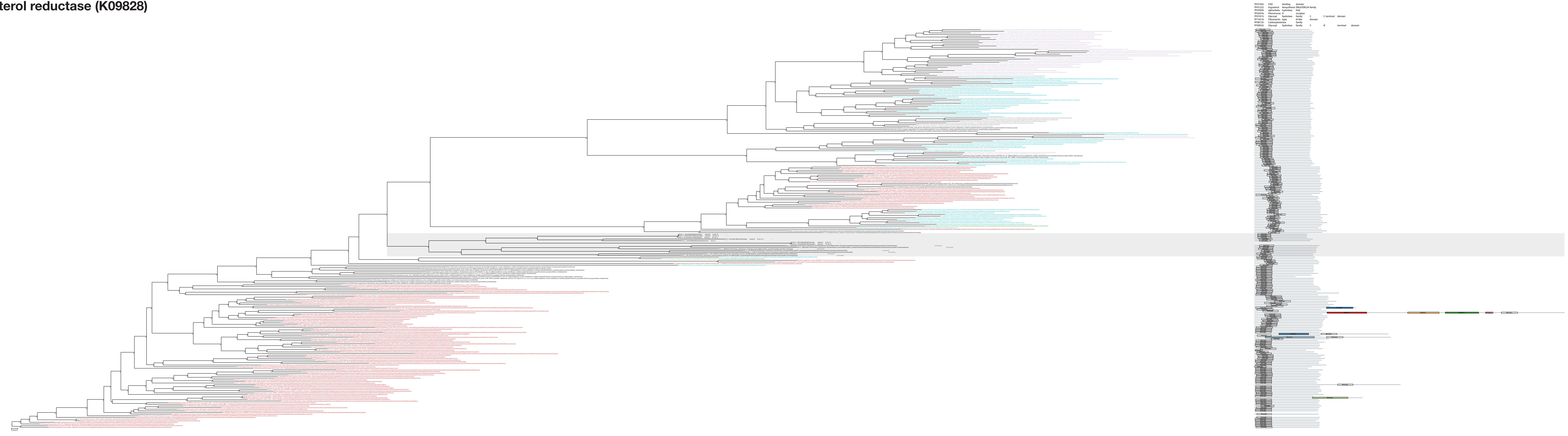

**SSU rRNA gene phylogeny**
sponge-associated chlamydiae
in context of chlamydial diversity
**Maximum-likelihood**
**Ultrafast bootstraps**
**GTR+F+R10**

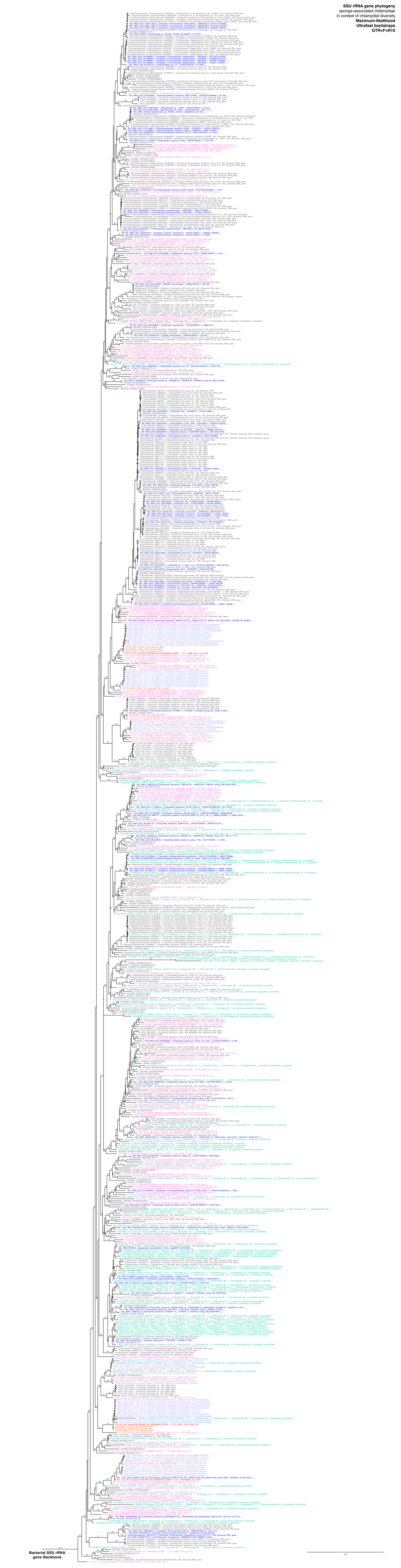
